# Tripartite Inhibition of SRC-WNT-PKC Signalling Consolidates Human Naïve Pluripotency

**DOI:** 10.1101/2020.05.23.112433

**Authors:** Jonathan Bayerl, Muneef Ayyash, Tom Shani, Yair Manor, Ohad Gafni, Yael Kalma, Alejandro Aguilera-Castrejon, Mirie Zerbib, Hadar Amir, Daoud Sheban, Shay Geula, Nofar Mor, Leehee Weinberger, Vladislav Krupalnik, Bernardo Oldak, Nir Livnat, Shadi Tarazi, Shadi Tawil, Lior Lasman, Suhair Hanna, Noa Novershtern, Dalit Ben-Yosef, Sergey Viukov, Jacob H. Hanna

## Abstract

Different conditions have been devised to isolate MEK/ERK signalling independent human naïve pluripotent stem cells (PSCs) that are distinct from conventional primed PSCs and better correspond to pre-implantation developmental stages. While the naïve conditions described thus far endow human PSCs with different extents of naivety features, isolating human pluripotent cells that retain characteristics of ground state pluripotency while maintaining differentiation potential and genetic integrity, remains a major challenge. Here we engineer reporter systems that allow functional screening for conditions that can endow both the molecular and functional features expected from human naive pluripotency. We establish that simultaneous inhibition of SRC-NFκB, WNT/ßCATENIN and PKC signalling pathways is essential for enabling expansion of teratoma competent fully naïve human PSCs in defined or xeno-free conditions. Divergent signalling and transcriptional requirements for maintaining naïve pluripotency were found between mouse and human. Finally, we establish alternative naïve conditions in which MEK/ERK inhibition is substituted with inhibition for NOTCH/RBPj signalling, which allow obtaining alternative human naïve PSCs with diminished risk for loss of imprinting and deleterious global DNA hypomethylation. Our findings set a framework for the signalling foundations of human naïve pluripotency and may advance its utilization in future translational applications.

**Highlights of key findings:**

- Combined inhibition of SRC, WNT and PKC signaling consolidates human naïve pluripotency
- Stable expansion of DNA/RNA methylation-independent and TGF/ACTIVIN-independent human naïve PSCs
- Opposing roles for ACTIVIN and WNT/ßCATENIN signaling on mouse vs. human naive pluripotency
- 2i and MEK/ERKi independent alternative human naïve PSC conditions via inhibiting NOTCH/RBPj signaling

## Introduction

A continuum of pluripotent configurations represent changes occurring during *in vivo* transition of naïve pre-implantation pluripotency toward that of primed post-implantation pluripotent state, can be captured *in vitro* to various extents ((Brons et al., 2007; Hackett and Surani, 2014; Nichols and Smith, 2009; Tesar et al., 2007; Weinberger et al., 2016). Many naïve and primed pluripotency properties can be individually characterized and attributed to pluripotent stem cells (PSCs) expanded in distinct conditions. In mice, defined serum free 2i/LIF conditions (2 inhibitors of MEK and GSK3 supplemented with LIF) have been extensively characterized where many naïve molecular and functional properties are endowed by this combination (Ying et al., 2008a). The latter include global DNA hypomethylation, loss of bivalency over developmental genes (Marks et al., 2012), exclusive nuclear localization of TFE3 transcription factor (Betschinger et al., 2013), tolerance for lack of exogenous L-glutamine (Carey et al., 2015a), tolerance for loss of repressors like DNMT1, METTL3 and DGCR8/DICER (Geula et al., 2015). Mouse ESCs expanded in Fetal Bovine Serum (FBS)/Lif conditions are also considered naïve and possess features such as retention of pre-X inactivation state, ability to tolerate lack of repressors like Mettl3 and Dnmt1 (Geula et al., 2015. However, they do not retain a global hypomethylated epigenome and acquire H3K27me3 over developmental genes (Weinberger et al., 2016a), and thus are considered relatively less naïve than 2i/LIF grown mouse PSCs (Marks et al., 2012). Rodent EpiSCs expanded in Fgf2/Activin A conditions show further consolidation and acquisition of their milieu of primed pluripotency characteristics (Brons et al., 2007; Tesar et al., 2007). EpiSC lines are heterogeneous in their epigenetic and transcriptional patterns (Kojima et al., 2014), and while they are pluripotent and give rise to differentiated cells from all three germ layers, they are epigenetically restricted as evident for example in their reduced ability, after long-term maintenance in FGF2/ACTIVIN A conditions, to differentiate into primordial germ cells (PGCs) (Hayashi et al., 2011) or contribute to chimera formation when injected in the pre-implantation ICM (Wu et al., 2015a).

While conventional human embryonic stem cells (hESCs) and iPSCs (hiPSCs) growth conditions entailed FGF/TGF as typical for murine EpiSC, these two cell types are not identical, and hESC share several molecular features with naïve mESCs including expression of E-CADHERIN (rather than N-CADHERIN) (Weinberger et al., 2016). Further, conventional human ESCs express high levels of PRDM14 and NANOG as murine naïve ESCs, and they are functionally dependent on their expression (Chia et al., 2010). Still however, hESCs retain a variety of epigenetic properties that are consistent with possessing a primed pluripotent state. This includes inability to tolerate MEK/ERK signaling inhibition (Weinberger et al., 2016), predominant utilization of the proximal enhancer element to maintain OCT4 expression, tendency for initiation of X chromosome inactivation (XCI) in most female ESC lines (Mekhoubad et al., 2012), high levels of DNA methylation, prominent deposition of H3K27me3 and bivalency acquisition on lineage commitment regulators (Gafni et al., 2013).

The proof of concept for the metastability between naïve and primed state in rodents (Hanna et al., 2009), have raised the possibility that the human genetic background is more “stringent” in regards to requirement for exogenous factors provided in allowing preservation of ground state-naïve pluripotency in comparison to rodents. Indeed, several groups have previously established condition to derive naïve MEK/ERK signaling independent genetically unmodified human PSCs. For example, NHSM conditions do not require the use of exogenous transgenes or feeder cells, maintain teratoma formation competence and entail the following components: 2iLIF, P38i/JNKi, PKCi, ROCKi, ACTIVIN and FGF2. NHSM conditions endow human PSCs with variety of naïve features including maintaining pluripotency while MEK/ERK signaling is inhibited, predominant TFE3 nuclear localization, resolution of bivalent domains over developmental regulators, *in vitro* reconstitution of human PGCLC and a mild reduction in DNA methylation (Gafni et al., 2013; Irie et al., 2015). The latter effect is profoundly weaker than that seen in mouse pluripotent cells, suggesting sub-optimal human naïve pluripotency growth conditions.

Exciting alternative conditions that generate MEK independent human naïve cells and retain a more compelling milieu of transcriptional markers expressed in the human ICM were developed. Several components found in NHSM conditions (2i, ROCK inhibitor, ACTIVIN, FGF2) were supplemented with BRAF inhibitors, to generate MEF dependent naïve cell lines (termed: 5iLA-, 5iLAF-, 6i/LA- and 4i/LA-MEF) (Theunissen et al., 2014). Globally these conditions generated more pronounced downregulation in DNA methylation and upregulation of naïve pluripotent cell markers. However, the hypomethylation in these conditions was accompanied by immediate and global loss of imprinting over all imprinted loci (Theunissen, 2016) (Bar et al., 2017; Pastor et al., 2016) and obligatory confounding chromosomal abnormalities in nearly 100% of the lines generated within 10 passages only (Liu et al., 2017). Derivation of human naïve ESC in t2iL-Go conditions has been reported with and without the use of exogenous transgenes induction. In both cases, the generated cell lines do not form teratomas *in vivo* and can only differentiate *in vitro* after an extended 3-week transfer to primed-like conditions, thus questioning their pluripotent functionality and stability (Guo et al., 2016; Liu et al., 2017; Takashima et al., 2014). The latter is in striking difference from rodent ground state naïve PSCs, which are fully pluripotent and can initiate differentiation *in vivo* following autonomous induction of the needed priming (capacitation) signals toward differentiation (Ko et al., 2009; Müller et al., 2010; Takahashi et al., 2003; Ying et al., 2008).

Therefore, these prior studies collectively suggest that none of the conditions devised so far are optimal or enable closing the gap between mouse and human naive pluripotency characteristics. In this study, we set out to devise enhanced defined conditions that enable the stabilization of hPSCs with stringent <u>functional and molecular</u> properties, previously attributed to mouse ground state naïve ESCs and to human preimplantation stages, while preserving cell-autonomous teratoma formation competence and decipher defining features of their signaling and regulatory circuitry. We also aimed to establish principles for capturing alternative human naïve PSCs without inhibiting MEK/ERK pathway that tends to compromise DNA imprinting stability in both mouse and human naive PSCs (Choi et al., 2017; Di Stefano et al., 2018; Yagi et al., 2017).

## Results

### Defining human naïve pluripotency conditions that endow tolerance for depletion of RNA/DNA methylation

A major challenge for capturing human naïve PSC conditions has been the lack of stringent functional assays that allow screening for conditions endowing functionally naïve human PSCs (De Los Angeles et al., 2015a). Recent studies have defined the functional ability of mouse naïve 2i/LIF conditions to maintain their pluripotency upon knocking-out epigenetic repressors like Mettl3 and Dnmt1 (m^6^A mRNA and DNA methyltransferase enzymes, respectively), while EpiSCs differentiate and undergo apoptosis upon complete ablation of the same factors (Geula et al., 2015, Liao et al., 2015). Further, DNMT1 ablation leads to cell death and differentiation of human primed ESCs further underscoring this primed aspect of their molecular identity (Liao et al., 2015). As such, we hypothesized that the latter functional attributes might be utilized to screen and define conditions endowing this stringent naïve pluripotency feature (i.e. maintenance of pluripotency in the absence of defined epigenetic repressors). We engineered human WIBR3 hESC lines with conditional inducible ablation of the expression of METTL3 (**Fig. 1a**). This was obtained via introducing an exogenous METTL3 transgene under the regulation of Tet-OFF promoter (Liao et al., 2015), followed by CRISPR/Cas9 mediated ablation of both endogenous human METTL3 alleles (**Fig. 1a**). Two resultant clones were validated for METTL3 expression only from the exogenous allele, which can be shut off by addition of DOX to the media (Tet-OFF-METTL3 lines) (**Fig. 1b**).

**Figure 1.**
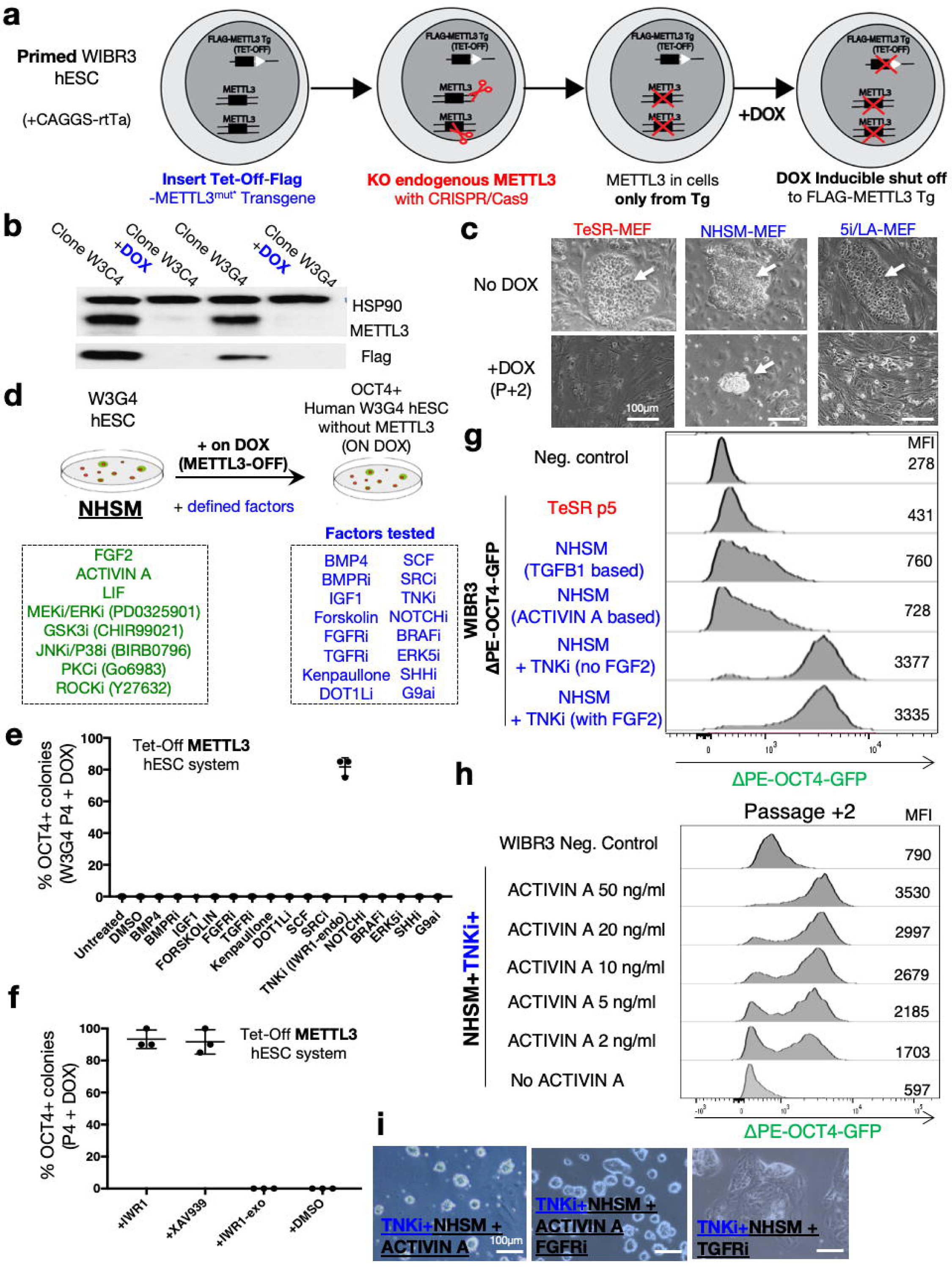
Reporter systems for functional screening for enhanced human naive pluripotency conditions. **a.** Strategy for generating human ESCs with TET-OFF regulated expression of METTL3 m^6^A methyltransferase enzyme. **b**. Western blot analysis for correctly engineered human ESCs with TET-OFF METTL3 regulation, before and after DOX treatment. **c**. Representative images for human METTL3 TET-OFF engineered cells in different conditions with or without DOX addition. White arrow indicated viable PSC at P+2. Previously described primed and naïve conditions do not maintain their pluripotency and viability when DOX is added (METTL3 is depleted). **d.** Scheme depicting strategy for conducting a screen for identifying a small molecule or cytokine additive to previously described human naïve NHSM conditions that allow maintaining pluripotency in TET-OFF METTL3 human stem cells after adding DOX. **e**. W3G4 cells were maintained in the presence of DOX for up to 4 passages in different conditions and stained for OCT4 to quantify percentage of cells that retained their pluripotency. Graph shows that supplementing NHSM conditions with an inhibitor for Tankyrase (TNKi) allows maintaining pluripotency in majority of cells expanded (>75% positive OCT4 staining). **f.** OCT4+ pluripotency maintenance in NHSM conditions supplemented with various TNK inhibitors and DOX to repress METTL3 expression. **g.** Quantification of ΔPE-OCT4-GFP knock in naïve pluripotency reporter, in variety of primed (red) and naïve conditions (blue). Mean fluorescence intensity values (MFI) are indicated. **h.** Quantification of ΔPE-OCT4-GFP knock in naïve pluripotency reporter in variety of conditions with various concentrations of ACTIVIN A recombinant cytokine. Figure shows that in NHSM+TNKi conditions the naivety of human ESCs is still dependent on ACTIVIN A supplementation. **i**. Representative phase contrast images in human ESCs expanded in NHSM+TNKi conditions showing their maintenance of pluripotent domed-like morphology even in the presence of FGFRi. However, upon blocking of TGF/ACTIVIIN A signaling (with A83-01 designated as TGFRi), the cells in NHSM+TNKi lose their pluripotent dome-like shaped morphology and differentiate.

Primed Tet-OFF-METTL3 hESCs expanded in primed TeSR or KSR/FGF2 conditions could not be sustained in the presence of DOX more than four passages (both on MEF or Geltrex coated dishes) and resulted in massive cell death and differentiation (**Fig. 1c**), analogous to the result previously reported for mouse EpiSCs (Geula et al., 2015). The latter is also consistent with lack of reports showing generation of complete KO METTL3 cells in human primed cells (Bertero et al., 2018). In the presence of MEFs, Tet-OFF-METTL3 could be poorly maintained in previously described human naive NHSM conditions and with a very slow proliferation rate, but not in 4iLA-MEF, 5iLAF-MEF, 5iLA-MEF, 6iLA-MEF, TESR/3iL-MEF previously described naïve conditions (**Fig. 1c, S1a**). However, in the absence of MEFs, also NHSM conditions could not support maintenance of pluripotency when METTL3 was completely ablated, suggesting that NHSM conditions can possibly be enhanced to endow the cells with such ability (**Fig. S1a**). We thus set out to test candidate molecules that may enrich NHSM condition and allow to maintain Tet-OFF-METTL3 on Geltrex coated plates in the presence of DOX.

We tested whether supplementing NHSM conditions with individual small molecules that have been previously implicated in boosting mouse or human naïve pluripotency formation or iPSC derivation efficiency might achieve this goal (**Fig. 1d)**. Remarkably, supplementing NHSM conditions with the Tankyrase inhibitor named IWR1, but not any of the other 15 compounds tested, enabled expanding Tet-OFF-METTL3 on DOX with great homogeneity (**Fig. 1e)**. IWR1 is a WNT inhibitor (WNTi) small molecule that stabilizes AXIN protein in the cytoplasm by inhibiting Tankyrase enzyme (abbreviated herein as TNK inhibitor – TNKi)(Huang et al., 2009). It was included serendipitously among our compounds of interest since when combined with GSK3 inhibitors on mouse and human primed cells, this effect lead to enriching βCatenin only in the cytoplasm and such primed cells could be expanded without exogenous FGF2 (Kim et al., 2013a). Using an independent specific TNKi, XAV939, yielded a similar effect, while using exo-IWR1 an inactive modified version of IWR1 failed to do so, supporting specific inhibition of Tankyrase as the target yielding stability of these pluripotent cells (**Fig. 1f)**. Notably, a number of recent publications have included TNKi among other ingredients for both naïve, extended potential or primed hPSCs, however none of these different conditions that employ TNKi (Bredenkamp et al., 2019a; Smith et al., 2012; Yang et al., 2017a, 2017b; Zimmerlin et al., 2016a) allow to maintain OCT4+ in the absence of METTL3 or DNMT1, likely due to the absence of other key ingredients synergistically used in NHSM conditions but not in these previously described conditions (i.e. combination of ACTIVIN, PKCi, P38i/JNKi together with TNKi combination) (**Fig. S1b**).

We used previously described WIBR3 hESC line carrying knock-in ΔPE-OCT4-GFP reporter (Theunissen et al., 2014) (**Fig. S1c**) and found that supplementation of TNKi to NHSM conditions yielded a dramatic increase in GFP signal when compared to primed, NHSM or 4i-LA conditions (**Fig. 1g, Fig. S1c)**. Consistent with studies conducted in mice (Kim et al., 2013a), including TNKi rendered exogenous supplementation of FGF2 dispensable even in feeder free conditions (**Fig. 1g**). Further, as XAV939 inhibits WNT signaling, we validated that including GSK3 inhibitor is dispensable and, in fact, compromises the intensity of ΔPE-OCT4-GFP signal (**Fig. S1d**). JNK/P38 inhibition boosted naïve pluripotency marker expression and cell viability, therefore were maintained in the media used herein (**Fig. S1e**). Importantly, after optimizing NHSM conditions, we retried to substitute TNKi with other components included in the screen, to exclude the possibility that the latter optimizations may facilitate a different screening result. However, none of them allowed expanding METTL3 depleted cells *in vitro* as seen with supplementing TNKi (including VPA, BRAFi, Forskolin, Kenopaullone, SHHi, DOT1Li, LSD1i, TGFRi, ERK5i) (**Fig. S1f**).

### Defining human naïve pluripotency conditions independent of TGF/ACTIVIN signaling

Following the latter modifications human ESCs maintained uniformly high ΔPE-OCT4-GFP levels only in the presence of exogenous ACTIVIN A, and consistently differentiated when TGFR inhibitor was provided (**Fig. 1h-i, Fig. S1g).** We next tested whether a second targeted screening strategy would allow us to identify a small molecule whose supplementation will render human PSCs that are independent of exogenous ACTIVIN/TGF supplementation. It should be noted that none of the previously described human naïve conditions have been able to maintain teratoma competent pluripotent cells that can be maintained long term and validated for their naïve identity after prolonged specific inhibition of ACTIVIN/NODAL signaling (Liu et al., 2017). To do this, the latter TNKi supplemented NHSM conditions were used in the absence of ACTIVIN A, and candidate molecules were added to screen for allowing expanding OCT4+ PSCs independent of METTL3 expression (on DOX) and the absence of exogenous ACTIVIN A (**Fig. 2a**). While in most conditions OCT4+ cell fraction rapidly deteriorated, we noted that a validated SRC inhibitor (SRCi = CGP77675) dramatically maintained the stability of dome like cells that were uniformly OCT4+ (**Fig. 2b**). The latter led us to assemble a defined FGF/TGF/ACTIVIN/MEF independent and serum free conditions which we term Human <u>E</u>nhanced Naïve Stem cell Medium – **HENSM** (**Fig. 2c-d**), which we systematically characterize and validate herein. We also note that supplementation of SRCi in ACTIVIN A containing conditions, although was not essential to maintain ΔPE-OCT4-GFP+ when ACTIVIN A was provided, supported consistency in domed like morphology among naïve cells **(Fig. S1h)** (we term this conditions **HENSM-ACT** and is highlighted herein when used).

**Figure 2.**
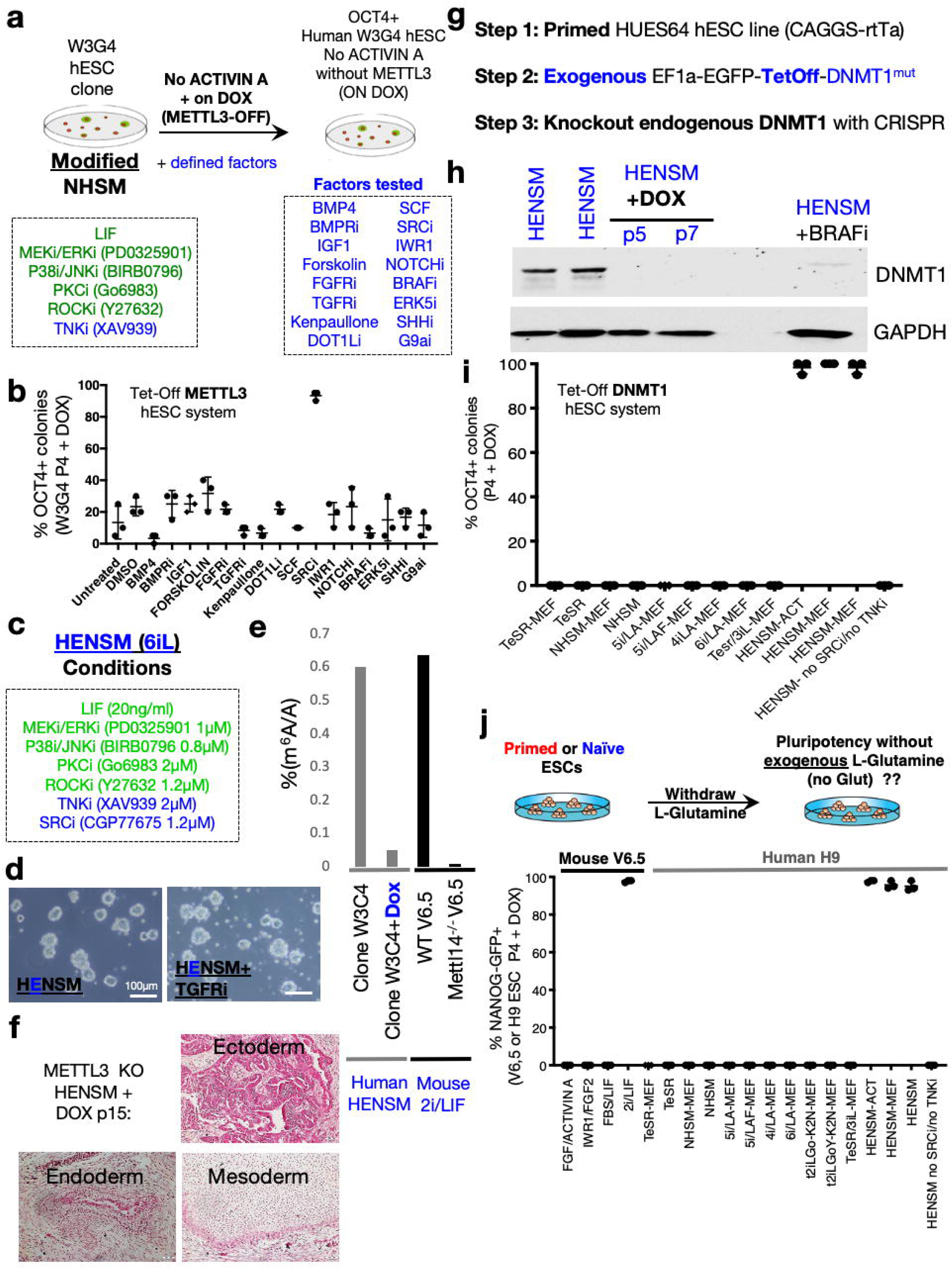
Defining enhanced human naive conditions compatible with blocking TGF/ACTIVIN signalling. **a.** Scheme depicting strategy for conducting a screen for identifying a small molecule or cytokine additive to optimized NHSM conditions after addition of TNKi, allow to maintain pluripotency in TET-OFF METTL3 human stem cells (clone W3G4) and without supplementing exogenous TGF or ACTIVIN. **b**. OCT4+ pluripotency maintenance in optimized naive conditions without TGF/ACTIVIN, indicated that supplementing SRCi CGP77675 allows maintaining OCT4+ cells in optimized conditions and without METTL3 expression (n=3 per condition). **c**. Summary of small molecules and their concentrations used in the optimized HENSM conditions used herein. **d**. Representative phase contrast images showing naïve domed-like morphology of human ESCs expanded in HENSM conditions, that is maintained even when TGFRi small molecule A83-01 is supplemented. **e**. Mass spectrometry-based quantification of m^6^A on isolated mRNA from the indicated cell lines and conditions. Depletion of m^6^A in human cells was validated in HENSM + DOX conditions. **f**. Mature teratoma obtained following injection of METTL3 TET-OFF human ESCs expanded for 15 passages (P15) in HENSM + DOX conditions. Please note that no *in vitro* priming or media other than HENSM+DOX was used before the cells were injected into the mice to test for teratoma formation. **g**. Strategy for generating human ESCs with TET-OFF regulated expression of DNA methyltransferase enzyme, DNMT1. **h**. Western blot analysis for DNMT1 expression in HENSM conditions supplemented with either DOX of BRAF inhibitor (SB590885 – 0.25µM). Please note that DOX ablates DNMT1 expression and that BRAFi depletes DNMT1 expression to much lower levels than seen in HENSM conditions (without DOX), yet still they retain residual DNMT1 expression that is necessary for their survival and viability when BRAFi is added to HENSM conditions. **i.** DNMT1 TET-OFF ESC clone was maintained in the presence of DOX for up to 4 passages in different conditions and stained for OCT4 to quantify percentage of cells that retained their pluripotency. Graph shows that only HENSM and HENSM-ACT conditions (with and without MEFs) maintain robust expansion of dome-like undifferentiated human PSCs in vitro. Omitting TNKi and SRCi (in other words WNTi and SRCi) form HENSM leads to loss of ability to maintain DNMT1 depleted human naïve PSCs, as evident from loss of OCT4+ cells. **j**. Mouse or human ESCs carrying NANOG-GFP pluripotency reporter were expanded in the indicated naïve and primed conditions in the absence of exogenous L-Glutamine supplementation for 4 passages. Percentage of pluripotent cells was quantified based on GFP expression levels. Graph shows that only HENSM and HENSM-ACT conditions (with and without irradiated feeder cells – MEFs) maintain expansion and stability of human NANOG+ pluripotent cells when exogenous L-Glutamine is omitted, and that this is similar to 2iL conditions on mouse naïve ESCs. Omitting TNKi and SRCi form HENSM leads to loss of ability to maintain human naïve PSCs without exogenous L-Glutamine supplementation, as evident from loss of NANOG-GFP+ cells.

Both with and without METTL3 depletion in HENSM conditions, WIBR3 cells maintained their typical domed like morphology and uniformly expressed pluripotency markers including KLF17 that is specific to the human naïve state (**Fig. S2a-b**). Measurement of m^6^A on mRNA showed over 90% depletion of total levels after DOX addition (**Fig. 2e**), comparable to those seen upon knockout of the Mettl3/14 in mouse naïve cells. ESCs maintained in the absence or presence of DOX for over 30 passages remained pluripotent and, as it is typically standardized for murine ESCs and iPSCs, the human cells were capable of generating mature teratomas *in vivo* without the need to passage them first for a period of time in primed conditions or capacitation via unique conditions (Rostovskaya et al., 2019) prior to their injection in immune-compromised mice (**Fig. 2f**). The latter validate maintenance of naïve pluripotency in human PSCs expanded in HENSM conditions, including when METTL3 protein and m^6^A levels on mRNA were ablated.

To extend the previous findings to another repressor machinery, ΔPE-OCT4-GFP-WIBR3 reporter human ESCs were targeted by CRISPR/Cas9 to generate DGCR8 null cells (**Fig. S2d-e**). While conducting such targeting on primed cells did not yield any null clones (**Fig. S2d**), consistent with inability to sustain mouse primed and human ESCs without DGCR8 due to its essentiality in this state (Geula et al., 2015), DGCR8 null cells could be obtained when the targeted cells were expanded in HENSM conditions (**Fig. S2e-f)**. DGCR8 KO cells expanded in HENSM retained human naïve pluripotency specific traits despite lacking microRNA expression equivalent to results obtained in mouse naïve PSCs (**Fig. S2g**). To test which of the naïve conditions enable expanding human naïve PSCs in the absence of DNMT1 cells, a similar approach to that applied for making Tet-OFF-METTL3 herein, has been recently used to generate TET-OFF DNMT1 in HUES64 ESC line (**Fig. 2g**)(Liao et al., 2015). Cells expanded in previously described naïve conditions including NHSM, 5i/LA. t2iL-GO (or PXGL) and TESR/3iL-MEF could not be maintained in the presence of DOX for more than 3 passages (**Fig. 2h-i, S2h**). HENSM and HENSM-ACT conditions allowed stable and unlimited expansion of DNMT1 depleted human naïve ESCs both in feeder and feeder free conditions (**Fig. 2i, S2h)**. Whole Genome Bisulfite Sequencing (WGBS) confirmed global loss of DNA methylation in naïve DNMT1 depleted cells expanded in HENSM conditions (**Fig. S2i**) that also maintained expression of all pluripotency markers (**Fig. S2j**). Removal either of ERKi, PKCi, SRCi or TNKi in HENSM conditions resulted in loss of pluripotency when DNMT1 or METTL3 were depleted (**Fig. S2k**). Collectively these results demonstrate that HENSM conditions closely mimic stringent characteristics of mouse naïve ground state ESCs and, for the first time, enable generation of human PSCs ablated for epigenetic repressors (both in feeder and feeder free conditions) and that are independent from ACTIVIN/TGF signaling.

### Tolerance for absence of exogenous L-Glutamine in HENSM conditions

Murine naïve ESCs retain bivalent metabolic capability utilizing both oxidative phosphorylation (OXPHOS) and glycolytic metabolism, while upon priming they become dependent only on glycolytic metabolism. As shown previously, naïve hPSCs in NHSM, 5i-LA and transgene induced t2iL-Go conditions increase OXPHOS activity leading to retention of a bivalent metabolic profile (Sperber et al., 2015). HENSM conditions were similarly tested herein. Measured basal oxygen consumption rate (OCR) was substantially higher in HENSM conditions than in primed PSC (**Fig. S2l**). Higher electron transport chain activity in HENSM was evidenced by a greater OCR increase in response to the mitochondrial uncoupler FCCP (**Fig. S2l**). Both NHSM, HENSM and transgene dependent t2iL-Go reset conditions, but not primed or 5iLA conditions, PSCs maintained pluripotency when glucose uptake was blocked with 1mM 2-Deoxy-D-Glucose (DG), similar to what was observed with mouse naïve PSCs in 2i/LIF (**Fig. S2m)**.

More stringently, a newly identified metabolic feature in naive ESCs in 2i or 2i/LIF is that they can endogenously synthesize glutamine at sufficient levels to maintain adequate alpha-ketogluterate (αKG) levels (Carey et al., 2015a). While they benefit from exogenous L-Glutamine supplementation, it is not essential for their stability or pluripotency as they can metabolically synthesize it endogenously as part of their altered metabolic configuration (Carey et al., 2015a). FBS/LIF naïve murine ESCs or primed EpiSCs cannot be maintained in the absence of exogenous L-Glutamine for an extended period of time (Carey et al., 2015a). To compare the latter observation and apply them on distinct human pluripotent states, WIBR3-OCT4-GFP knock-in ESC line, ΔPE-WIBR3-OCT4-GFP knock-in ESC line and H9-NANOG-GFP ESC lines were then tested for ability to maintain pluripotency in the presence and absence of exogenously added L-Glutamine (**Fig. 2j)**. Importantly, we failed to maintain primed PSCs or other previously described naïve PSCs in the absence of L-Glutamine (in NHSM, 4i/LA-MEF, 5i/LAF-MEF, 6iLA-MEF, TESR/3iL-MEF, t2iLGo even in feeder cells presence for more than 10 days (**Fig. 2j**). However, ΔPE-WIBR3-OCT4-GFP expanded in HENSM was not compromised when L-glutamine was not included in HENSM conditions (**Fig. S3a**) and GFP signal was positive for H9-NANOG-GFP ESC both in the presence and absence of L-Glutamine (both on feeder and feeder free conditions) (**Fig. S3a**). Cells expressed general and naïve specific pluripotency markers in HENSM with and without exogenous L-Glutamine and generated differentiated teratomas without the need for *in vitro* passaging in primed conditions before subcutaneous *in vivo* injection (**Fig. S3b-c**). Other previously published combination that entailed TNKi among other ingredients for both naïve and primed PSCs (Bredenkamp et al., 2019; Wu et al., 2015; Yang et al., 2017, Zimmerlin et al., 2016), failed to preserve pluripotency in the absence of exogenous L-Glutamine as seen in HENSM based conditions described herein (**Fig. 2j**). The latter is due to the absence of other key ingredients synergistically used in NHSM conditions contributing to naive pluripotency that are used herein but not in these previously described conditions (i.e. the unique combination of SRCi, PKCi together with TNKi combination in HENSM) (**Fig. 2j**). Collectively, these results validate that HENSM conditions can maintain naïve pluripotency characteristics and endow the cells with ability to be expanded in the absence of exogenous L-Glutamine as previously described for murine 2i/LIF naïve PSCs (Carey et al., 2015a).

### Transcriptional characterization of human PSCs in HENSM conditions

We next aimed to revert previously established primed PSCs lines and to derive new lines directly in HENSM-ACT and HENSM conditions from the ICM of human blastocysts. Human blastocysts were plated on mouse embryonic fibroblast (MEF) coated plates and medium successfully generated domed cell outgrowths following 6-8 days of plating. ICM derived outgrowths were then trypsinized and passaged. Subsequently, we were able to establish and characterize 5 newly derived stem cell lines termed LIS36, LIS42 and LIS46 in HENSM-ACT and LIS41 and LIS49 ESCs in HENSM conditions (**Fig. S4a)**. Multiple conventional (hereafter will be named “primed”) hESC lines (WIBR1, WIBR2, WIBR3, HUES64, H1, H9) were plated on Geltrex coated dishes in HENSM medium **(Fig. S4b)**. Within 4-8 days of applying this protocol, dome-shaped colonies with packed round cell morphology, typical of mESCs, could be readily isolated and further expanded **(Fig. S4b)**. Adult human dermal fibroblast cells or peripheral blood cells were reprogrammed to iPSCs in HENSM conditions following either lentiviral transduction with DOX inducible OKSM factors (BF1 hiPSC) or by non-integrating sendai viruses (JH1and MECP5 hiPSC) (**Fig. S4c).** All polyclonal and subcloned hESC and iPSC lines expanded in HENSM conditions were uniformly positive for pluripotent markers alkaline phosphatase (AP), OCT4, NANOG, SSEA4, TRA1-60 and TRA1-81 (representative images in **Fig. S5)** and robustly formed mature teratomas *in vivo* without the need for short- or long-term exposure to primed growth conditions prior to their injection into host mice, and as typically observed with rodent ground state naïve PSCs (**Fig. S6a**). Naïve lines were passaged with TryplE every 3-5 days and had single cell cloning efficiency up to 40%, while primed cell single cell cloning increased only up to 10% even when ROCK inhibition was used. Human naïve pluripotent lines maintained normal karyotype after extended passaging in HENSM-ACT or HENSM in most lines of tested (**Fig. S7a)**. In some cultures (<10%) and after over 30 passages, minor aneuploidy cells were observed however we did not observe a recurrent abnormality between any of these lines as determined by G-banding of metaphase chromosomes (**Fig. S7a)**. The latter results were corroborated by performing e-karyotyping that is based on RNA-seq expression of HENSM derived lines (**Fig. S6b**), thus indicating that epigenetic resetting in HENSM does not cause obligatory chromosomal abnormalities nor select for pre-existing abnormal variants, as has been seeing for other naïve conditions like 5iLA-MEF or t2iL-Go conditions that induce 96-100% chromosomal abnormality already by passage 10 (Liu et al., 2017).

We compared global gene expression patterns between naïve and primed hESCs and hiPSCs, many of which were genetically matched. Unbiased clustering of genome-wide expression profiles demonstrated that naïve hESC and hiPSCs possess a distinct gene expression pattern and clustered separately from conventional/primed hESCs and hiPSCs **(Fig. 3a-b)**. Transcripts associated with naïve pluripotency were significantly upregulated in naïve cells (Blakeley et al., 2015a). The later included NANOG, TFCP2L1, KLF17, KLF4, STELLA (DPPA3), DPPA5, DNMT3L, KHDC1L and ARGFX shown in two independent datasets **(Fig. S8a and Table S1-S2)**. RT-PCR analysis validated the dramatic upregulation in naïve PSCs expanded both in HENSM and HENSM-ACT conditions **(Fig. 3c)**. When including previously and independently published naïve datasets generated in 5iLA-MEF, 4i-LAF and t2i-LIF-GO-NK2, we note that cells generated in HENSM and HENSM-ACT conditions clustered with all the latter naïve conditions and not with primed samples (**Fig. 3a-b, Fig. S8b-f**). The same transcriptional qualities were detected in human PSCs expanded in HENSM Xeno-Free (HENSM-XF conditions) on BioLaminin511 (**Fig. 3a).** FACS analysis confirmed upregulation of previously identified human naïve pluripotency markers Gb3/CD77 and IL6ST/CD130 in HENSM conditions, and primed pluripotency marker CD24 was downregulated in HENSM conditions (**Fig. S8g**) (Collier et al., 2017). SUSD2 surface marker was also expressed on all naïve cells lines tested, however was meaningfully expressed on nearly 50% human primed cell lines tested **(Fig. S9e)**, which provides a cautionary note regarding recent claims (Bredenkamp et al., 2019) that it is an exclusive and decisive marker to exclude the human primed state *in vitro.* Importantly, naïve pluripotent cells had profoundly down regulated transcripts associated with lineage commitment genes including T, ZIC2 and VIM1 that are expressed in primed hESCs (**Fig. S8a, S9a-b)**. By comparing the transcriptome to that described during early human pre-implantation development, cells expanded in HENSM conditions, but not primed cells, showed specific enrichment to profile of late morula-early blastocysts specific signature (**Fig. 3d**). We additionally introduced STELLA-CFP knock-in allele via CRISPR/Cas9 (**Fig. S9c-d**), to monitor pluripotency maintenance in the different tested conditions, and STELLA-CFP was upregulated in both HENSM and 5iLA conditions (**Fig. S9d**). Similar to its upregulation and importance in maintaining human naïve pluripotency in 5i/LA conditions (Pastor et al., 2018), TFAP2C KO lines (**Fig. S10a-d)** showed that it is essential for deriving and maintaining human naïve PSCs in both HENSM and HENSM-ACT conditions (**Fig. S10e**). ESRRB is not detectable in human genetically unmodified cells expanded in HENSM conditions (**Fig. S10f-j)**, consistent with its negative expression in the human ICM (Blakeley et al., 2015b; Petropoulos et al., 2016). The latter results confirm that HENSM conditions attain consensus transcriptional features observed in other previously published naïve hPSCs studies or *in vivo* measured human embryo transcriptional data, while uniquely retaining the critical ability to generate teratomas after being directly injected as naïve cells in immune-compromised mice.

**Figure 3.**
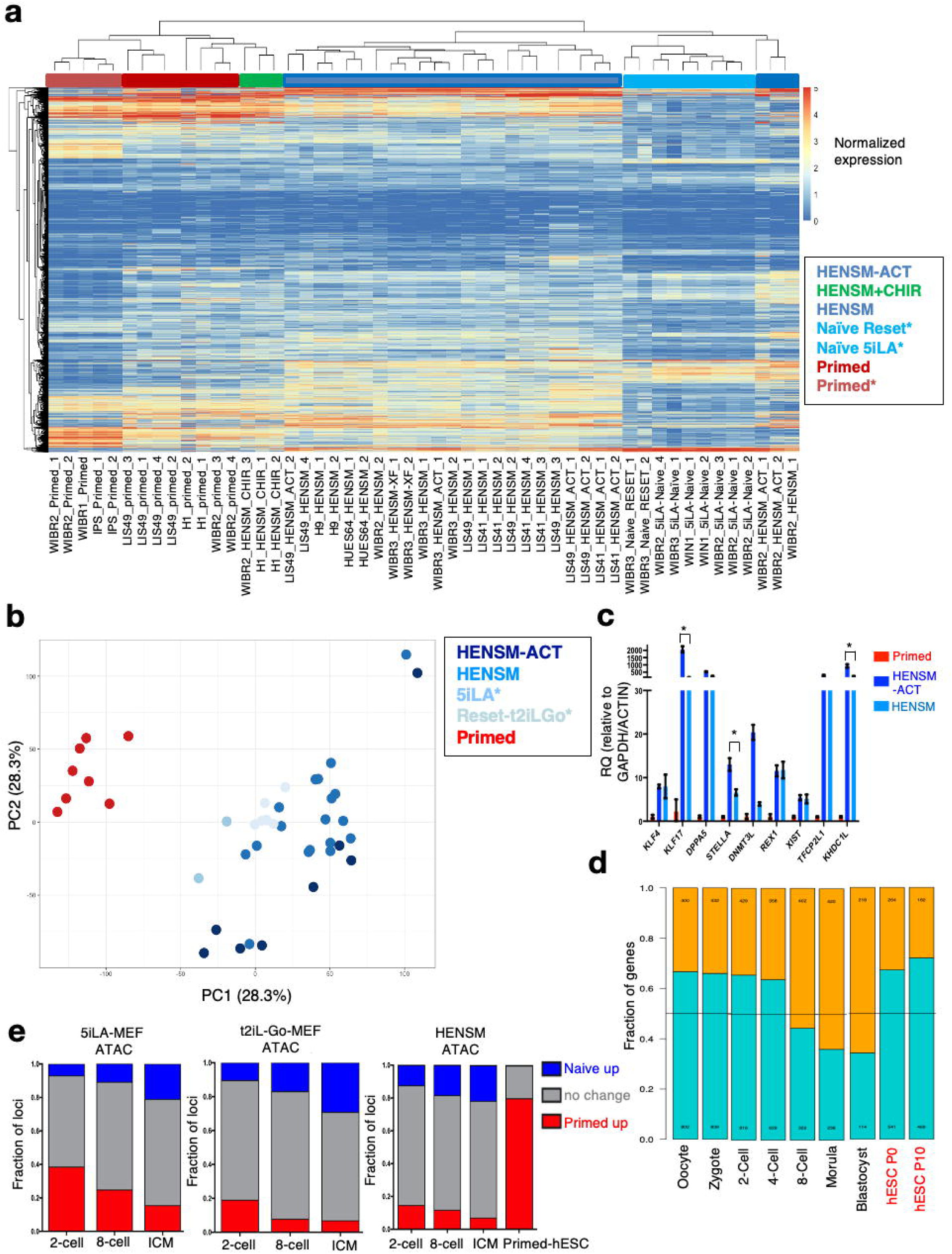
HENSM endows human PSCS with canonical naïve-like transcriptional features. **a.** Unbiased hierarchical clustering was performed on RNA-seq measurement obtained from different human ESC and iPSCs expanded in HENSM, HENSM-ACT naïve and TeSR primed conditions. The data was also clustered with previous independently generate RNA-seq on human PSCs expanded in 5iLA conditions (Naïve 5iLA*, Theunissen et al. Cell Stem Cell 2016), Reset conditions (Naïve Reset*, Takashima et al. Cell 2014 – composed of NANOG-KLF2 transgenes and 2iLGo) or Primed conditions (Primed*). Figure shows that HENSM and HENSM-ACT conditions (Dark blue) resemble 5iLA and Reset conditions (light blue) and cluster separately from primed cells (Red and orange). **b**. PCA analysis of samples represented in **a**, showing that HENSM and HENSM-ACT conditions (Dark blue) resemble 5iLA and Reset conditions (light blue) and cluster separately from primed cells (Red). **c**. RT-PCR analysis for naïve pluripotency markers. Values were normalized to ACTIN and GAPDH. Primed expression levels were set as 1. *t-test *p* Value < 0.01. Please note that HENSM-ACT show higher expression of naïve pluripotency markers than HENSM, consistent with the notion that ACTIVIN A supports naïve pluripotency in humans. **d.** Correspondence between gene expression in naïve/primed ESCs and single-cell human embryonic stages (Yan et al. 2013). For every stage of human embryonic development, a statistical test was performed to find the genes that have a different expression level compared to other stages. The proportions of developmental stage-specific genes that are upregulated (p <0.05, 2-fold change) in naïve or primed cells are indicated in orange and blue, respectively. **e.** Uniquely accessible regions were recognized in 2/8-cell and ICM of human early development. The percentage of up-regulated regions (FC>2) in naïve (Blue) or primed (Red) are indicated. Analysis was done for 3 distinct naïve systems: HENSM, t2iL/Gö and 5iL/A (previously mapped in Wu et al). Accessible regions in naïve tend to have a higher overlap with ICM regions than with 2- or 8-cell regions.

We profiled and compared Transposable Element (TE)-derived transcripts in conventional and naïve human PSCS expanded in HENSM conditions (Theunissen et al., 2016). The top 5,000 TEs with largest SD separated naïve and primed samples both in hierarchical clustering (**Fig. S11a, Table S3**) and in PCA based analysis (**Fig. S11b**). Members of the SINE-VTR-Alu (SVA) family of TEs and HERVK-associated LTR were transcribed almost exclusively in HENSM conditions similar to previously obtained in 5i/LA and transgene dependent t2iLGo conditions (**Fig. S12)** (Theunissen et al., 2016). We used TE profiling to measure the degree to which HENSM and primed conditions resemble pluripotent cells in early human embryos *in vivo*. HENSM naïve, but not primed cells, demonstrated the most significant overlap with the human morula and epiblast stages when looking at TEs (**Fig. S11c**), as was similarly shown for coding genes (**Fig. 3d**).

### Epigenetic characterization of human PSCs in HENSM conditions

To further delve into the patterns which distinguish the naïve condition from its primed counterpart in hESC and converge with its *in vivo* equivalents, we explored change in chromatin accessibility of each condition by ATAC-seq. By comparing unique chromatin accessibility loci found in different pre-implantation stages (i.e. 2-cell, 8-cell and ICM) we compared how much of these loci show an increase or decrease in chromatin accessibility of naïve compared to primed hESCs. Like in datasets generated in 5iLA-MEF and t2iL-Go-MEF, HENSM hESCs chromatin accessibility corresponds more to the ICM than earlier stages **(Fig. 3e).** These results support the endowment of late pre-implantation like transposon expression and chromatin accessibility profile in PSCs expanded in HENSM conditions *in vitro*.

We next moved to test whether HENSM conditions endow human naive PSCs with a pre-X chromosome configuration. We used previously generated primed human WIBR2 hESCs carrying knock-in MECP2-tdTomato and MECP2-GFP alleles (**Fig. 4a**) (Theunissen et al., 2016). Correctly targeted clone #29-9 expresses only the red allele, however upon transferring the cells into HENSM conditions >99% of cells expressed both fluorescent markers consistent with reactivation of both x chromosome alleles. Transferring the cells into primed media allowed inactivation of X chromosome as evident by obtaining GFP-/tdTomato+ pattern >95% of the reprimed cells (**Fig. 4a**). We carried out FISH analysis for ATRX in female cells as this locus is expressed from one copy even in human primed WIBR3 hESCs that have undergone erosion of X chromosome (Xe). Indeed, two ATRX foci could be uniformly found in naïve, but not primed, human female PSCs supporting reactivation of X chromosome (**Fig. 4b**). SNP based analysis of X chromosome allele expression as detected in RNA-seq datasets showed biallelic expression of X chromosome encoded genes in HENSM but not primed conditions (**Fig. 4c**), consistent with functional reactivation of X chromosome in female naïve PSCS induced in HENSM (Bar et al., 2019).

**Figure 4.**
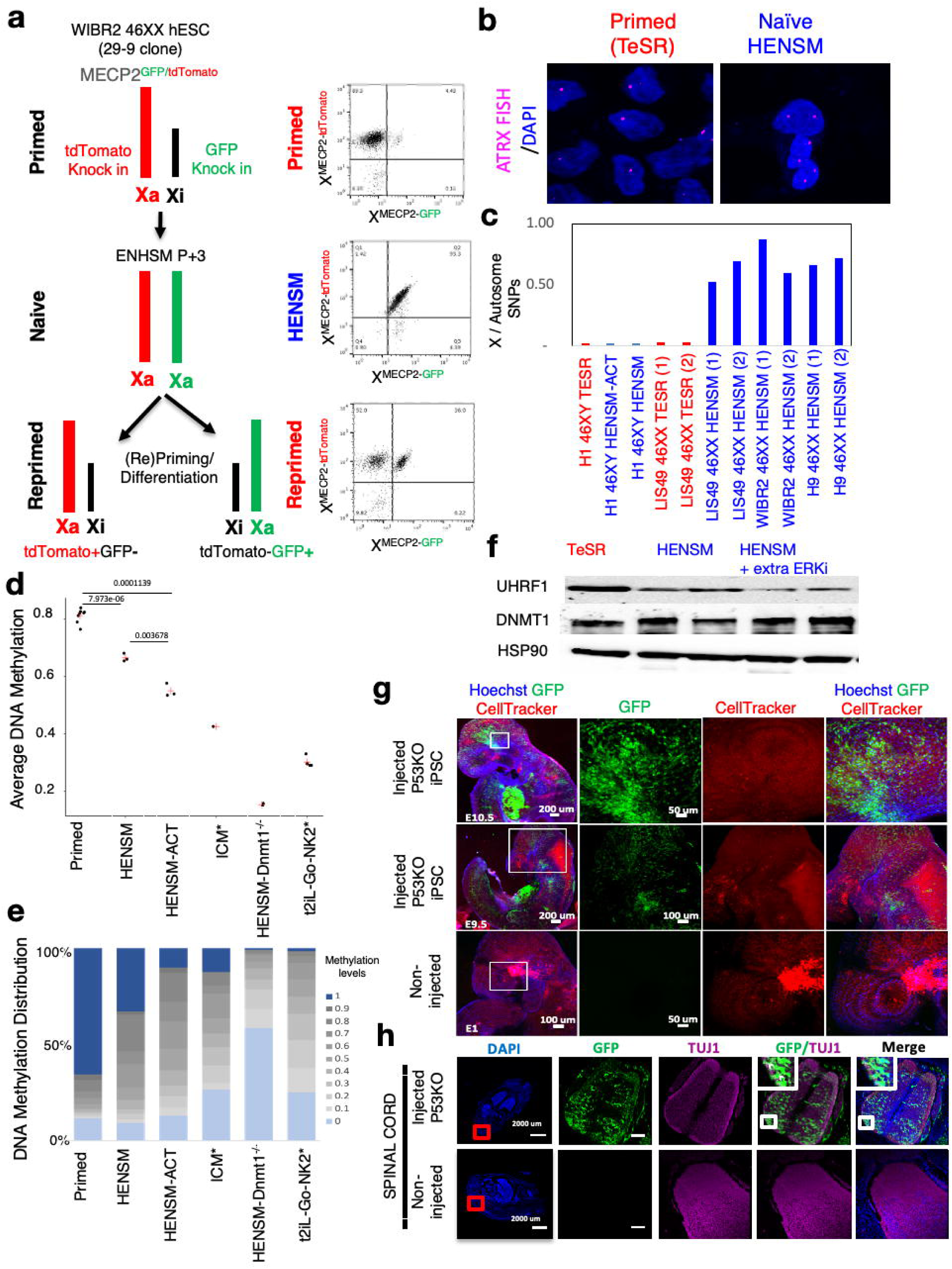
Chromosome X reactivation status and DNA methylation profile in HENSM based conditions. **a**. Schematic and FACS results following using WIBR2 (female 46XX) 29-9 hESC line that carries GFP and tdTomato on each of the X chromosomes in the MECP2 locus. Parental 29-9 clone has the X chromosome carrying mCherry allele in the active state, and thus is positive only for tdTomato and negative for GFP in the primed state. Upon transfer to HENSM conditions, all cells turn on both X chromosomes and thus become double positive for both fluorescent markers (GFP and tdTomato). After transfer into primed conditions (i.e. repriming), cells start to inactivate the X chromosome again. **b**. RNA-FISH analysis for ATRX transcription in primed and HENSM WIBR3 cells. Note that ATRX is active on both X chromosomes only in HENSM conditions. **c**. X:Autosome allelic ratios calculated for primed and naïve PSCs (value of each single RNA-seq sample). **d.** Average methylation as calculated from primed samples, and naïve samples that were maintained in various HENSM conditions, along with previously published Reset-naïve samples (Takashima et al), and human ICM samples (Guo et al, Nature 2014). DNMT1^-/-^ (from TET-OFF lines) samples were used as negative control for methylation. **e.** Global methylation histogram measured on the same samples as in (d). Dark blue - percentage of highly methylated CpGs (>0.9 methylation level), light blue – percentage of lowly methylated CpGs (<0.1 methylation level). **f**. Western blot analysis for DNA methylation regulators, DNMT1 and UHRF1 enzymes. DNMT1 protein levels are maintained in all conditions. UHRF1 is partially depleted in HENSM conditions, and this decrease is more enhanced when ERKi concentration is increased. **g.** Representative images of whole-mount *in-toto* imaged mouse embryos, after microinjection of the indicated human naïve iPSCs, are shown in comparison to non-injected wild-type embryos. White squares in tiles outline zoomed-in regions in subsequent panels. GFP staining was used to trace hiPSC-derived progeny and CellTracker and Hoechst as counter staining. **h.** Representative images of IHC for TUJ1 and GFP of injected (upper panels) and non-injected E15.5 mouse embryos (lower panels) for spinal cord region are shown. GFP served as human cell tracer and TUJ1 as neural marker. GFP, TUJ1, overlap as well as merged constitute zoomed-in regions of tissues depicted in red squares in the tiles. White arrowheads in insets depict co-localization of GFP and TUJ1. TUJ1, neuron-specific class III beta-tubulin; GFP, green fluorescent protein; p53, tumor protein p53; iPSC, induced pluripotent stem cell. Tile scale bar 200 and 100µm. Zoomed-in scale bar 50µm.

We sampled human naïve PSCs in HENSM and HENSM-ACT for DNA methylation status by Whole genome Bisulfite Sequencing (WGBS). Lines tested displayed profound downregulation of global methylation levels **(Fig. 4d-e)** from 80% in primed hPSCs to 50% in HENSM-ACT and to 65% in HENSM expanded human hPSCs. It has been previously shown in mouse ground state naive conditions that rapid loss of global DNA methylation level result from partial decrease in UHRF1 protein expression levels while maintaining DNMT1 levels (von Meyenn et al., 2016). Indeed, in human cells expanded in HENSM, DNMT1 methyltransferase protein expression is maintained, while UHRF1 protein is partially (50-60%) depleted **(Fig. 4f)** which may underlie the global downregulation in DNA methylation in HENSM conditions. It is important to note that supplementing HENSM with BRAF inhibitor used in 5i/LAF conditions leads to dramatic downregulation also in DNMT1 **(Fig. 2h)**. This pattern is also observed in 5i/LAF condition which might explain the immediate and global loss of imprinting (loss of imprinting of >95% of imprinted loci within 5 passages). As DNMT1 levels are maintained in HENSM conditions and UHRF1 protein downregulation was partial, we did not observe immediate global loss of all imprinted genes in HENSM or HENSM-ACT conditions **(Fig. S13)** as has been previously seen in 5iLA or 2iL-Go based conditions (Pastor et al., 2016b; Theunissen, 2016b). However, loss of imprinting in HENSM occurred sporadically only at some individual loci, at lower frequencies and only after prolonged cell passaging (>15-20 passages, **Fig. S13**), which constitutes an improvement over previously reported hypomethylated human naïve PSCs and comparable frequency to those observed in human primed cells (Bar et al., 2017). Collectively, the above findings indicate that HENSM and HENSM-ACT conditions consolidate human naïve pluripotency identity and endow them with nearly all known naïve pluripotency and differentiation features that have been attributed to human ICM *in vivo*, previously derived human naïve cells and murine naïve ground state cells.

### Differentiation competence of naive hiPSC following interspecies chimaerism in mice

Given that the naïve hPSCs described above maintained genetic integrity and were capable of forming teratoma without the need for exogenous priming or capacitation prior to injection, we next moved to evaluate whether human naïve iPSCs can integrate and differentiate following microinjection into host mouse blastocyst (Gafni et al., 2013b; Wu and Izpisúa Belmonte, 2016). Microinjection of human GFP labeled naïve iPSCs (maintained for at least 3 passages in HENSM based conditions) into mouse E2.5 morulas showed robust integration in mouse *in vitro* developed ICMs at E3.5. Blastocysts obtained after micro-injection with human naïve iPSCs were implanted in pseudo-pregnant female mice and their survival and integration was assayed throughout next 14 days using various imaging techniques. HENSM-derived hiPSCs are able to colonize mouse embryos up to E17.5 at various anatomic regions of different embryonic germ layers (**Fig. S14-S15**). Specific marker staining for human nuclei excluded any contamination by mouse PSC lines (**Fig. S14**).

Our results indicate that the contribution levels of human GFP+ cells in mouse embryos, although meaningful and reproducible, are relatively low (∼3% of mice are chimeric with naïve hiPSC derived cells). Thus, we sought to develop technical platforms that can enhance such integration and also allow us to better visualize integrating human cells in the developing mouse embryo. Recently, mouse ESC depleted for p53 outcompete host pluripotent cells after blastocyst microinjection by cell competition phenomenon that has been recently documented in mice (Dejosez et al., 2013), and can generate very high contribution chimeras by blocking their tendency for apoptosis. Based on the latter, we tested whether manipulating these pathways might similarly endow injected naïve hiPSCs with competitive advantage and increase human cell chimerism in developing mouse embryos. GFP hiPSC lines were targeted with CRISPR/Cas9 to generate P53^-/-^ cell lines (**Fig. S16, S17**). Remarkably, after microinjection into mouse morulas or aggregation with 2-cell embryos (Fig. S18), a 4-fold increase in the number of chimeric animals was obtained (**Fig. 4g, Fig. S18-S25**) and a dramatic increase in the extent of chimerism per embryo (up to 20% cells per embryo **Fig. S25c**). Furthermore, differentiation was validated in chimaeric embryos for major developmental markers covering all three embryonic germ layers overlapping extensively with human-derived GFP signal (**Fig. 4h, Fig. S18-S25**). Taken together, these results substantiate generation of advanced human-mouse interspecies chimaera with various engraftment in many defined lineages up to E17.5 from HENSM based conditions.

### WNT/ß-CATENIN and SRC/NFkB signaling are major priming pathways compromising human naïve pluripotency

The results above indicate that functional naïve pluripotency in HENSM composition not only relies on inhibition of ERK and PKC pathways, but also on inhibition of TNK and SRC. Depletion of any of the inhibitors for these 4 pathways compromised naïve pluripotency as exemplified by X chromosome inactivation in reporter female hESCs (**Fig. 5a**). Combined depletion of TNKi and SRCi pushed human PSCs toward complete loss of pluripotency within a number of passages (**Fig. 5b, S26a**), underlining the conclusions that ERKi together with simultaneous tripartite inhibition of PKCi-SRCi-TNKi are essential for functional defined conditions for human naïve pluripotency. We next aimed to define the signaling pathway downstream of Tankyrase inhibition that facilitate human naïve PSCs stabilization. WNT signaling is a well-established promoter of rodent naive pluripotency where it leads to ß-CATENIN (ßCAT) nuclear localization which in turn neutralizes the repression by TCF3 effector and derepresses naïve pluripotency targets (ten Berge et al., 2011; Wray et al., 2011). Although some studies have indicated that titrating WNT stimulation level is needed when working with human naïve cells via using lower doses of GSK3 inhibitor or by adding Tankyrase inhibitor together with GSK3i (leading to high cytoplasmic ß-CAT)(Theunissen et al., 2014; Zimmerlin et al., 2016), they concluded that WNT stimulation is helpful for promoting human naïve pluripotency exactly as seen in mouse PSCs. Other studies, indicated that WNT inhibition is beneficial for initial reversion of primed to naïve pluripotency, however its continued application compromised human naive pluripotency and that 2i with titrated WNT stimulation is the optimal pattern for maintaining human naive PSCs (Bredenkamp et al., 2019a, Bredenkamp et al., 2019b), and that WNT inhibition, rather than stimulation, was needed for priming and capacitation of human naïve pluripotency (Bredenkamp et al., 2019a).

**Figure 5.**
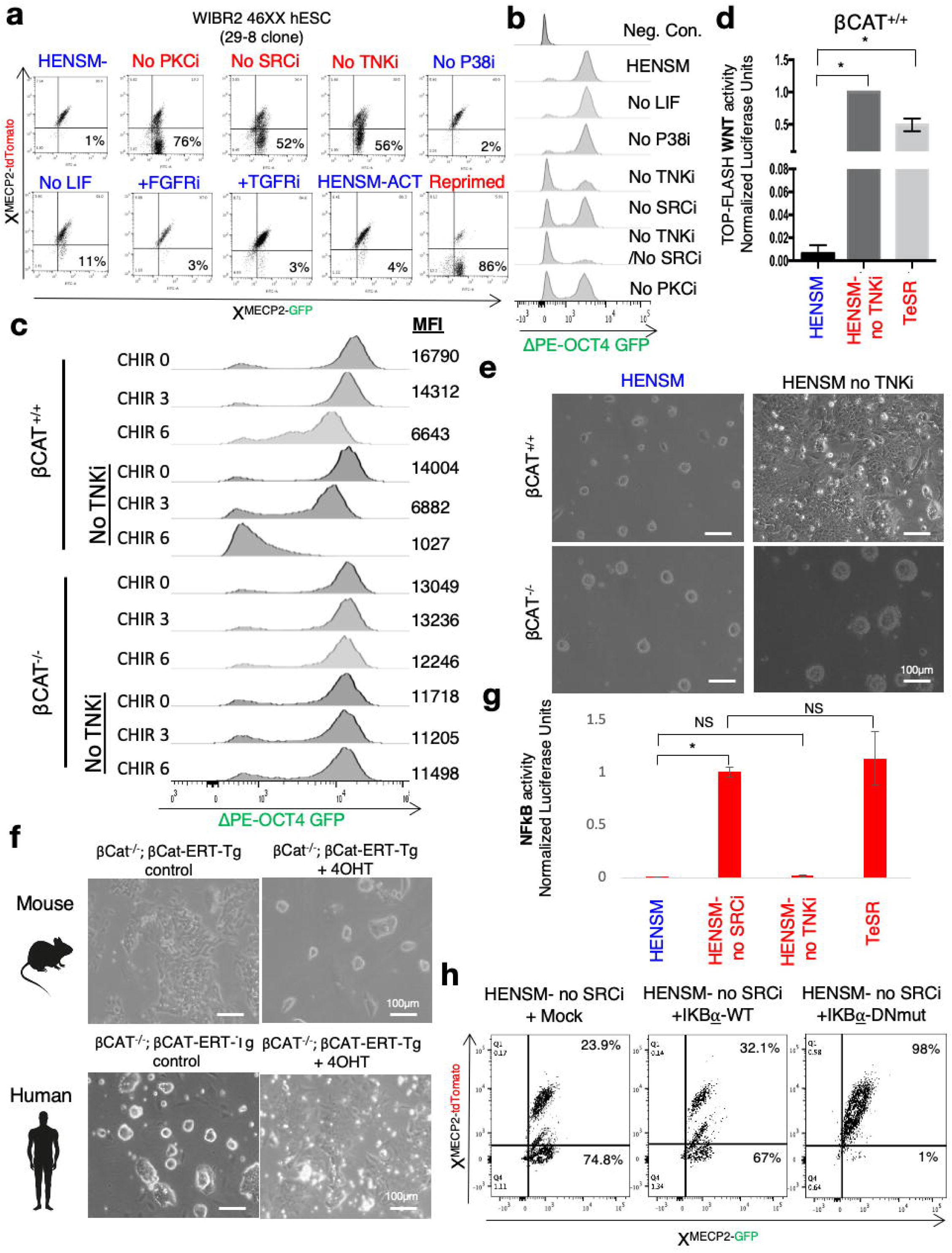
Signalling foundations for human naïve pluripotency. **a**. FACS results following using WIBR2 (female 46XX) 29-8 hESC line that carries GFP and tdTomato on each of the X chromosomes in the MECP2 locus. Parental 29-8 clone has the X chromosome carrying GFP allele in the active state, and thus is positive only for GFP and negative for tdTomato in the primed state. Upon transfer to HENSM conditions, all cells turn on both X chromosomes and thus become double positive for both fluorescent markers (GFP and mCherry). Indicated components from HENSM conditions were individually depleted or added, and cells were subjected to FACS analysis after 10 days. Withdrawal of PKCi, SRCi or WNTi compromises XaXa state and cells start inactivating one of the X chromosomes, indicating loss of naïve state identity. Figure shows that upon LIF omission, cells do not become primed and cells maintain XaXa state. **b**. Quantification of ΔPE-OCT4-GFP knock in naïve pluripotency reporter in variety of conditions after 2 passage transfer from HENSM starting conditions. *t-test *p* Value < 0.01. **c.** Quantification of ΔPE-OCT4-GFP knock in naïve pluripotency reporter in βCAT^+/+^ and isogenic βCAT^-/-^ cells in variety of conditions after 3 passage transfer from HENSM starting conditions. **d.** Luciferase assay for measuring WNT activity via luciferase reporter assay. Values were normalized to HENSM no TNKI conditions (defined as 1; n=3 per condition). **e.** Morphological changes in βCAT^+/+^ and isogenic βCAT^-/-^ cells after 2 passages from TNK withdrawal. Knock in naïve pluripotency reporter in variety of conditions after 2 passage transfer from HENSM starting conditions **f.** Mouse and human βCAT^-/-^ ESCs were rendered transgenic with a validated Tamoxifen inducible βCAT^-^ ERT transgene. Acquisition or loss of naïve domed like morphology other parameters were assayed after 2 passages of the indicated conditions. **g.** Luciferase assay for measuring NFkB activity via luciferase reporter assay. Values were normalized to HENSM no SRCi conditions (defined as 1; n=3 per condition). *t-test *p* Value < 0.01; NS-not significant. **h.** 29-8 hESC line that carries GFP and tdTomato on each of the X chromosomes in the MECP2 locus, were rendered transgenic with IKB⍰ WT or dominant negative mutant allele (IKB⍰-DNmut), and cells were FACS analyzed following 10-day expansion in the indicated conditions. in the indicated conditions.

Given that prior studies did not rely on exacting reporter and knockout models for WNT signaling components and did not use teratoma competent cell lines, we turned to generate ßCAT KO cells on naïve pluripotency reporter cell lines to give a definitive answer for this fundamental question **(Fig. S27)**. ßCAT-KO hESCs maintained high levels of ΔPE-OCT4-GFP in HENSM condition, and upon removal of XAV939, GFP level was not decreased in BCAT-KO, but only in WT ESCs (**Fig. 5c**). Supplementing naïve cells with WNT stimulator, like CHIR99021, or depleting BCAT inhibitor (TNKi), activated WNT signaling in human PSCs (**Fig. 5d**) compromised ΔPE-OCT4-GFP levels (**Fig. 5c**), compromised their domed shape like morphology (**Fig. 5e**) and their naïve-like transcriptional profile (**Fig. 3a**). We introduced tamoxifen induced ßCAT-ERT transgene into ßCAT-KO hESCs, and we noted that in human ΔPE-OCT4-GFP signal, naïve transcription profile and domed morphology were compromised upon 4OHT stimulation (**Fig. 5f, Fig S26b-d**). This is in striking contrast to mouse ΔPE-Oct4-GFP ESCs expanded in N2B27 LIF conditions that upon tamoxifen treatment induced ΔPE-Oct4-GFP reporter and naive characteristic domed like morphology (**Fig. 5f**). Similarly, while KO of Tcf3 boosts mouse naïve pluripotency and alleviates the need for WNT stimulation (Wray et al., 2011b), TCF3 KO human ESCs (**Fig. S28a-b**) still required WNTi to maintain their naïve identity in HENSM conditions highlighting a major difference between mouse and human naïve pluripotency (**Fig. S28c-d**). Finally, supplementing HENSM conditions with CHIR compromised their ability to maintain pluripotency upon depletion of DNA and RNA methylation or omitting L-Glutamine from the culture conditions (**Fig. S28e**). Cross-species comparison of *in vivo* mouse and human RNA-seq data of pre-implantation blastocysts shows a strong signature for AXIN1/2 upregulation in human but not mouse ICM (**Fig. S28f**). Collectively, these findings clearly establish WNT-ßCAT signaling as a major priming agent for human, but not mouse, naïve pluripotency and establish that ablation of ßCATENIN can substitute for the need for Tankyrase or WNT pathway inhibition.

SRC inhibition has been shown previously to deplete activation of downstream effectors including ERK, PKC and NFkB signaling (Lee et al., 2007; Lluis et al., 2007). Given that SRCi was needed in HENSM conditions despite continued direct blocking of ERK and PKC pathways, this has led us to focus on NFkB as a potential effector mediating the beneficial effect of the use of SRCi (Torres and Watt, 2008). Luciferase reporter assay activity showed pronounced activity for NFkB signaling in primed conditions in comparison to HENSM naïve expanded cells (**Fig. 5g**), and activity of NFkB became prominently induced upon omission of SRC in HENSM conditions (**Fig. 5g, Fig. S29a**). The transfection of dominant negative IkBl⍰ subunit, which blocks NfKB signalling, in ßCAT-KO;ΔPE-OCT4-GFP hESCs allowed maintenance of ΔPE-OCT4-GFP not only in HENSM without TNKi, but also without SRCi (**Fig. 5h**). These results establish that WNT-BCAT and SRC-NFkB pathways compromise human naïve pluripotency. Further, the results on ΔPE-OCT4-GFP, RT-PCR and DNA hypomethylation establish that ACTIVIN A, rather than BMP4, supports human naïve pluripotency (**Fig. 1h, 3c and 4d-e**). The latter can be consistent with the observations that single cell-RNA-seq data on mouse and human ICMs show that human ICM retains high NODAL expression but not that of BMP4 which is highly expressed only in the mouse ICM (Blakeley et al., 2015a; Weinberger et al., 2015).

In mouse ground state naïve conditions, LIF/Stat3 has been shown to be a booster for naïve marker expression however they can be omitted without entire collapse of the naïve PSC circuit (Ying et al., 2008b). By omitting LIF from HENSM conditions and by generating STAT3 KO human naïve PSCs, we show that LIF can slightly boost the purity of undifferentiated cells in culture (**Fig. 5a**) and naïve marker expression by RT-PCR (**Fig. S29a**) however it is dispensable and human naive PSCs can maintain their naïve identity even in the absence of LIF/STAT3 signaling (**Fig. S29b-d**) as has been previously shown for rodent ground state naïve PSCs in 3i conditions (3 inhibitors of ERK, GSK3 and FGFR) (Ying et al., 2008b).

### KLF17 is functionally essential for human, but not mouse, naïve pluripotency

Analysis of chromatin accessibility revealed differences in accessibility of DNA binding proteins and protein families, unique for each pluripotent configuration. Binding motifs of different TFs like KLF17 and KLF4 were found to be significantly enriched (corrected p-value ∼ 0) in naïve chromatin accessible loci whereas GATA family are more prone to be found in the primed condition (**Fig. 6a, Table S4**). We initially focused on KLF17 gene, which is specifically expressed in human naïve conditions and has been shown to promote ERV expression pattern in humans (Pontis et al., 2019) however the essentiality of its functional importance for maintaining human naïve circuitry remains to be established. We generated single and double KO ESC lines for KLF4 and KLF17 TFs (**Fig. S30**). Remarkably KO of KLF17 led to collapse of naïve pluripotency in HENSM conditions, while cells maintain their pluripotency in HENSM-ACT as measured by OCT4-GFP reporter (**Fig. 6b**). The latter also support the notion how ACTIVIN A supplementation promotes naïve pluripotency in humans. However, double KO naïve ESCs for KLF4/KLF17 deteriorated toward differentiation and most cells lost OCT4-GFP expression in both HENSM and HENSM-ACT conditions (**Fig. 6b**). In mice, Klf17 is not expressed in the mouse ICM or ESCs, but rather only until the 4 Cell stage *in vivo* (**Fig. S31a**). Newly generated Klf17 KO mice were viable and pups were obtained at the expected mendelian ratios and were fertile (**Fig. S31b-f)**. Unlike the strong phenotype seen in human KLF17 KO ESCs in HENSM conditions, mouse Klf17 KO ESCs were indistinguishable from WT cells and did not show any compromise in their naïve pluripotency marker expression (**Fig 6c, Fig. S31g**). These results establish a great dependency of human, but not mouse, naïve pluripotency on KLF17 that greatly stabilizes the TF circuitry.

**Figure 6.**
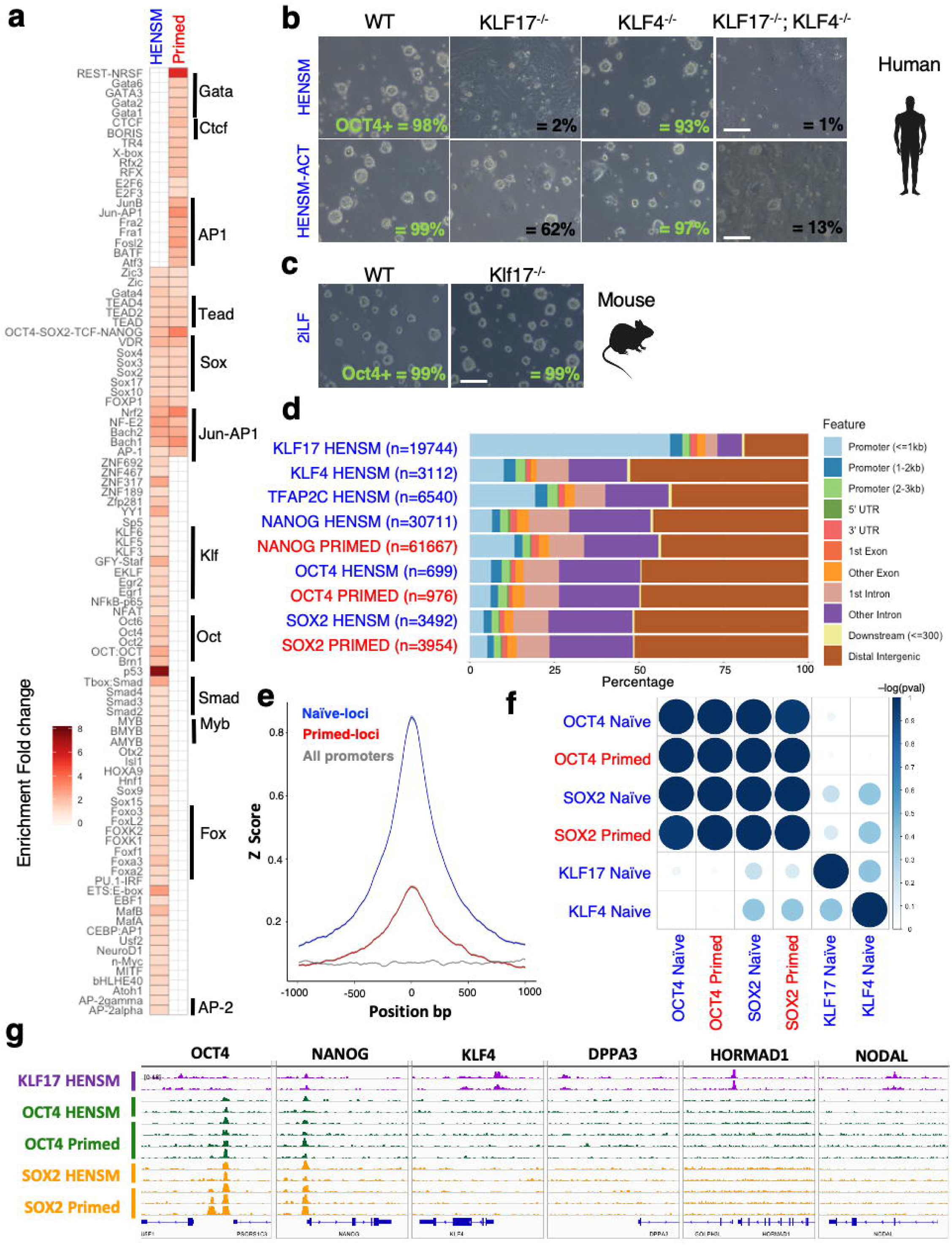
KLF17 is a master regulator of human naïve pluripotency. **a.** Motif enrichment in HENSM-specific and primed-specific accessible chromatin regions (n=20642 and b=36927, respectively). Motif families are indicated at the right. Shades represent enrichment fold-change. **b.** WT and KO hESCs for KLF4 and/or KLF17 were expanded in HENSM and HENSM-ACT conditions, imaged and assayed for OCT4-GFP by FACS analysis (values indicated per frame). **c**. WT and KO mESCs for Klf17 were expanded in 2iL, phase imaged and assayed for OCT4-GFP by FACS analysis (values indicated per frame). **d**. Genomic annotation of binding sites of KLF17, KLF4, TFAP2C, NANOG, OCT4 and SOX2 measured in HENSM naïve or primed conditions, showing the preference of KLF17 to bind promoters, compared to all TFs that prefer to bind enhancers (Distal intergenic + Introns annotations). **e.** KLF17 binding pattern (average z-score + STD) in naïve-specific regions (n=20642, blue), primed-specific regions (n=36927, red), or in all promoters (n=43463, grey). **f.** Overlap between target genes of the indicated transcription factors, showing highest overlap between Sox2 and Oct4 target genes (p-value∼0), and a relative lower overlap with KLF17 and KLF4 targets (p-value <10^−21^). Scaled p-value is presented (Methods) **g.** ChIP-seq profile of KLF17, OCT4 and SOX2 in selected regions and different conditions in human PSCs, showing that in some gene regions such as KLF4, DPPA3, HORMAD1 and NODAL; KLF17 is solely bound. IGV range common to all tracks is indicated.

To better understand the role of naïve specific pluripotency factors like KLF4 and KLF17 versus regulators of both naïve and primed cells like OCT4 and SOX2, we examined the DNA binding profile of the master pluripotent TF regulators OCT4 and SOX2 and naïve promoting TFs like KLF17, KLF4 and TFAP2C using ChIP-seq. KLF17 binds 12197 genes, mostly in their promoter (59% in <=1kb from TSS, **Fig. 6d**), unlike OCT4/SOX2 that bind typically to enhancers (72% and 85% in Distal intergenic regions and introns, respectively). KLF17 binds more substantially to naïve-specific open regions (**Fig. 6e**). Interestingly, OCT4 and SOX2, known to work as a complex (Merino et al., 2014), show high correlation between their binding targets, yet share significantly less reciprocal binding sites with other naïve specific KLF17 and KLF4 **(Fig. 6f).** The latter explain why despite the presence of OCT4 and SOX2 in human naïve PSCs, they could not compensate for depletion of KLF4 or KLF17 as the latter constitute major regulators of pluripotency specific genes and that are not bound by OCT4 or SOX2 (e.g. KLF4, DPPA3, NODAL, HORMAD1, **Fig. 6g, Table S5**). The latter also provides wider basis for the importance of KLF17 in human naïve pluripotency.

### Inhibition of NOTCH/RBPj pathway facilitates maintenance of human naïve pluripotency without ERK inhibition

As has been previously shown in mice (Choi et al., 2017; Yagi et al., 2017), the use of MEK/ERK inhibition is the major mediator for inducing global DNA hypomethylation which in turns leads to sporadic erosion of imprinting that gets more severe with extended passaging. In mice, using alternative naïve conditions that do not employ ERK inhibitor or titrating ERKi, allows isolating murine PSCs with all features of naivety except from global hypomethylation (Choi et al., 2017; Yagi et al., 2017). The latter murine cells are naïve by nearly all features and are capable of generate all-iPS mice with contribution to the germline, and thus provide a safer route for exploiting mouse naïve PSCs.

Although erosion of imprinting was slow and sporadic on few loci in HENSM conditions and only after extended passaging (**Fig. S13**), this may complicate the use of naïve cells in future clinical applications. We thus aimed to define **alternative** HENSM (**a**HENSM) conditions that allow naïve hPSC isolation and maintenance but without global DNA hypomethylation. Notably, recent studies showing titration of ERKi in 5i/LA conditions slightly improved the chromosomal aberrancies frequency per line in hESCs, still all lines had at least one genetic abnormality, and titrating ERKi levels in these conditions did not abrogate global hypomethylation and global loss of imprinting at all loci (Di Stefano et al., 2018). The latter was likely because these conditions still had BRAF inhibitor which is upstream to MEK/ERK pathway and contributes to the rapid and radical DNA hypomethylation induced in these cells (Pastor et al., 2016).

Withdrawal or partial (33%) titration of ERKi from HENSM compromised the naivety of human ESC as evident by a decrease in ΔPE-OCT4-GFP levels and loss of X-reactivation state in most of the cells within the expanded population (**Fig. 7a-b**). Supplementing ACTIVIN A upon omission of ERKi from HENSM conditions did not block loss of X reactivation in female cell lines (**Fig. 7d – lower panels**). Thus, we set out to screen for added compounds that would enable maintenance of pre-x inactivation upon omitting or titrating MEKi/ERKi (**Fig. 7c)**. Remarkably, we noted that adding of gamma secretase inhibitor DBZ, which blocks NOTCH pathway (Ichida et al., 2014), allowed robust and feeder free maintenance of human naïve cells when ERKi was omitted (**0**HENSM conditions) or titrated down from 1 μM to 0.33 μM termed (**t**HENSM conditions) (**Fig. 7d-e)**

**Figure 7.**
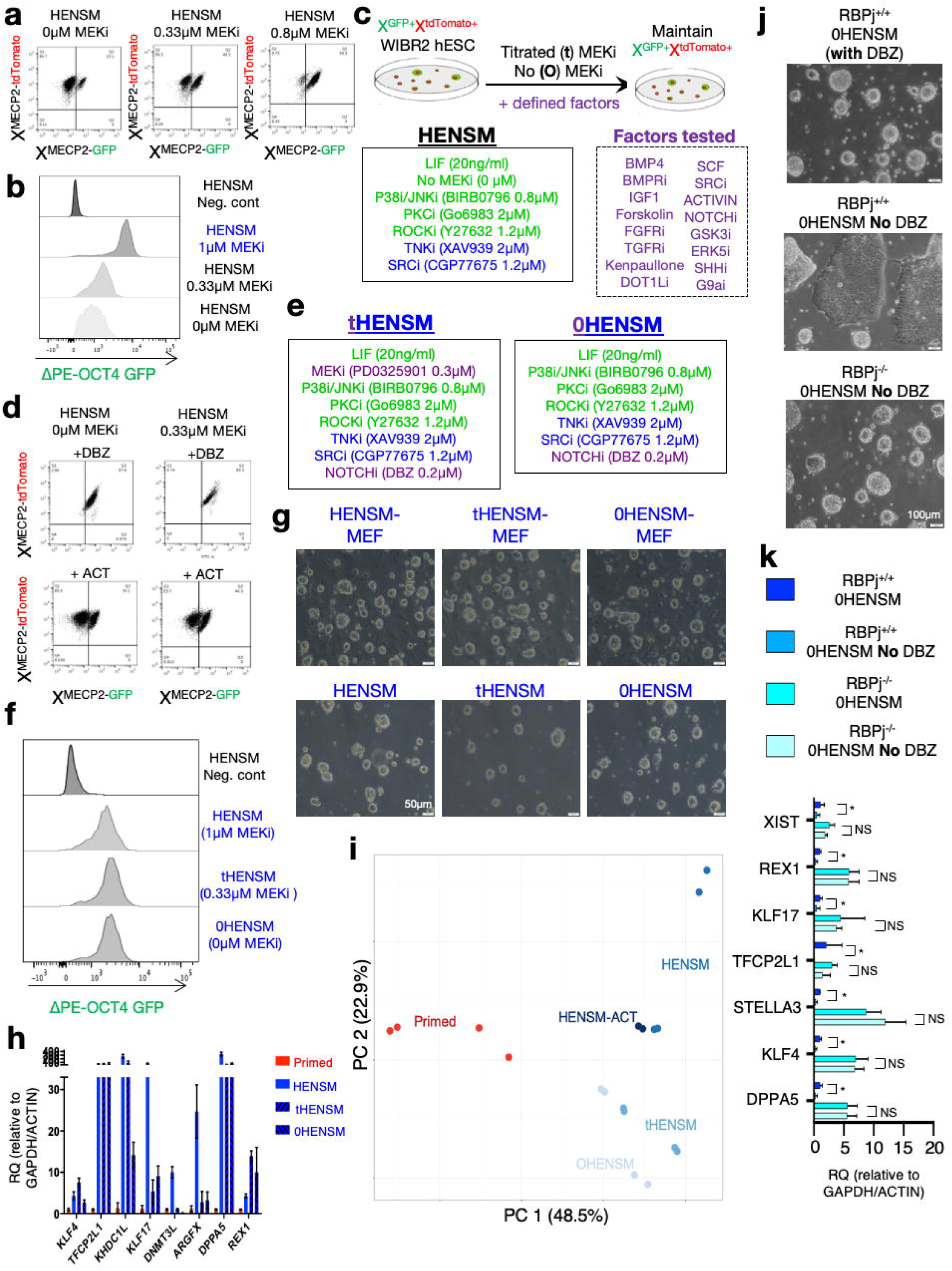
NOTCH/RBPj inhibition allows maintaining alternative human naïve cells without using MEK/ERKi. **a**. FACS analysis showing status of X activation in female 29-9 cells following decreasing concentrations of ERKi in HENSM conditions. Note that the fraction of cells inactivating X chromosome increases with depleting ERKi concentrations. b. FACS analysis for ΔPE-OCT4-GFP knock in naïve pluripotency reporter upon reducing ERKi in HENSM conditions. **c**. Schematic showing screen strategy for finding small molecule supplements that could allow maintaining GFP+/tdTomato+ cells in HENSM conditions in which ERKi is completely omitted (0HENSM) or partially depleted (tHENSM). **d.** FACS analysis following supplementing 0HENSM or tHENSM with ACTIVIN A or DBZ, a gamma secretase small molecule inhibitor that blocks NOTCH signaling (NOTCHi). **e**. Summary of small molecules and their concentrations used in the optimized tHENSM and 0HENSM conditions used herein. **f**. ΔPE-OCT4-GFP knock in naïve pluripotency reporter in HENSM, tHENSM and 0HENSM conditions. **g**. Phase images of WIBR1 hESCs in different MEF supplemented or feeder free naïve pluripotency conditions. **h**. RT-PCR analysis for naïve pluripotency markers in different naïve and primed conditions. Values were normalized to ACTIN and GAPDH. Primed expression levels were set as 1. **i.** PCA of gene expression profiles of cells grown in Primed condition (red), or in HENSM, HENSM-ACT, tHENSM or 0HENSM conditions (blue shades). **j.** Phase images of RBPj^+/+^ and RBPj^-/-^ isogenic WIBR3 hESCs upon removal of DBZ from OHENSM conditions. Arrows indicate flatted and loss of domed colony morphology. **k.** RT-PCR analysis for changes in naïve pluripotency marker expression in RBPj^+/+^ and RBPj^-/-^ isogenic WIBR3 hESCs upon removal of DBZ from OHENSM conditions. Values were normalized to ACTIN and GAPDH. RBPj^+/+^ HENSM sample expression levels were set as 1. *t-test *p* Value < 0.01; NS-not significant.

Human PSCs expanded in aHENSM conditions (0HENSM and tHENSM) maintain ΔPE-OCT4-GFP signal equivalent to HENSM conditions (**Fig. 7f)**, and maintained pre-X inactivation state in female cell lines (**Fig. 7d)**. RT-PCR analysis showed that the lines expressed naïve pluripotency markers (**Fig. 7h)**, although levels were less induced in comparison to when ERKi was used. Global gene expression analysis showed that the cells clustered with HENSM naïve PSCs rather than primed PSCs (**Fig. 7i, Fig S32a**). At the functional levels, the cells showed no enhanced tendency for obtaining chromosomal abnormalities and were competent in differentiating into PGCLCs in vitro (**Fig. S32e**) and making teratomas without any need for priming *in vitro* before injections (**Fig. S6a**). Cells could be maintained upon depletion of DNMT1, METTL3 or exogenous L-glutamine and these qualities depended on the presence of DBZ (**Fig. S32b-d)**. WGBS analysis showed that these alternative 0HENSM and tHENSM conditions did not upregulate DNMT3L enzyme and did not show trends of global hypomethylation that were seen in HENSM or HENSM-ACT even after extended passaging (**Fig. 4d-e, Fig. S33a**) and consistently did not show accelerated loss of imprinting compared to parental primed cells **Fig. S33b-c**). Generation of RBPj KO hESCs (**Fig. S34)** allowed expanding human naïve PSCs in tHENSM and 0HENSM without adding DBZ, proving that NOTCH/RBPj (Castel et al., 2013) is the main mediator of dismantling naïve pluripotency in human cells when ERKi is depleted (**Fig. 7j-k)**. These results show that, equivalent to what was obtained in rodent *in vitro* PSCs with t2iL and a2iL mouse conditions that endow naïve features in murine PSCs without compromising global DNA methylation and imprinting regulation (Choi et al., 2017; Yagi et al., 2017), this can now be obtained also with human naïve-like PSCs.

## Discussion

The findings reported herein consolidate the previously raised notion that different genetic backgrounds assume distinct states of pluripotency *in vitro*, the stability of which is regulated by endogenous genetic determinants that remain unknown, and can be modified by defined exogenous factors that support the naïve ground state of pluripotency (Reik and Surani, 2015). The stringency in requirement for these exogenous factors appears to be different among distinct species, as exemplified by the requirement for simultaneous inhibition of SRC, PKC, NOTCH and multiple MAPK kinase pathways in human naïve cells. However, certain pathway like WNT and ACTIVIN signaling are divergent between mice and humans, as only in humans WNT inhibition and ACTIVIN stimulation promote naïve pluripotency and contribute to the unique molecular and epigenetic configuration of naïve pluripotency described herein. The molecular basis for the latter findings remains to be mechanistically defined. Our results also suggest that epigenetic priming of human pluripotent cells is predominantly caused by supplementation of WNT and FGF.

The above results on the requirement for integrated WNT-SRC-PKCi to maintain human naive pluripotency can explain why previous described conditions that applied only some of these three essential components had compromised and unstable naïve pluripotency features. For example, Zambidis and colleagues indicated Tankyrase to promote human PSCs in ERKi containing conditions, however did not include PKCi, SRCi and still used WNT stimulation (Zimmerlin et al., 2016), which yields cells with low ΔPE-OCT4-GFP signal and lack of X reactivation. Similarly, conditions describing extended potential cells in mouse or humans did not utilize PKCi or SRCi and still used WNT activation together with TANKYRASE inhibition (Yang et al., 2017), explaining again why these cells have unique characteristics, they fail to contain naïve pluripotency defining features and cannot sustain stable *in vitro* naive pluripotency independent of repressive DNA and RNA methylation marks. 5iLA conditions did not include PKCi and did not actively inhibit WNT pathway (Theunissen et al., 2014).

Equivalent to what has been previously established for rodent ESCs in 3i conditions (Ying et al., 2008), we now establish the ability to expand transgene independent human naïve PSCs independent of any exogenous cytokine supplementation (i.e. without LIF or ACTIVIN) and by only inhibiting human PSC autologous driven signaling pathways with small molecule inhibitors. We can also expand bona fide human naïve pluripotent cells by omitting MEK/ERK inhibitors. It will be important to decipher the downstream regulators of pluripotency influenced by NOTCH/RBPj, SRC-NFkB and PKC pathways. We cannot exclude that alternative growth conditions may be devised to capture human naïve pluripotent cells with features similar to those described herein, or that HENSM conditions might be modified for improving and consolidating the extent of naïve features in human pluripotent cells. Indeed, X reactivation in the naïve conditions described here in still cannot yield random inactivation upon differentiation of human PSCs and this is similar to previously generated conditions (Sahakyan et al., 2016).

The fact that murine naïve cells, rather than primed cells, are tolerant to ablation of epigenetic repressors supports the concept of naïve pluripotency as a synthesis with a relatively minimal requirement for epigenetic repression (in comparison to primed pluripotent and somatic cells) (Geula et al., 2015; De Los Angeles et al., 2015). In this study, conditional mutants for METTL3 and DNMT1 were used for optimizing more stringent human naïve pluripotency growth conditions. Further, our results in mice and human indicate that the ability to tolerate complete and permanent ablation of such repressors may be one of the key features of naïve pluripotency across different species and might prove an efficient method for defining ground state of pluripotency from other mammalian species *in vitro* (e.g. monkeys, bovine, swine).

## Acknowledgements

This work was funded by Pascal and Ilana Mantoux; Nella and Leon Benoziyo Center for Neurological Diseases; David and Fela Shapell Family Center for Genetic Disorders Research; Kekst Family Institute for Medical Genetics; Helen and Martin Kimmel Institute for Stem Cell Research; Flight Attendant Medical Research Council (FAMRI); Helen and Martin Kimmel Award for Innovative Investigation; Dr. Beth Rom-Rymer Stem Cell Research Fund; Edmond de Rothschild Foundations; Zantker Charitable Foundation; Estate of Zvia Zeroni; European Research Council (ERC-CoG); Israel Science Foundation (ISF); Minerva; Israel Cancer Research Fund (ICRF) and BSF. We thank Dr. Rada Massarwa for help in setting up roller culture incubators. The experimental setting established here can be purchased from Arad Technologies Ltd., Ashdod, Israel. We thank D. Trono and J. Pontis for TE analysis, N. Benvenisty and S. Bar for RNA-seq based karyotyping and SNP analysis for LOI, R. Jaenisch for sharing WIBR2 X chromosome reporter cell lines, A. Meissner for sharing HUES64 with DNMT1 TET-OFF reporter cell line and constructs, A. Smith for H9-NK2 reset cell line, R. Massarwa for technical advice on embryo imaging, Q. Ying for mouse ßCatenin mutant cells and constructs.

## METHODS

### HENSM conditions

The following serum free conditions, termed HENSM (**H**uman **E**nhanced **N**aïve **S**tem cell **M**edium) were used to isolate, generate, derive and stabilize naïve human pluripotent stem cells (iPSCs and ESCs) with the unique biological properties described in this study. HENSM medium was generated by including: 240 ml Neurobasal (ThermoFisher 21103049) and 240ml of DMEM-F12 without HEPES (ThermoFisher 21331020), 5 ml N2 supplement (ThermoFisher 17502048 or in-house prepared), 5mL GlutaMAX (ThermoFisher 35050061), 1% nonessential amino acids (BI 01-340-1B), 1% Penicillin-Streptomycin (BI 03-031-1B), 10ml B27 supplement (Invitrogen 17504-044 or in-house made or Xenofree B27 Invitrogen A1486701 (for XF conditions)), 0.8mM Dimethyl 2-oxoglutarate (aKG) (add 60µL from purchased solution from Sigma 349631), Geltrex (Invitrogen A1413202/A1413302 - add 1ml rapidly in media to obtain 0.2% final conc), 50µg/ml Vitamin C (L-Ascorbic acid 2-phosphate sesquimagnesium salt hydrate - Sigma A8950), 20 ng/ml recombinant human LIF (Peprotech 300-05) and the following small molecule inhibitors: MEKi/ERKi (PD0325901 1 µM - Axon Medchem 1408); WNTi/TNKi (IWR1-endo 5µM - Axon Medchem 2510 or XAV939 2µM - Axon Medchem 1527); P38i/JNKi (BIRB0796 0.8µM - Axon Medchem 1358), PKCi (Go6983 2µM - Axon Medchem 2466), ROCKi Y27632 (1.2µM - Axon Medchem 1683), SRCi (CGP77675 1.2µM – Axon Medchem 2097). For HENSM-ACT, the latter HENSM conditions were supplemented with 4-20 ng/ml recombinant human ACTIVIN A (Peprotech 120-14E). We avoided using β-mercaptoethanol or adding additional BSA. Cells in HENSM were grown on 1% Growth factor reduced Matrigel (BD) or 1% GELTREX (ThermoFisher A1413202 or A1413302) or Biolaminin-511 (Biolamina Inc; including for HENSM-XF conditions) coated plates for at least 1 hour in 37°C. Notably, coated plates with 0.2% gelatin and irradiated ICR-DR4 MEFs can also be used to maintain human naïve cells in HENSM conditions. Enhanced single cell cloning efficiency can be obtained with HENSM supplementation with additional 5µM of Y-27632 ROCKi for 24 hours after cell passaging. TrypLE (ThermoFisher 12604013 or 12604054) (diluted into 0.5X - EDTA 0.75mM) is optimal for single cell passaging (2mL per 10cm dish, 0.75ml per 6 well - for 5 min at 37°C). After removal of TrypLE and air-drying at RT for 3 min., cells are resuspended in PBS. Rapid 0.05% trypsinization (0.5-1 min at 37C) is optimal for passaging as small clumps. Longer 0.05% trypsinization for 3-5 minutes at 37°C is optimal for microinjection into as the cells come out less sticky. Harvested cells should be centrifuged for 4 min x 1300RPM. Cells were expanded in 5% O_2_ and 5% CO_2_ conditions and passaged every 4-5 days. Medium was freshly replaced every 24 hours. Assembled media can be used for up to 7 days after preparation if kept at 4C. Passage numbers of naïve-hiPSC/hESCs indicate number of passages counted after induction or stabilization of the naïve state (and do not include previous passages when the cells were established and maintained in other conditions). For transfection of mouse and human naïve iPSCs/ESCs, 10 million cells were harvested with 0.05% trypsin-EDTA solution (Invitrogen), and cells resuspended in PBS+/+ were transfected with 100-150 μg DNA constructs (Gene Pulser Xcell System; Bio-Rad; 220V, 25μF, square wave, 20 ms, 4 mm cuvettes). Cell lines were monthly checked for Mycoplasma contaminations (LONZA – MYCOALERT KIT), and all samples analyzed in this study were not contaminated. Instead of adding 5ml of commercially available N2 supplements, individual components can be added to 500 ml media bottle at the indicated final concentrations: 1) Recombinant Human Insulin (Sigma I-1882) – 12.5 µg/ml final concentration; 2) Apo-Transferrin (Sigma T-1147) – 500 µg/ml final concentration; 3) Progesterone (Sigma P8783) – 0.02 µg/ml final concentration; 4) Putrescine (Sigma P5780) – 16 µg/ml final concentration; 5) Sodium Selenite (Sigma S5261) - add 5 µL of 3 mM stock solution per 500ml HENSM media. For **0HENSM** conditions – PD0325901 was omitted and replaced with DBZ (0.15-0.3 µM – TOCRIS 4489). For **tHENSM** conditions titrate concentration of ERKI were used (0.33microM PD0325901 - Axon Medchem 1408) together with DBZ (0.15-0.3 µM – TOCRIS 4489). Previously described NHSM conditions (Gafni et al., 2013a) were assembled as follows: 240 ml Neurobasal (ThermoFisher 21103049) and 240ml of DMEM-F12 without HEPES (ThermoFisher 21331020), 5 ml N2 supplement (ThermoFisher 17502048 or in-house prepared), 5mL GlutaMAX (ThermoFisher 35050061), 1% nonessential amino acids (BI 01-340-1B), 1% Penicillin-Streptomycin (BI 03-031-1B), 10ml B27 supplement (Invitrogen 17504-044), 50µg/ml Vitamin C (L-Ascorbic acid 2-phosphate sesquimagnesium salt hydrate - Sigma A8950), 20 ng/ml recombinant human LIF (Peprotech 300-05) and the following small molecule inhibitors: MEKi/ERKi (PD0325901 1 µM - Axon Medchem 1408); CHIR99021 (0.3-1µM - Axon Medchem 2510); P38i/JNKi (BIRB0796 2µM - Axon Medchem 1358), PKCi (Go6983 2µM - Axon Medchem 2466), ROCKi Y27632 (1.2µM - Axon Medchem 1683), 20 ng/ml recombinant human ACTIVIN A (Peprotech 120-14E) and 8ng/ml bFGF (Peprotech).

### Small molecule compounds and cytokines used in small scale screens

Small molecules and cytokines were supplemented as indicated in screening related figure legends: recombinant human BMP4 (10ng/ml, Peprotech), BMPRi (LDN193189 0.3µM, Axon Medchem 1509), recombinant human IGF1 (10ng/ml, Peprotech), Forskolin (FK, 10µM, TOCRIS), FGFR inhibitor PD173074 (0.1µM, TOCRIS), TGFRßi/ALK inhibitor A83-01 (0.3µM, Axon Medchem 1421), Kenopaullone (KP, 5µM, Sigma Aldrich), DOT1L inhibitor (SGC0946 0.5µM, SigmaAldrich SML1107), recombinant human SCF (10ng/ml, Peprotech), SRCi CGP77675 (CGP77675 1µM – Axon Medchem 2097), TNKi (IWR1-endo 4µM - Axon Medchem 2510), Gamma Secretase / NOTCH pathway inhibitor (DBZ 0.35µM – Axon Medchem 1488), BRAFi (SB590885, 0.5µM, Axon Medchem 2504), ERK5i (BIX02189 5µM, Axon Medchem 1809), SHHi (RU-SKI43 3µM, Axon Medchem 2035), G9a inhibitor (BIX01294 0.5µM – Axon Medchem 1692), Medium with inhibitors was replaced every 24 hours. Doxycycline (DOX - SigmaAldrich D9891) was used at a final concentration of 2µg/ml and refreshed every 48 hours upon media exchange. For exogenous L-GLUTAMINE withdrawal assay (“no GLUT assay”) (Carey et al., 2015b), media indicated were used without addition of non-essential amino acids, without L-Glutamax, without added aKG, and without Matrigel/Geltrex as some of these components can introduce L-Glut precursors as well. Laminin511 or Gelatin & irradiated feeders were used as coating material for no-GLUT assay.

### ESC derivation from human blastocysts in HENSM conditions

The use of human preimplantation embryos for ESC derivation was performed in compliance with protocols approved by LIS hospital ESCRO committee, Lis hospital Institutional review committee and Israeli National Ethics Committee (7/04-043) and following the acceptance of a written informed consent. The couples’ participation in the study was voluntary after signing informed consent forms and there was no monetary compensation for their embryo donation. Inner cell masses (ICMs) were isolated mechanically by laser-assisted micromanipulation from spare IVF embryos, at day 6-7 following fertilization. The intact ICM clumps were placed on a feeder cell layer of irradiation treated DR4 mouse embryonic fibroblasts and cultured in HENSM or HENSM-ACT media. Initial outgrowths of proliferating ESCs were evident by days 4-8, and were trypsinized into single cells, 6-10 days following ICM plating. The newly established cell lines were further propagated by 0.05% trypsin and then either frozen or used for further analysis.

### Culture of conventional/primed human ESCs and iPSCs

The following already established conventional human ESCs and iPSC lines were used (indicated passage number of the cell line taken for conversion into naïve pluripotency is indicated in parentheses): Human induced pluripotent stem cells C1 (P25) hiPSC lines and the human embryonic stem cell (hESC) lines H1 (P42), H9 (P39), WIBR1 (P51), WIBR2 (P44), WIBR3 (P42), HUES64, LIS1, WIS1, LIS2 hESCs were maintained in 5% O2 conditions (unless indicated otherwise) on irradiated mouse embryonic fibroblast (MEF) feeder layers or Matrigel coated plates as indicated. Two conventional and established conditions were used as indicated throughout the figures and manuscript: TeSR conditions on Matrigel coated plates (Stem Cell Technologies) or FGF/KSR conditions on MEF substrate: DMEM-F12 (Invitrogen 10829) supplemented with 15% Knockout Serum Replacement (Invitrogen 10828-028), 1mM GlutaMAX (Invitrogen), 1% nonessential amino acids (Invitrogen), 1% Sodium-pyruvate (BI 03-042-1B), 1% Penicillin-Streptomycin (BI 03-031-1B), 0.1mM β-mercaptoethanol (Sigma), and 8ng/mL bFGF (Peprotech) and 2ng/ml recombinant human TGFβ1 (Peprotech). Cultures were passaged every 5–7 days either manually or by trypsinization (24 hours pre and 24 hours after addition of ROCK inhibitor at 5-10µM concentration). For transfection of hiPSC and hESC lines, cells were cultured in Rho kinase (ROCK) inhibitor (Y27632) 24 h before electroporation. Primed human ESC and iPSC cells were harvested with 0.25% trypsin-EDTA solution (Invitrogen), and cells resuspended in PBS were transfected with 75μg DNA constructs (Gene Pulser Xcell System; Bio-Rad; 250 V, 500 μF, 0.4-cm cuvettes). Cells were subsequently plated on MEF feeder layers (DR4 MEFs for puromycin or neomycin selection) in hESC medium supplemented with ROCK inhibitor for the first 24 hours, and then antibiotic selection was applied. Previously described 5iLA or 4iLA were assembled and applied as previously described (Theunissen et al., 2016). T2iL-Go reset and PXGL conditions were used and assembled as previously described (Brendenkamp et al., 2019a; Takashima et al., 2014b). Media for extended potential stem cells or region specific primed cells were also devised where indicated as originally reported (Wu et al., 2015; Yang et al., 2017).

### Mouse naïve and primed stem cell lines and cultivation

Murine naïve V6.5 ESCs (C57B6/129sJae) pluripotent cells were maintained and expanded in serum-free chemically defined N2B27-based media: 240 ml Neurobasal (ThermoFisher 21103049) and 240ml of DMEM-F12 with HEPES (SIGMA D6421), 5ml N2 supplement (Invitrogen; 17502048), 5ml B27 supplement (Invitrogen; 17504044), 2mM glutamine (Invitrogen), 1% nonessential amino acids (Invitrogen), 0.1mM β-mercaptoethanol (Sigma), 1% penicillin-streptomycin (Invitrogen), 5mg/ml BSA (Sigma). Naïve 2i/LIF conditions for murine PSCs included 20 ng/ml recombinant human LIF (in-house made). Where indicated 2i was added: small-molecule inhibitors CHIR99021 (CH, 3μM-Axon Medchem) and PD0325901 (PD, 1μM – Axon Medchem 1408). Murine naïve ESCs and iPSCs were expanded on gelatin-coated plates, unless indicated otherwise. Primed 129Jae EpiSC line (derived from E6.5 embryos) or C57BL6/129sJae EpiSC line (derived following in vitro priming of V6.5 mouse ESCs) were expanded in N2B27 with 12ng/ml recombinant human bFGF (Peprotech) and 20ng/ml recombinant Human ACTIVIN A (Peprotech). Murine EpiSC were expanded on feeder free Matrigel coated plates. Cell lines were routinely checked for Mycoplasma contaminations every month (LONZA – MYCOALERT KIT), and all samples analyzed in this study were not contaminated.

### Reprogramming of somatic cells and cell infection

Virus-containing supernatants of the different reprogramming viruses: STEMCCA-OKSM polycistronic vector (DOX inducible and constitutive expression) was supplemented with the FUW-lox-M2rtTA virus (when necessary) and an equal volume of fresh culture medium for infection. For derivation of iPSCs directly from fibroblasts (and not from already established iPSC lines), fibroblasts were grown in the presence of DOX in HENSM conditions on Matrigel-coated plates until initial iPSC colonies were observed and subcloned. Generation of JH1 hiPSCs from human PBMCs was conducted by OSKM Sendai virus Cyto-Tune-iPS2.0 Kit according to manufacturer’s instructions but in HENSM conditions applied starting from day 4 of the reprogramming process.

### Sources of cell lines and plasmids

WIBR3-OCT4-GFP knock in reporter ESCs and ΔPE-OCT4-GFP WIBR3 hESCs were previously described (Theunissen et al., 2016). 29-8 WIBR2 and 29-9 WIBR2 hESCs reporters for X chromosome reactivation were previously described (Theunissen et al., 2016). Targeting strategy and CRIPS/Cas9 used for targeting of human ESCs are delineated in supplementary figures. Correct targeting was validated by southern blot analysis and normal karyotype was revalidated before use of all generated new lines or subclones. Klf17^-/-^ mouse ESCs (and its littermate control WT ESC) were derived from blastocysts obtained following mating of Klf17^-/-^ adult mice. ßCAT-ERT transgenic construct and ßCat-KO mESCs were previously described and validated (Kim et al., 2013). pBabe-Puro-IKBalpha-mut (super-repressor) (dominant negative mutant) (Addgene 15291) and its control pBabe-Puro-IKBalpha-WT (Addgene 15290) were used to generate transgenic lines analyzed in Fig. 5h. TET-OFF DNMT1 HUES64 hESC line was previously described and validated (Liao et al., 2015). H9 hNanog-pGZ hESC that contains NANOG-GFP reporter was obtained from WiCell.

### Mouse embryo micromanipulation and imaging

Human naïve pluripotent iPSCs were treated with 20μM ROCKi the night before injection and trypsinized and microinjected into E2.5 BDF2 diploid morulas or aggregated with 2-Cell BDF2 embryos, and were subsequently allowed to develop *in vitro* until E3.5 (in KSOM medium). Microinjection into E2.5 morulas placed in M16 medium under mineral oil was done by a flat-tip microinjection pipette. A controlled number of 4-8 cells were injected into the morulas. After injection, morulas were returned to KSOM media (Invitrogen) supplemented with 10-20 um ROCKi for 4h, then transferred to KSOM without ROCKi and placed at 37°C until transferred to recipient females at E3.5. Ten to fifteen injected blastocysts were transferred to each uterine horn of 2.5 days post coitum pseudo-pregnant females. 120-150 embryos were obtained and transferred for each injection batch. Embryos were dissected at the indicated time points and imaged for GFP+ cell localization. The latter experiments were approved by Weizmann Institute IACUC (00330111-2) and by the Weizmann Institute Bioethics and ESCRO committee. *In Toto* confocal imaging of chimeric mouse embryos was conducted after that the embryos were removed from the uterus into Tyrode’s solution, the decidua removed and the embryo carefully isolated with yolk sac intact. Embryos were moved into a droplet of culture media on a paper filter attached to a coverslip with vacuum grease. Starting farthest away from the embryo, the yolk sac was gently peeled open and lightly pressed onto the paper filter, which has adherence qualities, to anchor the embryo to the filter. To expose the tissue of interest, the embryo can be stretched and lightly pressed onto the filter. After mounting the embryos, a glass-bottomed dish was placed over the embryos using vacuum grease drops for spacing. The dish was inverted, filled with culture media and placed in the imaging chamber on an inverted Zeiss confocal LSM700 microscope. No blinding or randomization was conducted when testing outcome of microinjection experiments. Mounted embryos were placed in a heat- and humidity-controlled chamber (37°C; 5% O2, 5% CO2 and 90% N2 gas mixture) on a motorized stage of an inverted Zeiss LSM700 confocal microscope. Images were acquired with a 20x/0.8 M27 lens with 0.5-0.6x digital zoom-out or 10x/0.3 Ph1 M27 lens with 0.5-0.6x digital zoom-out. For GFP detection, a 448-nm laser (30-50% power) was used. Orange-Tracker (Molecular Probes, cat #C2927. Used at 20 μM) was excited by a 555-nm laser (30% power). Hoechst (Sigma, cat #B2261, used at 10 μg/ml) was excited by a 405-nm laser (40% power). RRX dyes were excited by 594 laser (20% power) and far-red dyes were excited by 647 laser (30% power). Images were acquired at 1024×1024 resolution. The thickness of *z-slices* appears in the relevant figure legends.

### Imaging, quantifications, and statistical analysis

Images were acquired with D1 inverted microscope (Carl Zeiss, Germany) equipped with DP73 camera (Olympus, Japan) or with Zeiss LSM 700 inverted confocal microscope (Carl Zeiss, Germany) equipped with 405nm, 488nm, 555nm and 635 solid state lasers, using a 20x Plan-Apochromat objective (NA 0.8). All images were acquired in sequential mode. For comparative analysis, all parameters during image acquisition were kept constant throughout each experiment. Images were processed with Zen blue 2011 software (Carl Zeiss, Germany), and Adobe Photoshop CS4.

### Immunohistochemistry analysis of mouse embryos

Mouse embryos were harvested at the indicate time points, washed once in 1xPBS and fixated overnight in 4% PFA. Thereafter, embryos were washed three times in 1xPBS and dehydrated in sequential Ethanol dilutions. After standard paraffinization and embedding, tissue blocks were sectioned on a microtome and mounted onto Superfrost plus slides (Thermo Scientific, Menzel-Glaser), dried at 38° C for 4 hours and stored at 4° C until further processing. Tissue sections were stained for following indicated antibodies: goat anti-GFP (Abcam ab6673), chicken anti-GFP (Abcam ab13970), rabbit anti-Numa (Abcam ab84680), mouse anti-human nuclei (Millipore MAB1281), mouse anti-TUJ1 (Covance MMS 435P), goat anti-SOX2 (R&D AF2018), goat anti-SOX17 (R&D AF1924). All secondary antibodies were used from Jackson ImmunoResearch including donkey anti-rabbit Biotin and Streptavidin-Cy3 for signal enhancement. For PFE-staining, sections were rehydrated, antigen retrieved in Sodium citrate buffer and pressure cooker, rinsed in PBS and treated with 0.3% H2O2 to reduce background staining. After permeabilization in 0.1% Triton X-100 in PBS three times for 2 min., samples were blocked in 10% normal donkey serum in PBS in humidified chamber for 20 min. at RT. Slides were then incubated in the appropriate primary antibody diluted in 1% BSA in 0.1% Triton X-100 at 4 °C overnight. Sections were then washed three times (5 min each) in washing buffer (0.1% Triton X-100 in PBS) incubated with appropriate fluorochrome-conjugated secondary antibodies diluted in 1% BSA in 0.1% Triton X-100 at RT for 1 h in the dark. For human cell specific NUMA staining, signal enhancement was performed by usage of Biotin-SP conjugated 2nd antibody for 30 min and subsequent incubation with Streptavidin-Cy3 antibody for 30 min. All sections were washed three times in washing buffer for 10 min each, counter stained with DAPI for 20 min, rinsed twice in washing buffer and mounted with Shandon Immuno-Mount (Thermo Scientific, 9990412). For OCT-staining, embryos were fixated o/n in 4% PFA at 4°C, washed three times in PBS for 10 min each and submerged first in 15% Sucrose/PBS and then 30% Sucrose o/n at 4°C. The day after, samples were subjected to increasing gradient of OCT concentration in Sucrose/PBS followed by embedding in OCT on dry ice and stored at - 80°C until further processing. Cryoblocks were cut with LEICA CM1950 and washed once with 1xPBS and incubated with 0.3% H_2_O_2_ for 20 min. After permeabilization with 0.1% Triton X-100 in PBS for 10 min., slides were again washed three times with 1xPBS for 2 min. each and blocked with blocked in 10% normal donkey serum in PBS in humidified chamber for 20 min. at RT. Antibody incubations were carried out analogously to PFE stainings. All images were collected on a Zeiss (Oberkochen, Germany) LSM700 confocal microscope and processed with Zeiss ZenDesk and Adobe Photoshop CS4 (Adobe Systems, San Jose, CA). For FACS analysis of mouse embryos, they were harvested and washed once in PBS. Thereafter, they were trypsinized in 0.05% trypsin/EDTA for 5 min and filtered through a cell strainer in MEF-medium before subjected to FACS analysis using FACS ARIA III analyzer and sorter system. Non-injected mouse embryos were used for proper gating and exclusion of auto-fluorescence.

### Luciferase activity assays

WNT TOP-Flash reporter constructs (ten Berge et al., 2011) were used to determine the regulation pattern of WNT signalling activity and were electroporated into 0.5–3 × 10−6 cells along with the pRL-TK (Renilla) vector for normalization. Assays were performed 48 h later using the Dual-Glo Luciferase Assay System (Promega). The basal activity of the empty luciferase vector was set as 1.0. Luciferase assay for NF-kB Reporter activity was measured with NF-kB Reporter kit (BPS Bioscience #60614) according to manufacturer instructions and following using Lipofectamine Stem Transfection reagent (ThermoFisher Scientific).

### RT-PCR analysis

Total RNA was isolated using Trizol (Ambion life technologies) and phenol-chloroform extraction. 1 μg of total RNA was reverse transcribed using a High-Capacity Reverse Transcription Kit (Applied Biosystems) and ultimately re-suspended in 200 µl of water. Quantitative PCR analysis was performed in triplicate using 1/100 of the reverse transcription reaction in a Viia7 platform (Applied Biosystems). Error bars indicate standard deviation of triplicate measurements for each measurement. RT-PCR primers used herein are: XIST-Forward: GGGTTGTTGCACTCTCTGGA; XIST-Reverse: TCATTCTCTGCCAAAGCGGT; OCT4-Forward: GCTCGAGAAGGATGTGGTCC; OCT4-Reverse: CGTTGTGCATAGTCGCTGCT; SOX1-Forward: GGGAAAACGGGCAAAATAAT; SOX1-Reverse: TTTTGCGTTCACATCGGTTA; PAX6-Forward: TGTCCAACGGATGTGTGAGT; PAX6-Reverse: TTTCCCAAGCAAAGATGGAC; ZIC2-Forward: TGCCTCATAAAAAGGAACAC; ZIC2-Reverse: TGTCCATTTGTAAAACTCCG; NANOG-Forward: GCAGAAGGCCTCAGCACCTA; NANOG-Reverse: AGGTTCCCAGTCGGGTTCA; DNMT3L-Forward: TGAACAAGGAAGACCTGGACG; DNMT3L-Reverse: CAGTGCCTGCTCCTTATGGCT; KHDC1L-Forward: ACGAGTGCTCTCAGCAAGGA; KHDC1L-Reverse: GTACGTGTCATCAAGTCCGAAGA; ARGFX-Forward: CCGGAGTCAACAGTAAAGGTTTG; ARGFX-Reverse: GGTGGGCACATTCTTCTTGGA; KLF17-Forward: AGCAAGAGATGACGATTTTC; KLF17-Reverse: GTGGGACATTATTGGGATTC; TFCP2L1-Forward: TCCTTCTTTAGAGGAGAAGC; TFCP2L1-Reverse: ACCAACGTTGACTGTAATTC; KLF4-Forward: CGCTCCATTACCAAGAGCTCAT; KLF4-Reverse: CACGATCGTCTTCCCCTCTTT; STELLA-Forward: CGCATGAAAGAAGACCAACAAACAA; STELLA-Reverse: TTAGACACGCAGAAACTGCAGGGA; DPPA5-Forward: TGCTGAAAGCCATTTTCG; DPPA5-Reverse: GAGCTTGTACAAATAGGAGC; OTX2-Forward: CATCTGATCAAAGTTCCGAG; OTX2-Reverse: TTAAGCAGATTGGTTTGTCC; PRDM14-Forward: CCTGACACTTTAATTCCACC; PRDM14-Reverse: AAGGTAATAACTGGGAGGTC; FGF4-Forward: AGAATGGGAAGACCAAGAAG; FGF4-Reverse: TGCATTAAACTCTTCATCCG; REX1-Forward: GGAATGTGGGAAAGCGTTCGT; REX1-Reverse: CCGTGTGGATGCGCACGT; ACTIN-Forward: CCACGAAACTACCTTCAACTCC; ACTIN-Reverse: GTGATCTCCTTCTGCATCCTGT; GAPDH-Forward: CTCCTGCACCACCAACTGCT; GAPDH-Reverse: GGGCCATCCACAGTCTTCTG; VIMENTIN-Forward: TGTCCAAATCGATGTGGATGTTTC; VIMENTIN-Reverse: TTGTACCATTCTTCTGCCTCCTG; ESRRB-Forward: TCTCACCCAGCACTAGGACACCAG; ESRRB-Reverse: ACACCCACTTGCTCCAAGCCA; ESRRB-Taqman: Hs01584024_m1

### Detection of miRNA expression in human ESCs/iPSCs

RNA harvest was performed with miRNeasy Micro Kit (Qiagen) and cDNA synthesized with miScript II RT kit (Qiagen) using miScript HiFlex buffer. Real-time quantitative RT-PCR was performed using the miScript SYBR Green PCR kit (Qiagen) and a ViiA 7 Real-Time PCR System (Applied Biosystems). The primer concentration used was 1 µM in a final reaction volume of 10µl. Each reaction was performed in triplicate. The thermal cycling parameters were according to manufacturer orders (miScript SYBR green PCR kit). The following primers were used: miR302b: TAAGTGCTTCCATGTTTTAGTAG; miR200c: TAATACTGCCGGGTAATGATGGA; U6: CGATACAAGGCTGTTAGAGAG

### Human mitochondrial DNA PCR

DNA was harvested with home-made lysis buffer (100 mM Tris pH8.0, 0.5 mM EDTA, 0.2% SDS, 200 mM NaCl, 200 ug/ml Proteinase K), precipitated with isopropanol and washed 1x with 70% Ethanol and finally eluted in 10 mM Tris-HCL pH 8, 0.1 mM EDTA. 50 ng template was used per reaction with SYBR Green Mastermix in ViiA 7 Real-Time PCR System (Applied Biosystems). As endogenous control, UCNE expression was used and compared to amplification of human mitochondrial element DNA. Quantification was done with delta/delta ct-method. The following primers were used: UCNE-forward: AACAATGGGTTCAGCTGCTT; UCNE-reverse: CCCAGGCGTATTTTTGTTCT; hMIT-forward: AATATTAAACACAAACTACCACCTACC; hMIT-reverse: TGGTTCTCAGGGTTTGTTATAA.

### Immunofluorescence staining

Cells were grown for two days on glass cover slips (13mm 1.5H; Marienfeld, 0117530) fixed with 4% paraformaldehyde/phosphate buffer for 10 min at room temperature, washed three times with PBS, and permeabilized in PBS/0.1% Triton for 10 min. Cells were blocked with blocking solution (2% normal goat serum, 0.1% BSA in PBS/0.05% Tween) for 1h at RT and incubated with primary antibody diluted in blocking solution overnight at 4°C (Antibodies in this study have all been validated in the literature and by ourselves). Cells were then washed three times with PBS/0.05% Tween, incubated with secondary antibodies for 1 hour at room temperature, washed in PBS/0.05% Tween, counterstained with DAPI (1 μg/ml), washed again three times with PBS/Tween 0.05% and mounted with Shandon Immu-Mount (Thermo Scientific, 9990412), and imaged. All comparative experiments were done simultaneously. The following antibodies were use at the indicated dilutions: mouse anti-TRA-1-60 (Abcam ab16288, 1:500), mouse anti-TRA-1-81 (Abcam ab16289, 1:100), mouse anti-SSEA1 (Abcam ab16285, 1:100), mouse anti-SSEA4 (Abcam ab16287, 1:100), rat anti-SSEA3 (Abcam ab16286, 1:100), goat anti-NANOG (R/D AF1997, 1:50), rabbit anti-OCT3/4 (Santa Cruz SC9081, 1:400), mouse anti-OCT3/4 (Santa Cruz SC5279, 1:100), rabbit anti-KLF4 (Santa Cruz SC20691), mouse anti-ECAD (Abcam ab1416, 1:100), rabbit anti-NCAD (Abcam ab12221, 1:800), mouse anti-NR3B2 (R&D PPH670500, 1:1000), rabbit anti-TFE3 (SIGMA HPA023881, 1:500), rabbit anti-H3K27me3 (Abcam AB5603), goat anti-TFCP2L1 (R/D AF5726, 1:200), rabbit anti-NUMA (Abcam ab84680, 1:200), chicken anti-GFP (Abcam ab13970, 1:1000), rabbit anti-KLF17 (SIGMA HPA024629, 1:100), anti-TFAP2C (CSTH2320, 1:100, Cell Signaling), anti-STELLA (Santa Cruz, sc-376862, 1:100), anti-DNMT3L (Novus Biologicals, NPB2-27098, 1:200). All secondary antibodies were used from Jackson ImmunoResearch.

### Western blotting analysis

Whole-cell protein extracts were isolated from human cells with conventional RIPA lysis buffer and protein concentration assayed by BCA kit (Thermo Scientific). Blocking was carried out in 10% skim milk in PBST for 1 h. Blots were incubated with the following antibodies in 5% BSA in PBST: KLF4 (AF3158; 1:200; R&D), OCT4 (H-134; 1:1,000; Santa Cruz), NAANOG (397A; 1:1,000; Bethyl), METTL3 (A301-567A, 1:2000, Bethyl), HSP90beta (ab32568, 1:10000, Abcam), DNMT1 (ab87654, 1:1000, Abcam), GAPDH (ab181602, 1:10000, Abcam), ACTIN (ab6276, 1:10000, Abcam), UHRF1 (sc-373750, 1:1000, Santa Cruz), DGCR8 (10996-1-AP, 1:1000, Proteintech), P53 (D-01, courtesy from Varda Roter’s lab), β-catenin (sc-7963, 1:1000, Santa Cruz), TCF3 (CST2883, 1:1000, Cell Signaling), STAT3 (sc-482, 1:1000, Santa Cruz), KLF4 (sc-20691, 1:1000, Santa Cruz), RBPJ (C5 5313, 1:1000, Cell Signaling), TFAP2C (CSTH2320, 1:1000, Cell Signaling), STAT3 (sc-7993, 1:1000, Santa Cruz). Secondary antibodies were HRP-linked goat anti-mouse, goat anti-rabbit and rabbit anti-goat (1:10,000; Jackson). Blots were developed using SuperSignal West Pico Chemiluminescent Substrate (Thermo Scientific 34580).

### Whole-Genome Bisulfite Sequencing (WGBS) Library preparation

DNA was isolated from cells using the Quick-gDNA miniprep kit (Zymo). DNA (50ng) was then converted by bisulfite using the EZ DNA Methylation-Gold kit (Zymo). Libraries were prepared using the TruSeq kit (Illumina) and length distribution of each library was measured using the Bioanalyzer and product concentration was measured using Qubit Fluorometric Quantitation. For sequencing, the libraries, NextSeq 500/550 High Output v2 kit (150 cycles) was used. The following secure token has been created to allow review of record GSE125555 while it remains in private status: https://www.ncbi.nlm.nih.gov/geo/query/acc.cgi?acc=GSE125555

### Whole-Genome Bisulfite Sequencing (WGBS) analysis

The sequencing reads were aligned to the human hg19 reference genome (UCSC, 2009), using a proprietary script based on Bowtie2. In cases where the two reads were not aligned in a concordant manner, the reads were discarded. Methylation levels of CpGs calculated by WGBS were unified. Mean methylation was calculated for each CpG that was covered by at least 5 distinct reads (X5). Average methylation level was calculating by taking the average over all covered X5 covered CpG sites in that genome.

### ChIP-seq library preparation

Cells were crosslinked in formaldehyde (1% final concentration, 10□min at room temperature) or double crosslinked in DSG/formaldehyde (2mM DSG, 30 min at room temperature), then formaldehyde 1% final concentration, 10□min at room temperature) as detail bellow per antibody, and then quenched with glycine (2.5M 5□min at room temperature). Cells were lysed in 50□mM HEPES KOH pH□7.5, 140□mM NaCl, 1□mM EDTA, 10% glycerol, 0.5% NP-40 alternative, 0.25% Triton supplemented with protease inhibitor at 4□°C (Roche, 04693159001) for 10 min, and later centrifuged at 950*g* for 10□min. Supernatant was discarded and pellet was resuspended in RIPA-1 (0.2% SDS, 1□mM EDTA, 0.1% DOC, 140□mM NaCl and 10 mM Tris-HCl) with protease inhibitor. Cells were then fragmented with a Branson Sonifier (model S-450D) at −4□°C to size ranges between 200 and 800□bp and centrifugation at max speed for 10 min. Supernatant lysate was extracted and diluted with RIPA 2-3-fold (0.1% SDS, 1□mM EDTA, 0.1% DOC, Triton 1%, 140□mM NaCl and 10 mM Tris-HCl). Small amount of lysate were saved for whole cell extract at this point. Antibody was pre-bound by incubating with Protein-G Dynabeads (Invitrogen 10004D) in blocking buffer (PBS supplemented with 0.5% TWEEN and 0.5% BSA) for 1□h at room temperature. Washed beads were added to the lysate for incubation as detail per antibody. Samples were washed five times with RIPA buffer, twice with RIPA buffer supplemented with 500□mM NaCl, twice with LiCl buffer (10□mM TE, 250mM LiCl, 0.5% NP-40, 0.5% DOC), once with TE (10Mm Tris-HCl pH 8.0, 1mM EDTA), and then eluted in 0.5% SDS, 300□mM NaCl, 5□mM EDTA, 10□mM Tris HCl pH□8.0. Eluate was incubated treated sequentially with RNaseA (Roche, 11119915001) for 30□min in 37°C and proteinase K (NEB, P8102S) for 2□h in 37°C and de-crosslinked in 65□°C for 8□h. DNA was purified with Agencourt AMPure XP system (Beckman Coulter Genomics, A63881). Libraries of cross-reversed ChIP DNA samples were prepared according to a modified version of the Illumina Genomic DNA protocol, as described previously (Rais et al., 2013). <u>Treatment per antibody:</u> KLF17 – double crosslinked, cell amount: 30 million, antibody: 6ug HPA024629 - atlas antibodies, incubation time: overnight. KLF4 – crosslinked, cell amount: 30 million, antibody: 10ug AF3158-R&D systems, incubation time: overnight, comment: This is a mouse Klf4 antibody that was verified to work on human using IF staining. NANOG – crosslinked, cell amount:30 million, antibody: 6ug AF1997-R&D systems, incubation time: overnight. OCT4 – crosslinked, cell amount:30 million, antibody: 10ug SC8628 Santa-Cruz, incubation time: overnight. SOX2 – crosslinked, cell amount:30 million, antibody: 10ug AF2018-R&D systems, incubation time: overnight. TFAP2c – crosslinked, cell amount:30 million, antibody: 5ug sc-8977 Santa-Cruz, incubation time: overnight. H3K27ac – crosslinked, cell amount:5 million, antibody: 5ug ab4729 Abcam, incubation time: 6 hours.

### ChIP-seq analysis

We analyzed Chip-seq data of the following DNA-binding proteins: NANOG, SOX2, OCT4, KLF4, KLF17, TFAP2C, H3K27AC, in HENSM or primed conditions, with replicates or triplicates for each antibody per single condition. For alignment and peak detection, we used bowtie2 software to align reads to human hg19 reference genome (UCSC, 2009), with default parameters. We identified enriched intervals of all measured proteins using MACS version 1.4.2-1. We used sequencing of whole-cell extract as control to define a background model. Duplicate reads aligned to the exact same location are excluded by MACS default configuration. The number of peaks in Primed conditions were in-par with previous publications (e.g. PMID <u>25409831</u>, PMID 19829295, **Supplementary Table S5**). Peaks that were observed in at least two replicates were kept for further analysis. Peaks were assigned to genes using Homer software (Supplementary Table S5). Motif enrichment analysis was performed using Homers findmotifsgenome.pl, with parameter -size 50 (**Supplementary Table S4**). Significance of overlap between TF targets was calculated using Fisher exact test **(Figure 6h)**.

### RNA-seq library preparation

Total RNA was isolated from indicated cell lines and extracted from Trizol pellets by chloroform-phenol extraction protocol, then cleaned using Ribo-Zero (Illumina Ribo-Zero Plus rRNA Depletion Kit, ID-20037135) or Poly-A (Dynabeads mRNA DIRECT Kit (Invitrogen), ID-61012) depletion Kits. Cleaned RNA was next utilized for RNA-Seq by ScriptSeq Preparation Kit v2 (Illumina) according to manufacturer’s instruction.

### RNA-seq analysis

hESCs grown in naïve and primed conditions, from different cell lines (LIS41, LIS49, WIBR2, WIBR3, H1, H9, HUES64) were used for RNA-seq analysis. Overall 36 samples were sequenced using poly-A single-end strategy, and 19 samples were sequenced using ribo-zero paired-end strategy. STAR software version 2.5.2b was used to align reads to human GRCh38 reference genome (2013), using the following flags: --outFilterMultimapNmax 1 --outReadsUnmapped Fastx --twopassMode Basic -- outSAMstrandField intronMotif. Read count values were estimated with HTSeq V0.7.2 software over all aligned reads using GRCh38 general feature format (GFF), with the following flags: -a 10 -s no -t exon -i gene_id. Mitochondrial and ribosomal genes were filtered out. Genes with a sum count across all samples <10, were filtered out as well. The filtering was done independently in each analysis, therefore the number of genes included may change, as it is dependent on the samples that were included for that analysis. Count values were normalized using R DESeq2 V1.26 software, and corrected for batch effects using R Limma V3.42. Variance stabilizing transformation was performed to all datasets as well. Dataset analysis and plotting was performed by R V3.6. Sample correlation was performed using Spearman method. Hierarchical clustering was carried out using “complete” or “Ward.D2” methods. Heatmap plots were first Z-scored by sample to better emphasize gene separation. Differentially expressed genes between Naïve and Primed samples were performed by the following parameters: log2(Fold change)> 2 or log2(Fold change) < −2, and adjusted p-value< 0.1. RNA-Seq data are deposited under GEO no. GSE125555.

### ATAC-seq library preparation

Cells were trypsinized and counted, 50,000 cells were centrifuged at 500*g* for 3 min, followed by a wash using 50 μl of cold PBS and centrifugation at 500*g* for 3 min. Cells were lysed using cold lysis buffer (10 mM Tris-HCl, pH 7.4, 10 mM NaCl, 3 mM MgCl_2_ and 0.1% IGEPAL CA-630). Immediately after lysis, nuclei were spun at 500*g* for 10 min using a refrigerated centrifuge. Next, the pellet was resuspended in the transposase reaction mix (25 μl 2× TD buffer, 2.5 μl transposase (Illumina) and 22.5 μl nuclease-free water). The transposition reaction was carried out for 30 min at 37 °C and immediately put on ice. Directly afterwards, the sample was purified using a Qiagen MinElute kit. Following purification, the library fragments were amplified using custom Nextera PCR primers 1 and 2 for a total of 12 cycles. Following PCR amplification, the libraries were purified using a QiagenMinElute Kit and sequenced.

### ATAC-seq analysis

Reads were aligned to hg19 human genome using Bowtie2 with the parameter -X2000 (allowing fragments up to 2 kb to align). Duplicated aligned reads were removed using Picard MarkDuplicates tool with the command REMOVE_DUPLICATES=true. To identify chromatin accessibility signal we considered only short reads (≤ 120bp) that correspond to nucleosome free region. To detect and separate accessible loci in each sample, we used MACS version 1.4.2-1 with --call-subpeaks flag (PeakSplitter version 1.0). To generate the final ATAC loci of naïve and primed conditions, we selected peaks that appear in the majority of the replicates of each condition, i.e. 3 out of 4 naïve samples, and 2 out of 3 primed samples.

### Enhancer Identification

H3K27ac peaks were detected using MACS version 1.4.2-1 and merged for each condition (naïve and primed) using bedtools merge command. All ATAC peaks were filtered to include only peaks which co-localized with the merged H3K27ac peaks in at least one condition. Finally, peaks that co-localized with promoter or exon regions based on hg19 assembly (UCSC, 2009) were filtered out. Finally, we were left with defined genomic intervals which we annotated as enhancers.

### SNP Score Calculation from WGS

SNP score calculation was done as previously described (Bar et al., 2017). Briefly, we downloaded WGS data for H1 and H9 cell lines from the SRA database (http://www.ncbi.nlm.nih.gov/sra). The files were extracted using SRAtools and aligned to the hg19 reference genome using BWA. SAMtools was used to convert from SAM to BAM. We then detected polymorphisms with the same pipeline as described previously for RNA-seq (without filtering for FPKM). In order to test if genomic SNPs are properly detected in the RNA-seq analysis, we extracted SNPs from exons of control pluripotent genes and calculated the average percentage of overlapping SNPs between WGS and RNA-seq of the appropriate H1 and H9 samples. In order to test if there is a difference in the existence of genomic SNPs between imprinted genes governed by maternal and paternal gDMR, we extracted the SNPs located at exons of imprinted genes from WGS of H1 and H9 cell lines and then calculated the percentage of genes with genomic SNPs in each group. We used a chi-square test, which confirmed that the bias was not significant.

### Quantifying X:A allelic ratio

X:A allelic ration was done as previously described (Bar et al., 2019). Briefly, to generate a quantifiable score of allelic expression across the X chromosome, we developed the X:Autosomes (X:A) allelic ratio. For this we first counted the number of biallelic SNPs in each chromosome. Since our analysis is based on expressed SNPs, this sum is expected to be dependent on the number of genes in the chromosome. Therefore, to overcome this bias we generated a normalized SNP sum by dividing the SNP sum with the number of known genes in each chromosome. We validated that the average of this normalized SNP sum was similar across all autosomes. Next, for each sample we calculated the ratio of normalized SNP sum of X chromosome and the average normalized SNP sum of autosomes:

### NS (normalized sum)=Sum(biallelic SNPs) / No. of genes in chromosome X : A allelic ratio = X chromosome-NS / Average (Autosome-NS)

By comparing the normalized sum of X and autosomes within each sample, we avoid potential biases which are due to coverage and batch issues. Samples with very low normalized SNP sums in autosomes (for example haploid samples or their diploid genetic match which are completely homozygous), were discarded from this analysis.

### Transposon expression profiling

TE profiling was performed as previously described (Theunissen et al., 2016). For repetitive sequences an D. Trono lab curated version of the Repbase database was used (fragmented LTR and internal segments belonging to a single integrant were merged). TEs counts were generated using the multiBamCov tool from the bedtools software. TEs which did not have at least one sample with 20 reads (after normalizing for sequencing depth) were discarded from the analysis. TEs overlapping exons were also removed from the analysis. Normalisation for sequencing depth and differential gene expression analysis was performed using Voom (Law et al., 2014) as it has been implemented in the limma package of Bioconductor (Gentleman et al., 2004). A gene (or TE) was considered to be differentially expressed when the fold change between groups was bigger than 2 and the p-value was smaller than 0.05. A moderated paired t-test (as implemented in the limma package of R) was used to test significance. P-values were corrected for multiple testing using the Benjamini-Hochberg’s method (Benjamini, 1995). Single cell RNA-Seq data was downloaded from GEO (Edgar et al., 2002) (GSE36552). Data was mapped and processed as explained above. When merging with in-house RNA-Seq, only TEs which were expressed in both datasets were used. To be considered expressed, the TE needed to have at least as many reads across all samples as samples existed in each dataset. Correspondence between naive/primed ESCs and single cell expression data from human embryonic stages ws also done as described before (Theunissen et al., 2016). For every cell state we perform a statistical test to find the genes (or TEs) that have a different expression level compared to the other cell states (Yan et al., 2013). For this we use a moderated F-test (comparing the interest group against every other) as implemented in the limma package of Bioconductor. For a gene (or TE) to be selected as expressed in a specific cell state it needs to have a significant p-value (<0.05 after adjusting for multiple testing with the Benjamini and Hochberg method) and an average fold change respective to the other cell states greater than 10. Note that with this approach a gene (or TE) can be marked as expressed in more than one cell state. Once we have the genes (or TEs) that are expressed in a specific cell state we ask if those genes are more expressed in primed or in naive. For that we see how many of the genes are up (or down) regulated in the primed/naive pairwise comparison. For a gene to be considered differentially expressed it needs to have a p-value (after multiple testing correction with the Benjamini and Hochberg method) lower than 0.05 and a fold change greater than 2.

## Supplementary Figure Legends

**Figure S1. Reporter systems for screening for enhanced human naive pluripotency conditions.**

**a.** W3G4 METTL3 TET-OFF cells were maintained in the presence of DOX for up to 4 passages in different conditions and stained for OCT4 to quantify percentage of cells that retained their pluripotency. Graph shows that NHSM conditions without feeder cells and other previously described naïve and primed conditions for human ESCs/iPSCs failed to maintain pluripotency in majority of cells expanded in the presence of Dox. **b.** METTL3 TET-OFF and DNMT1 TET-OFF cells were maintained in the presence of DOX for up to 4 passages in different previously published naïve and primed conditions and stained for OCT4 to quantify percentage of cells that retained their pluripotency. **c**. Quantification of ΔPE-OCT4-GFP knock in naïve pluripotency reporter, in variety of primed (red) and naïve conditions (blue). Mean fluorescence intensity values (MFI) are indicated. Figure shows that supplementing NHSM conditions with TNKi like IWR1 small molecules boosts expression of GFP suggesting enhancement of naivety characteristics. **d**. Quantification of ΔPE-OCT4-GFP knock in naïve pluripotency reporter, in optimized naïve conditions and various concentrations of GSK3 inhibitor that leads to WNT activation (CHIR99021 is used as GSK3 inhibitor and is abbreviated as CHIR). Figure indicates that CHIR addition negatively influences human naive pluripotency as determined by ΔPE-OCT4-GFP intensity. **e.** RT-PCR analysis for naïve pluripotency markers in HENSM conditions with and without P38i/JNKi (BIRB0796). Values were normalized to ACTIN and values in Primed conditions were set as 1. *t-test *p* Value < 0.01. Figure indicates that use of BIRB0796 as P38i/JNKi boosts expression of naïve pluripotency markers. **f.** W3G4 METTL3 TET-OFF cells were maintained in the presence of DOX for up to 4 passages in the optimized HENSM conditions but without TNKi, in order to see if other molecules can substitute for TNKi after all optimizations were applies (e.g. adding of Geltrex, concentration optimization). **g.** FACS analysis for OCT4-GFP pluripotency reporter expression following addition of TGFRi. In optimized NHSM conditions that still lack SRCi, pluripotency is entirely and rapidly lost upon inhibition of TGFRi. **h.** Phase images showing how supplementation of 0.2% Geltrex (Life Technologies) in the growth media and SRCi additively enhance domed like morphology of human naïve PSCs in HENSM conditions optimized herein.

**Figure S2. Functional engineered systems for evaluating HENSM conditions.**

**a.** Immunostaining for W3G4 before and after DOX validating METTL3 protein downregulation after DOX addition in HENSM. **b**. Immunostaining of W3G4 cells in HENSM conditions with and without DOX. Cells expressed canonical OCT4 and naïve pluripotency specific markers like KLF17 in both conditions. **c**. Phase images of WIBR3 esc in NHSM and HENSM conditions, showing more domed and uniform morphology in HENSM conditions. **d**. Efficiency of generating KO and targeting DGCR8 in human primed (TeSR) and HENSM naïve conditions. 2 replicates for targeting were done for each growth conditions. Only in HENSM conditions we recovered DGRC8 KO clones based on genotyping and western blot analysis. **e**. CRISPR targeting strategy for DGCR8 locus, followed by western blot and sequencing validation of KO clones. **f**. RT-PCR analysis for the indicated microRNA in DGCR8 WT and KO human ESCs. Figure shows loss of microRNA expression in DGCR8 KO human naïve HENSM conditions as expected following ablation of DGCR8. *t-test *p* Value < 0.01. Error bars indicate SD from mean. **g.** FACS analysis showing preservation of ΔPE-OCT4-GFP naïve marker expression in both WT and DGCR8 KO human ESCS expanded in HENSM conditions**. h**. Representative images for human DNMT1 TET-OFF engineered cells in different conditions with or without DOX addition. Previously described primed and naïve conditions do not support maintain their pluripotency and viability when DOX is added (DNMT1 is depleted). Only HENSM and HENSM-ACT conditions (with and without irradiated feeder cells – MEFs) maintain robust expansion of dome-like undifferentiated human PSCs *in vitro* even after extended passaging in the presence of DOX. **i**. WGBS validates global reduction in CG methylation in HENSM conditions following DOX addition (shutdown of DNMT1 expression). **j**. Immunostaining validating OCT4+ expressed in DNMT1 TET-OFF cells expanded in HENSM + DOX conditions at P10. **k**. METTL3 TET-OFF and DNMT1 TET-OFF cells were maintained in the presence of DOX for up to 4 passages in different HENSM based conditions upon depletion of the indicated key inhibitors and stained for OCT4 to quantify percentage of cells that retained their pluripotency. **l**. Oxygen consumptions rate (OCR) measurement in different conditions. **m**. OCT4+ colony formation in 2-deoxyglucose (2DG). 1mM 2DG was added to the indicated conditions. Error bars indicate SD.

**Figure S3. Optimized HENSM conditions enable naïve pluripotency maintenance in the absence of L-Glutamine**.

**a.** FACS based validation of pluripotency maintenance in human stem cells expanded in HENSM conditions with and without exogenous L-Glutamine. Percentage of positive cells and intensity of naïve marker expression were not compromised upon omitting GLUT in HENSM based conditions. **b.** Representative phase contrast and fluorescent images of human cells expanded in HENSM conditions with and without exogenous L-Glutamine (GLUT). TeSR primed human ESCs were used as controls. **c.** human ESCs expanded for 10 passages in HENSM conditions without exogenous GLUT robustly formed teratomas (without any need for exogenously induced priming before they were injected).

**Figure S4. Derivation of new human ESC lines in HENSM conditions.**

**a**. 5 new lines were derived in HENSM or HENSM-ACT conditions directly from human blastocysts as indicated. At P6-8, a small portion of the cells was taken and expanded in primed conditions (and thus are labeled as “primed cells”. **b**. Representative images showing previously established human primed/conventional ESCS were transferred to HENSM conditions, and after at least 5 passages a small portion of them were transferred back into primed conditions (thus are referred to as “reprimed” cells). **c**. Newly derived iPSC lines from dermal fibroblasts or peripheral blood mononuclear cells (PBMCs) were obtained following OSKM transduction and cell culturing in HENSM conditions. Phase images show initial iPSC colony appearance before they were picked for further expansion.

**Figure S5. Pluripotency marker expression and characterization in HENSM conditions.**

Representative immunostaining for pluripotency markers in HENSM conditions are shown for LIS49 hESC line. Primed cells expanded in TeSR conditions are used as controls. Note that KLF17, DNMT3L, STELLA and TFCP2L1 are naïve pluripotency specific markers and are expressed in HENSM conditions and not in primed cells.

**Figure S6. HENSM conditions maintain teratoma formation competence of PSCs.**

**a.** Mature teratoma images are shown following their derivation from the indicated cell lines expanded in the different indicated conditions. Please note that without exception, all teratomas were formed following direct subcutaneous injections after being expanded only in the indicated media condition and without the need for any expansion in other primed conditions *in vitro* before injection. **b**. Results of e-karyotyping based on RNA-seq data for the indicated lines are shown.

**Figure S7. Chromosomal stability following long term expansion in HENSM based conditions.**

Metaphase chromosomal spreads are shown from the indicated human ESC and iPSC lines expanded in HENSM (**a**), tHENSM (**b**) and OHENSM (**c**) based conditions. Passage numbers are indicated throughout.

**Figure S8. Differentially expressed genes between human naïve and primed states highlights regulatory candidates.**

**a**. Volcano plot comparing change in expression of all genes (log2(Naïve HENSM/primed Fold-Change) in x-axis), to their statistic (-log10(q-value) in y-axis). Differentially expressed genes (Fold-change>2(<0.5), p-adjusted<0.1) are marked in red. Extreme genes are highlighted. **b.** Spearman correlation matrix of naïve HENSM and primed samples, along with previously published naïve and primed samples (Takashima et al, 2014). **c.** Clustered expression profile of differentially expressed genes (Fold change>2(<0.5), p-adjusted<0.1, n=2987) in naïve and primed samples, along with Takashima et al samples. **(d-f)** Same as **a-c**, done over an independent RNA-seq dataset. Number of differentially expressed genes here is 7087. **g.** FACS analysis for expression levels of the indicated surface markers. CD130 and CD77 are induced in HENSM conditions consistent with their previous designation as markers of human naïve pluripotency (Collier et al. Cell Stem Cell 2017). CD24 is depleted in HENSM naïve conditions as expected.

**Figure S9. Expression profile of selected sets of genes in HENSM naïve and primed conditions.**

**a**. Expression profile of selected sets of genes in naïve and primed conditions. **b**. RT-PCR validation of expression of primed pluripotency markers ZIC2 and OTX2. Both were significantly depleted in HENSM based naïve conditions. *t-test *p* Value < 0.001. **c.** Strategy for generating human STELLA-CFP knock in reporter cell line. Both HENSM and HENSM-ACT conditions upregulated STELLA expression in comparison to primed cells and consistent with transcriptome data. **d**. FACS analysis for STELLA-CFP knock-in reporter expression levels in the indicated primed and naïve conditions. **e**. FACS staining results on different primed (red colors) and HENSM conditions for SUSD2 expression. MFI values are indicated and compared to matched unstained negative controls.

**Figure S10. Generation of TFAP2C KO human ESCS with reporter for human naive pluripotent state**.

**a**. Strategy for generating TFAP2C human KO in WIBR2 human ESC line carrying GFP/tdTomato reporters on each of the X chromosomes respectively. **b**. Immunostaining analysis for TFAP2C (also known as AP2gamma) expression in WT and KO human ESC clones (WIBR2-29-8 hESC line). OCT4 expression was not affected in primed KO cells in comparison to WT primed control cells. **c**. Strategy for generating TFAP2C human KO for TFAP2C via simultaneous targeting of both alleles in WIBR3 ΔPE-O4G hESCs. **d**. Western blot analysis for validation of TFAP2C KO generation in primed human WIBR3-ΔPE-O4G hESCs. **e.** Phase images showing loss of pluripotent cells expansion within 2 passages in both HENSM and HENSM-ACT conditions. **f.** RT-PCR analysis in naïve HENSM and primed conditions for naïve pluripotency genes and ESRRB. (UD-Undetectable). **g.** Targeting strategy for generating ESRRB-mCherry knock-in reporter human WIBR3 hESC line. **h**. PCR and southern blot analysis for detecting and selecting correctly targeted clones. **i.** FACS analysis for detecting ESRRB-mCherry expression in human PSCs. (Tg = Transgene). ESRRB-mCherry signal was detected only upon ectopic expression of ESRRB and KLF2 exogenous transgenes in addition to applying naïve HENSM conditions.

**Figure S11. Human PSCs in HENSM conditions have a transposon element (TE) transcription signature of the human pre-implantation embryo**.

**a**. Heatmap of RNA-seq expression data form primed human ESCs and naïve cells expanded in HENSM and HENSM-ACT conditions. Data shown include 10000 TEs with the highest standard deviation between samples. Figure shows clear separation between naïve HENSM and primed datasets in TE expression and profile. **b**. Principal component analysis (PCA) of primed (in TeSR or KSR/FGF2 conditions) or HENSM naïve conditions based on the differential expression of transposable elements (TEs). **c.** Correspondence between TE expression in HENSM naïve vs. primed ESCs and single-cell human embryonic stages (Yan et al., 2013). For every stage of human embryonic development, a statistical test was performed to find the TEs that have a different expression level compared to the other stages. The proportions of developmental stage-specific TEs that are upregulated (p < 0.05, 2-fold change) in naive or primed cells are indicated in orange and blue, respectively, while TEs that did not change expression are indicated in gray. The samples include all HENSM based naive cells that we examined in the PCA analyses in **b**.

**Figure S12. Rewiring TE elements in naïve HENSM conditions.**

Changes in distinct TE families are with indications whether they were downregulated or upregulated between HENSM naïve and primed conditions.

**Figure S13. Full analysis of LOI in HENSM and HENSM-ACT PSCs.**

Heatmap of biallelic expression including the complete list of imprinted genes in hPSC samples. Genes are arranged according to their genomic proximity. NE - not expressed (FPKM < 0.2).

**Figure S14. HENSM-derived human naïve pluripotent stem cells are competent for interspecies chimaera.**

**(a)** Representative images of whole mount *in toto* microscopy of chimaeric embryos over several developmental stages (e11.5 till e13.5). Panels on the right are zoomed-in regions of tiles on the left side. Hoechst was used as counterstain and anti-GFP staining used to trace human iPSC-derived descendants.

**(b)** Frozen tissue sections of E17.5 chimaeric embryos were stained for GFP and Human-Nuclei to confirm human origin identity. Non-injected embryos served as negative control. GFP, Human-Nuclei, overlap as well as merged are zoomed-in regions of lung tissue depicted in red squares in the tiles. White arrowheads in insets point out co-localization between GFP and Human-Nuclei. GFP, green fluorescent protein; WT, wild type; iPSC, induced pluripotent stem cell. Tile scale bar 500 um. Zoomed-in scale bar 100 um.

**Figure S15. HENSM-derived human naïve pluripotent stem cells integrate successfully into mouse embryos and acquire respective tissue identity**.

Representative images of frozen tissue sections of E17.5 chimaeric embryos were stained for GFP and Pro-Spc for lung-specific alveolar-surfactant secreting cells. Non-injected embryos served as negative control. **a.** GFP, Pro-Spc, overlap as well as merged are zoomed-in regions of lung tissue depicted in red squares in the tiles. White arrowheads in insets point out co-localization between GFP and Pro-Spc. GFP, green fluorescent protein; WT, wild type; Pro-Spc, prosurfactant Protein C; iPSC, induced pluripotent stem cell. Tile scale bar 1000 um. Zoomed-in scale bar 50 um. **b-c.** AS above but for Aqp5 (aquaporin 5) and CC10 (Clara-cell 10) lung cell markers.

**Figure S16. P53 (TP53) targeting in human iPSCs**.

**a.** Design of CRISPR/Cas9 targeting Exon 4 of hTP53 to generate knock-out with the guide RNA in red and the PAM sequence in green. **B.** Western blot analysis showing complete depletion of TP53 protein in various chosen clones. DNA sequence alignment showing out-of-frame insertions/deletions in clone C2 and a point mutation in clone E7. **c-d.** Staining and karyotyping showed normal pluripotency marker expression and karyotype in representative clones. **e.** NUMA staining of human vs mouse teratoma to confirm human specificity of the NUMA antibody used in this study to trace GFP-labelled human iPSCs after transplantation. NUMA, nuclear mitotic apparatus protein. Tile scale bar 1000 um. Zoomed-in region scale bar 100 um.

**Figure S17. Integration of P53 null naïve hiPSCs into mouse blastocysts.**

**a.** Phase and GFP image of P53 null naïve hiPSCs C8 clone expanded in HENSM-ACT. **b**. Phase and GFP images of mouse blastocysts 24h after microinjection into mouse morulas were conducted. **c**. Representative immunostaining images of mouse blastocysts 24h after GFP labeled human naïve P53 null hiPSCs were microinjected into mouse morulas. **d**. Representative immunostaining images of mouse blastocysts 48h after GFP labeled human naïve P53 null hiPSCs were micro-aggregated with mouse 2-Cell embryos.

**Figure S18. Boost in cell chimerism contribution by depleting P53 endows in hiPSCs before microinjection.**

**a.** Representative images depicting integration of P53KO GFP-labelled hiPSCs into different locations within developing E9.5/E10.5 mouse embryo in comparison to WT GFP+-cells and non-injected embryos used as negative controls. Red squares in the first column represent zoomed-in areas shown in the following images 1 and 2. Tile scale bar 200 um. Inset scale bar 50 um. **b.** Abnormally developed e10.5 chimaeric embryo with abundant GFP contribution in comparison to non-injected WT embryo lacking any GFP signature. Hoechst and CellTracker were used for counterstaining. Scale bar 50 um. **c.** Representative images of whole-mount *in toto* imaged e15.5 mouse embryos are shown in comparison to non-injected wild-type embryos. White squares in tiles outline zoomed-in regions in subsequent panels. GFP staining was used to trace hiPSC-derived progeny and CellTracker and Hoechst as counterstaining. Scale bar 1 mm. GFP, green fluorescent protein; p53, tumor protein p53; iPSC, induced pluripotent stem cell; WT, wild type.

**Figure S19. Naïve hiPSCs contribution to chimaeric mouse embryos.**

**a.** FACS analysis for GFP detection in mouse embryos. Non-injected embryos and a parental GFP+ hiPSC line were used as controls. **b.** Human mitochondrial specific DNA detection assay for dilutions between human and mouse cells, chimaeric embryos, non-injected embryos as well as for different pure mouse tissues. Dotted line represents threshold level set in this PCR. Green arrows highlight mouse chimeric embryos with high levels of human DNA detection. **c-e.** IHC staining of various regions of chimaeric embryos for NUMA to trace GFP+ human P53KO hIPSC derived cells. GFP, NUMA, overlap as well as merged are zoomed-in regions of lung tissue depicted in red squares in the tiles. White arrowheads in insets point out co-localization between GFP and NUMA. Non-injected embryos served as negative control. Tile scale bar 2000µm. Zoomed-in region scale bar 100µm.

**Figure S20. Naïve hiPSC-derived cells colonize various anatomical regions throughout whole mouse embryo.**

**a-d.** IHC staining of various regions of chimaeric embryos for NUMA to trace P53KO hIPSC derived GFP+ human cells. GFP, NUMA, overlap as well as merged are zoomed-in regions of lung tissue depicted in red squares in the tiles. White arrowheads in insets point out co-localization between GFP and NUMA. Non-injected embryos served as negative control. Tile scale bar 1000 um and 2000 um. Zoomed-in region scale bar 100µm.

**Figure S21. Contribution of GFP+ hiPSC-derived progeny to ectodermal-neural lineages within chimaeric mouse embryos.**

**a-b.** Representative images of IHC staining of injected (upper panels) and non-injected E15.5 mouse embryos (lower panels) for (a) brain TUJ1 and (b) cochlea SOX2 respectively are shown. GFP served as human cell tracer of microinjected P53KO hiPSC, and TUJ1 and SOX2 as neural progenitor signature. GFP, TUJ1 or SOX2, overlap as well as merged constitute zoomed-in regions of tissues depicted in red squares in the tiles. White arrowheads in insets depict co-localization of GFP and TUJ1. GFP, green fluorescent protein; TUJ1, neuron-specific class III beta-tubulin; SOX2, sex determining region Y-box; P53KO, knock-out of tumor protein p53. Tile scale bar 2000 um. Non-injected scale bar 1 mm. Zoomed-in region scale bar 100µm.

**Figure S22. Contribution of GFP+ hiPSC-derived progeny to ectodermal-neural lineages within chimaeric mouse embryos.**

**a-c.** Representative images of IHC for TUJ1 and GFP of injected (upper panels) and non-injected E15.5 mouse embryos (lower panels) for each tissue type (a,b,c) respectively are shown. GFP served as human cell tracer of microinjected P53KO hiPSC. GFP, TUJ1, overlap as well as merged constitute zoomed-in regions of tissues depicted in red squares in the tiles. White arrowheads in insets depict co-localization of GFP and TUJ1. GFP, green fluorescent protein; TUJ1, neuron-specific class III beta-tubulin; P53KO, knock-out of tumor protein p53. Tile scale bar 2000µm. Zoomed-in region scale bar 100µm.

**Figure S23. Contribution of GFP+ hiPSC-derived progeny to ectodermal-neural lineages within chimaeric mouse embryos.**

**a-d.** Representative images of IHC for SOX2 of injected (upper panels) and non-injected E15.5 mouse embryos (lower panels) for each tissue type respectively are shown. GFP served as human cell tracer of microinjected P53KO hiPSC, and SOX2 as neural progenitor signature. GFP, SOX2, overlap as well as merged constitute zoomed-in regions of tissues depicted in red squares in the tiles. White arrowheads in insets depict co-localization of GFP and SOX2. GFP, green fluorescent protein; SOX2, sex determining region Y-box; P53KO, knock-out of tumor protein p53. Tile scale bar 1000µm. Zoomed-in region scale bar 100µm.

**Figure S24. Contribution of GFP+ hiPSC-derived progeny to endoderm and mesoderm lineages within chimaeric mouse embryos.**

**a-d.** Representative images of IHC for SOX17 of injected (upper panels) and non-injected E15.5 mouse embryos (lower panels) for each tissue type respectively are shown. GFP served as human cell tracer of microinjected P53KO hiPSC and SOX17 as endoderm progenitor and mesoderm-progeny tissue marker. GFP, SOX17, overlap as well as merged constitute zoomed-in regions of tissues depicted in red squares in the tiles. White arrowheads in insets depict co-localization of GFP and SOX17. GFP, green fluorescent protein; SOX17, SRY-related HMG-box 17; P53KO, knock-out of tumor protein p53. Tile scale bar 2000µm. Zoomed-in region scale bar 100µm.

**Figure S25. Contribution of GFP+ hiPSC-derived progeny to endoderm and mesodermal lineages within chimaeric mouse embryos.**

**a-b.** Representative images of IHC for SOX17 of injected (upper panels) and non-injected E15.5 mouse embryos (lower panels) for each tissue type respectively are shown. GFP served as human cell tracer of microinjected P53KO hiPSC and SOX17 as endoderm progenitor and mesoderm-progeny tissue marker. GFP, SOX17, overlap as well as merged constitute zoomed-in regions of tissues depicted in red squares in the tiles. White arrowheads in insets depict co-localization of GFP and SOX17. GFP, green fluorescent protein; SOX17, SRY-related HMG-box 17; P53KO, knock-out of tumor protein p53. Tile scale bar 2000 um. Zoomed-in region scale bar 100 um. **c.** Summarizing table for outcome of injections of different naïve and primed hiPSCs into mouse embryos.

**Figure S26. Nuclear** β**CATENIN signaling induces priming of human PSCs.**

**a**. FACS analysis for OCT4-GFP reporter in HENSM conditions before and after depletion of indicated media components. **b**. Mouse and human βCATENIN null ESCs were made transgenic for an exogenous validated tamoxifen inducible βCATENIN transgene (βCATKO-βCateninERT-Tg). Immunostaining confirms increase in nuclear βCATENIN upon tamoxifen addition (4OHT) in mouse and human validated clones used for analysis. **c**. RT-PCR analysis for naïve pluripotency maker expression in Human WIBR3-ΔPE-βCATKO; βCateninERT-Tg line before and after Tamoxifen addition for 48 hours. Values were normalized to ACTIN and GAPDH. Primed expression levels were set as 1. *t-test *p* Value < 0.01. Naïve pluripotency marker expression where significantly downregulating upon induction of nuclear βCATENIN signaling in human PSCs.

**Figure S27. Generation of** β**CATENIN knockout hESCs with pluripotency reporters.**

**a**. Scheme depicts CRISPR/Cas9 based strategy for generating βCATENIN KO in WIBR3-OCT4-GFP and WIBR3-ΔPE-OCT4-GFP primed human ESCs. **b.** Sequencing results and **c**. western blot validation for βCATENIN in correctly targeted WIBR3-OCT4-GFP clones. **d.** Sequencing results and **e**. western blot validation for βCATENIN in correctly targeted WIBR3-ΔPE-OCT4-GFP clones.

**Figure S28. Ablating TCF3 does not support human naïve pluripotency.**

**a.** Scheme depicts strategy for TCF3 knockout induction in human WIBR3-OCT4-GFP ESC. **b.** Western blot analysis validates TCF3 protein deletion in candidate correctly targeted clones**. c.** Phase contrast images of TCF3^-/-^ ESCs before and after TNKi removal from HENSM conditions. **d**. RT-PCR analysis for naïve pluripotency maker expression in the indicated conditions, with and without TNKi, in TCF3 WT and KO hESCs. Values were normalized to ACTIN and GAPDH. *t-test *p* Value < 0.01. TCF3^+/+^ HENSM expression levels were set as 1. **e**. Percentage of OCT4+ cells at P4 from TET-OFF-DNMT1, DNMT1-TET-OFF cell lines and OCT4-GFP cells in the absence of exogenous L-Glut in primed, naïve HENSM, and naïve HENSM supplemented with 1.5µM CHIR99021. **f.** Normalized expression pattern of TCF3 and AXIN1/2 in mouse and human, as reported in previous single-cell RNA-seq measurement in mouse and human embryos.

**Figure S29. LIF-STAT3 signaling supports human naïve pluripotency but overall can be dispensable. a.** RT-PCR analysis for naïve pluripotency maker expression in the indicated naïve and primed conditions, with and without LIF. Values were normalized to ACTIN and GAPDH. Primed expression levels were set as 1. *t-test *p* Value < 0.01; NS-not significant. **b.** Scheme depicts strategy for generating STAT3 knockout WIBR3-OCT4-GFP hESCs. **c**. Western blot validation for loss of STAT3 protein in correctly targeted clones that were validated by southern blot analysis. **d.** RT-PCR analysis for naïve pluripotency maker expression in the indicated lines and conditions. Values were normalized to ACTIN and GAPDH. Primed expression levels were set as 1. *t-test *p* Value < 0.01; NS-not significant. Naïve pluripotency marker remain highly expressed in STAT3 KO ESCs in HENSM conditions, when compared to primed cells that do not express at all these specific markers.

**Figure S30. Generation of KLF4 and/or KLF17 knockout human ESCs.**

**a.** Scheme depicts strategy for generating KLF17 null WIBR3-OCT4-GFP hESCs. **b**. Southern blot analysis validating correctly targeted clones for KLF17 KO. **c**. Immunostaining validation for loss of KLF17 protein in early passage KO clone expanded for 2 passages in HENMS-ACT. **d.** Scheme depicts strategy for generating KLF4 knockout WIBR3-OCT4-GFP hESCs. **e**. Southern blot analysis validating correctly targeted clones for KLF4 KO. **f**. Immunostaining validation for loss of KLF4 protein in early passage KO clone expanded for 2 passages in HENMS-ACT. Same strategy for making KLF4 knockout (**d-f**) was applied on KLF17^-/-^ cells to generate double knockout validated clones.

**Figure S31. Klf17 is dispensable for mouse naïve pluripotency *in vivo* and *in vitro*.**

**a.** KLF4 and KLF17 relative abundance in pre-implantation stages as measured in vivo by scRNA-seq. Please note that KLF17 is not expressed in the mouse ICM (but rather a 2 Cell stage in the mouse), while in humans it is rather upregulated at the morula-ICM stages. **b.** Scheme depicts strategy for generating Klf17 null allele in V6.5 mouse ESCs. **c**. PCR analysis for checking correctly targeted clones. **d**. Southern blot analysis validating correctly targeted clones. **e**. PCR analysis for confirming correct flipping out of antibiotic resistance cassette prior to ESC microinjection. **f**. Germ line transmission from Klf17 targeted mouse ESCs as confirmed by obtaining agouti colored pups. **g**. RT-PCR analysis for naïve pluripotency markers in WT and Klf17^-/-^ mouse ESCs expanded in 2i/LIF conditions. Values per each gene were normalized in WT ESCs as 1. Klf17^-/-^ did not show any significant change in naïve pluripotency marker expression. (NS – not significant).

**Figure S32. Alternative HENSM conditions for human naïve PSCs without using ERKi.**

**a.** Spearman correlation matrix of primed samples, along HENSM, HENSM-ACT, tHENSM and 0HENSM samples, as well as previously published naïve and primed samples (Takashima et al, 2014). **b-d**. Percentage of OCT4-GFP+ cells at P4 from WIBR3-OCT4-GFP cells in HENSM, tHENSM and 0HENSM conditions without METTL3 (**b**), DNMT1 (**c**) or exogenously added L-Glutamine (**d**). Note that removal of DBZ from tHENSM or 0HENSM results in loss of maintenance of pluripotency when these factors are depleted. **e**. Competence for human naïve PSCs for differentiating into PGCLCs in vitro was assays for NANOS3-mCherry knock in reporter line expanded for at least 3 passages in HENSM, tHENSM or 0HENSM conditions. Percentage of NANOS3+ cells detected by FACS analysis are shown.

**Figure S33. Pattern of DNA methylation regulators in alternative tHENSM and OHENSM conditions.**

**a**. Immunostaining for DNMT3L in primed and different naïve conditions are shown. DNMT3L levels are reduced in tHENSM and 0HENSM conditions as seen in primed conditions. **b**. Global methylation histogram calculated from primed samples, and naïve samples that were maintained in various conditions, including titrated ERKi supplementation, along with previously published Reset-naïve* and primed* samples (Takashima et al. Cell 2014), and human ICM samples. DNMT1^-/-^ (from TET-OFF lines) samples were used as negative control for methylation. Dark blue - percentage of highly methylated CpGs (>0.9 methylation level), light blue – percentage of lowly methylated CpGs (<0.1 methylation level). Yellow dots – sample methylation average. **c.** Average methylation as calculated from primed samples, and naïve samples that were maintained in various conditions, including titrated ERKi (tHENSM and 0HENSM) supplementation.

**Figure S34. Generation and validation of RBPj knockout human ESCs. a**. Scheme depicts targeting strategy for generating knockout human WIBR3 hESCs for RBPj. **b.** Correctly targeted clones validated by southern blot analysis were confirmed for loss of RBPj protein by western blot analysis.

## Supplementary Table Legends

**Table S1.** Diferentially expressed genes between naive and primed samples (Fold change >2, adjusted p-value <0.1) in dataset 1, along with normalized counts of all genes. Related to Fig. 3 & S8.

**Table S2.** Normalized counts of all genes in Dataset 2 (HENSM naive, primed, tHENSM naïve, 0HENSM naïve samples). Related to Fig. 7, S8, S9 & S32.

**Table S3.** Differentially expressed transposable elements, based on dataset 1. Related to Fig. S11-S12.

**Table S4.** Significant binding motifs identified by Homer software, calculated from the targets of KLF4, KLF17, TFAP2C, SOX2, OCT4, NANOG. Related to Fig. 6.

**Table S5.** Gene targets and sample statistics of ChIP-seq experiments of KLF4, KLF17, TFAP2C, SOX2, OCT4, NANOG in HENSM and Primed conditions. Related to Fig. 6.

