## Supplementary Figures S1-S34 for "Tripartite Inhibition of SRC-WNT-PKC Signalling Consolidates Human Naïve Pluripotency"

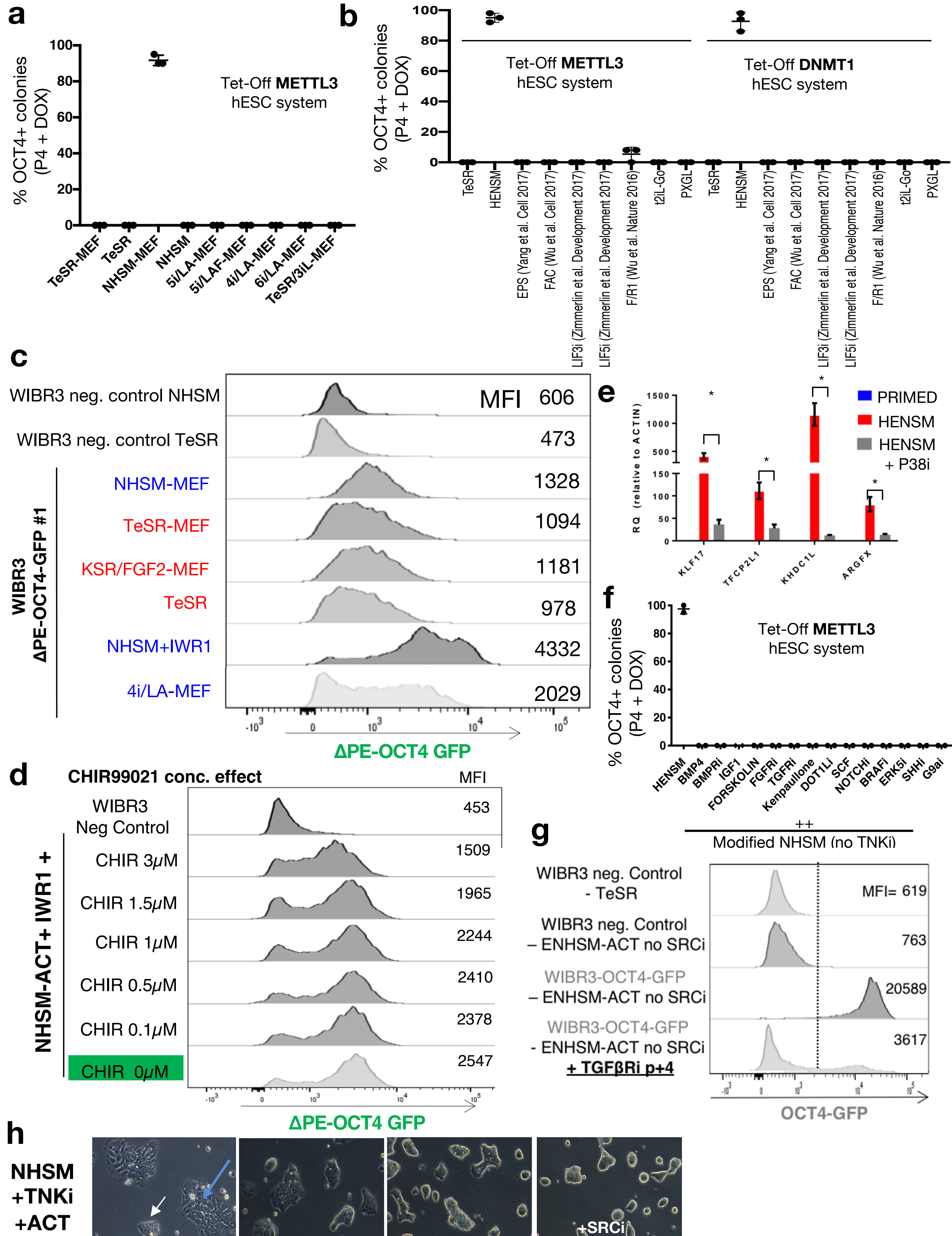
a
% OCT4+ colonies (P4 + DOX)
Tet-Off METTL3 hESC system
TeSR-MEF
TeSR
NHSM-MEF
NHSM
5i/LA-MEF
5i/LAF-MEF
4i/LA-MEF
6i/LA-MEF
TeSR/3iL-MEF
b
% OCT4+ colonies (P4 + DOX)
Tet-Off METTL3 hESC system
Tet-Off DNMT1 hESC system
TeSR
HENSM
EPS (Yang et al. Cell 2017)
FAC (Wu et al. Cell 2017)
LIF3i (Zimmerlin et al. Development 2017)
LIF5i (Zimmerlin et al. Development 2017)
F/R1 (Wu et al. Nature 2016)
t2iL-Go
PXGL
c
WIBR3 neg. control NHSM
WIBR3 neg. control TeSR
WIBR3 ΔPE-OCT4-GFP #1
NHSM-MEF
TeSR-MEF
KSR/FGF2-MEF
TeSR
NHSM+IWR1
4i/LA-MEF
MFI 606
473
1328
1094
1181
978
4332
2029
ΔPE-OCT4 GFP
d
CHIR99021 conc. effect
MFI
WIBR3 Neg Control
NHSM-ACT+ IWR1 +
CHIR 3μM
CHIR 1.5μM
CHIR 1μM
CHIR 0.5μM
CHIR 0.1μM
CHIR 0μM
453
1509
1965
2244
2410
2378
2547
ΔPE-OCT4 GFP
e
RQ (relative to ACTIN)
PRIMED
HENSM
HENSM + P38i
KLF17
TFCP2L1
KHDC1L
ARGFX
f
% OCT4+ colonies (P4 + DOX)
Tet-Off METTL3 hESC system
HENSM
BMP4
BMPRi
IGF1
FORSKOLIN
FGFRi
TGFRi
Kenpaullone
DOT1Li
SCF
NOTCHi
BRAFi
ERK5i
SHHi
G9ai
g
++
Modified NHSM (no TNKi)
WIBR3 neg. Control - TeSR
WIBR3 neg. Control – ENHSM-ACT no SRCi
WIBR3-OCT4-GFP – ENHSM-ACT no SRCi
WIBR3-OCT4-GFP - ENHSM-ACT no SRCi + TGFβRi p+4
MFI= 619
763
20589
3617
OCT4-GFP
h
NHSM +TNKi +ACT
Control
100μm
+SRCi
+ 0.2% Geltrex
+SRCi + 0.2% Geltrex

Figure S1

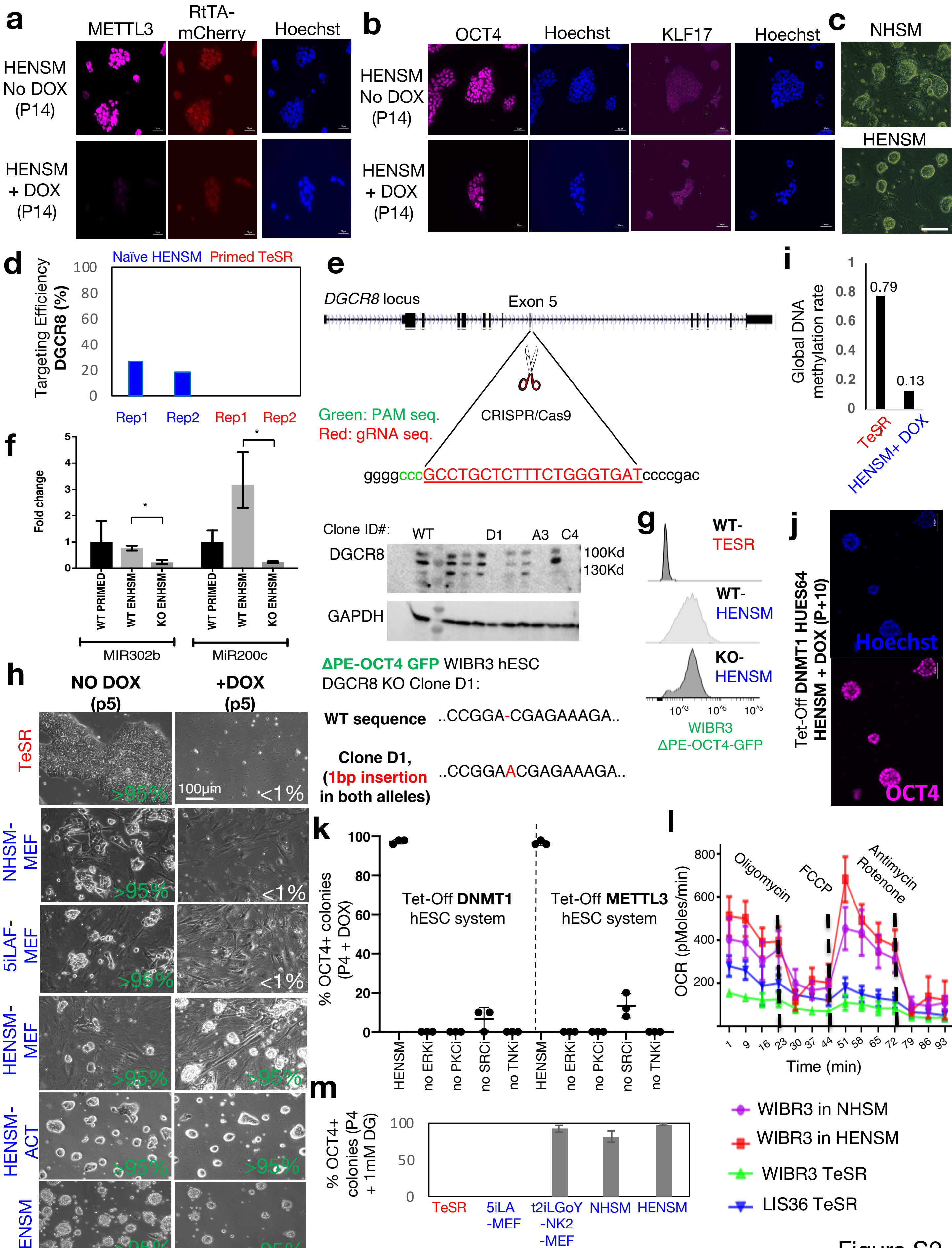

a
METTL3
RtTA-mCherry
Hoechst
HENSM No DOX (P14)
HENSM + DOX (P14)
b
OCT4
Hoechst
KLF17
Hoechst
HENSM No DOX (P14)
HENSM + DOX (P14)
c
NHSM
HENSM
d
Naïve HENSM
Primed TeSR
Targeting Efficiency DGCR8 (%)
100
80
60
40
20
0
Rep1
Rep2
Rep1
Rep2
e
DGCR8 locus
Exon 5
CRISPR/Cas9
Green: PAM seq.
Red: gRNA seq.
ggggcccGCCTGCTCTTTCTGGGTGATccccgac
Clone ID#:
WT
D1
A3
C4
DGCR8
100Kd
130Kd
GAPDH
ΔPE-OCT4 GFP WIBR3 hESC
DGCR8 KO Clone D1:
WT sequence
..CCGGA-CGAGAAAGA..
Clone D1, (1bp insertion in both alleles)
..CCGGAACGAGAAAGA..
i
Global DNA methylation rate
1
0.8
0.6
0.4
0.2
0
0.79
0.13
TeSR
HENSM+ DOX
f
Fold change
5
4
3
2
1
0
*
*
WT PRIMED
WT ENHSM
KO ENHSM
WT PRIMED
WT ENHSM
KO ENHSM
MIR302b
MiR200c
g
WT-
TESR
WT-
HENSM
KO-
HENSM
10^3
10^5
10^5
WIBR3
ΔPE-OCT4-GFP
j
Tet-Off DNMT1 HUES64 HENSM + DOX (P+10)
Hoechst
OCT4
h
NO DOX (p5)
+DOX (p5)
TeSR
>95%
100μm
<1%
NHSM-MEF
>95%
<1%
5iLAF-MEF
>95%
<1%
HENSM-MEF
>95%
>95%
HENSM-ACT
>95%
>95%
HENSM
>95%
>95%
k
% OCT4+ colonies (P4 + DOX)
100
80
60
40
20
0
Tet-Off DNMT1 hESC system
Tet-Off METTL3 hESC system
HENSM
no ERKi
no PKCi
no SRCi
no TNKi
HENSM
no ERKi
no PKCi
no SRCi
no TNKi
l
OCR (pMoles/min)
800
600
400
200
Oligomycin
FCCP
Antimycin Rotenone
1
9
16
23
30
37
44
51
58
65
72
79
86
93
Time (min)
WIBR3 in NHSM
WIBR3 in HENSM
WIBR3 TeSR
LIS36 TeSR
m
% OCT4+ colonies (P4 + 1mM DG)
100
50
0
TeSR
5iLA-MEF
t2iLGoY-NK2-MEF
NHSM
HENSM


Figure S2

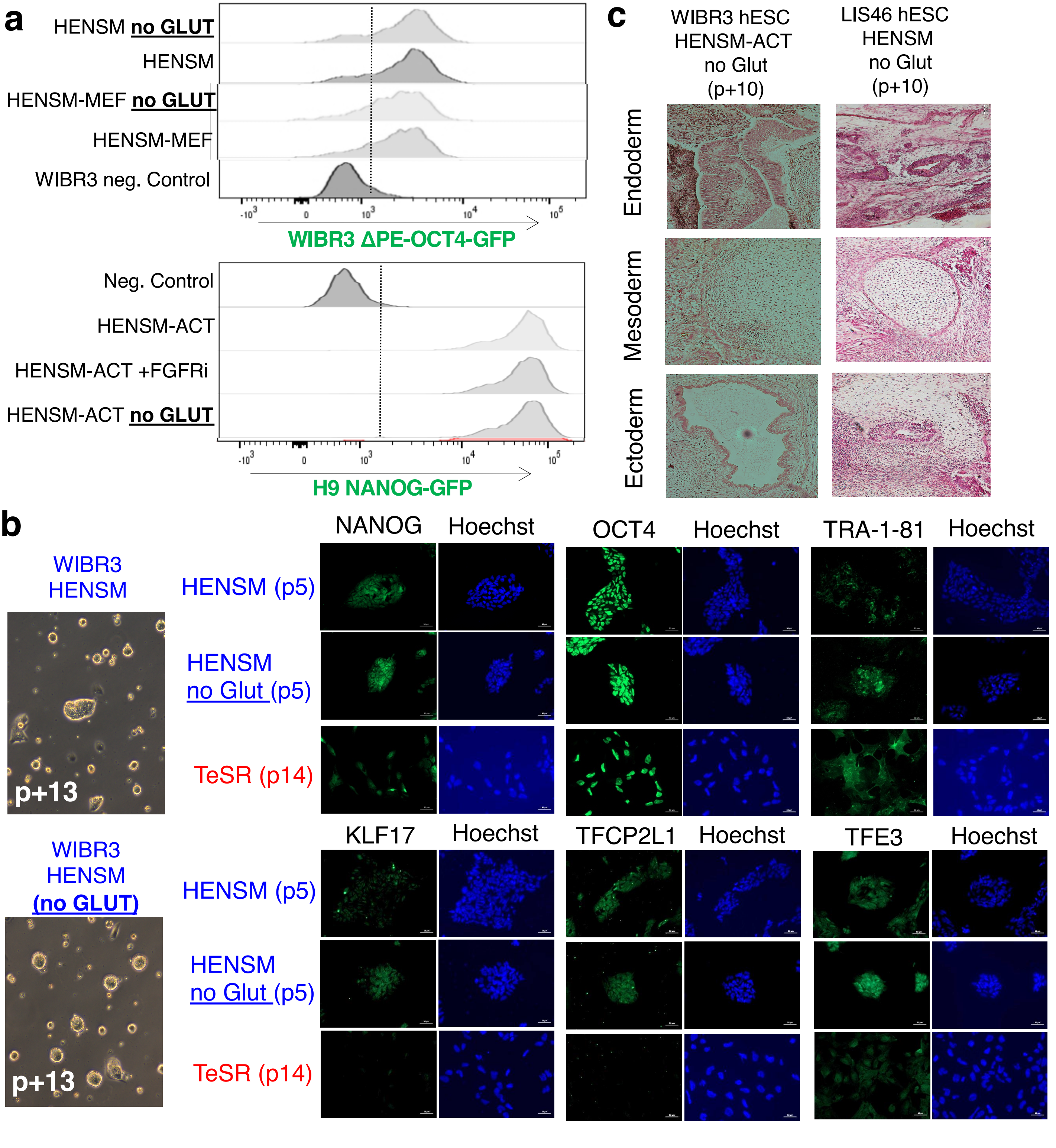
a
HENSM no GLUT
HENSM
HENSM-MEF no GLUT
HENSM-MEF
WIBR3 neg. Control
WIBR3 ΔPE-OCT4-GFP
Neg. Control
HENSM-ACT
HENSM-ACT +FGFRi
HENSM-ACT no GLUT
H9 NANOG-GFP
c
WIBR3 hESC HENSM-ACT no Glut (p+10)
LIS46 hESC HENSM no Glut (p+10)
Endoderm
Mesoderm
Ectoderm
b
WIBR3 HENSM
p+13
WIBR3 HENSM (no GLUT)
p+13
NANOG
Hoechst
OCT4
Hoechst
TRA-1-81
Hoechst
HENSM (p5)
HENSM no Glut (p5)
TeSR (p14)
KLF17
Hoechst
TFCP2L1
Hoechst
TFE3
Hoechst
HENSM (p5)
HENSM no Glut (p5)
TeSR (p14)

Figure S3

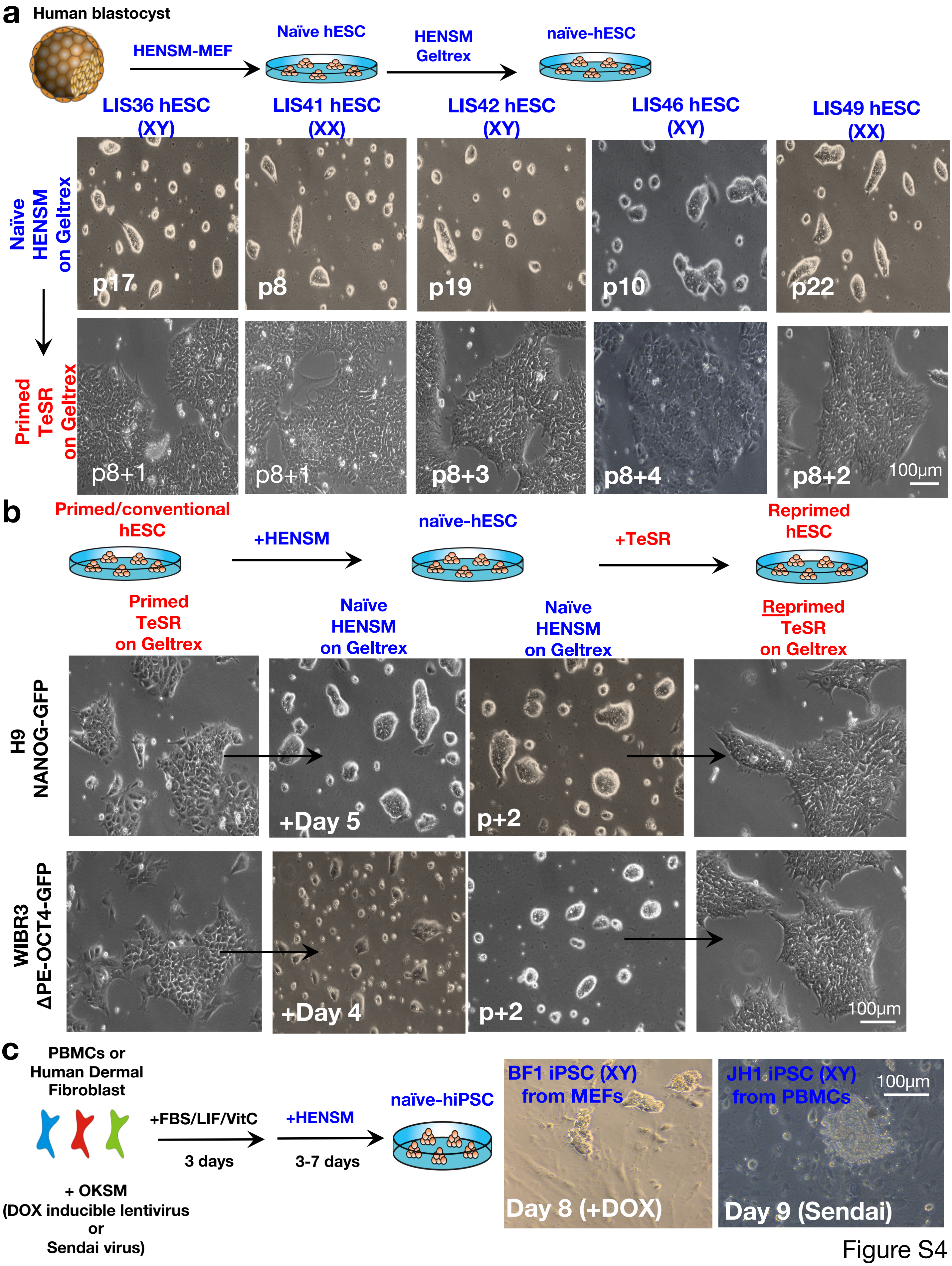
a
Human blastocyst
HENSM-MEF
Naïve hESC
HENSM
Geltrex
naïve-hESC
LIS36 hESC (XY)
LIS41 hESC (XX)
LIS42 hESC (XY)
LIS46 hESC (XY)
LIS49 hESC (XX)
Naïve HENSM on Geltrex
p17
p8
p19
p10
p22
Primed TeSR on Geltrex
p8+1
p8+1
p8+3
p8+4
p8+2
100μm
b
Primed/conventional hESC
+HENSM
naïve-hESC
+TeSR
Reprimed hESC
Primed TeSR on Geltrex
Naïve HENSM on Geltrex
Naïve HENSM on Geltrex
Reprimed TeSR on Geltrex
H9 NANOG-GFP
+Day 5
p+2
WIBR3 ΔPE-OCT4-GFP
+Day 4
p+2
100μm
c
PBMCs or Human Dermal Fibroblast
+FBS/LIF/VitC
3 days
+HENSM
3-7 days
naïve-hiPSC
+ OKSM
(DOX inducible lentivirus or Sendai virus)
BF1 iPSC (XY) from MEFs
Day 8 (+DOX)
JH1 iPSC (XY) from PBMCs
100μm
Day 9 (Sendai)

Figure S4

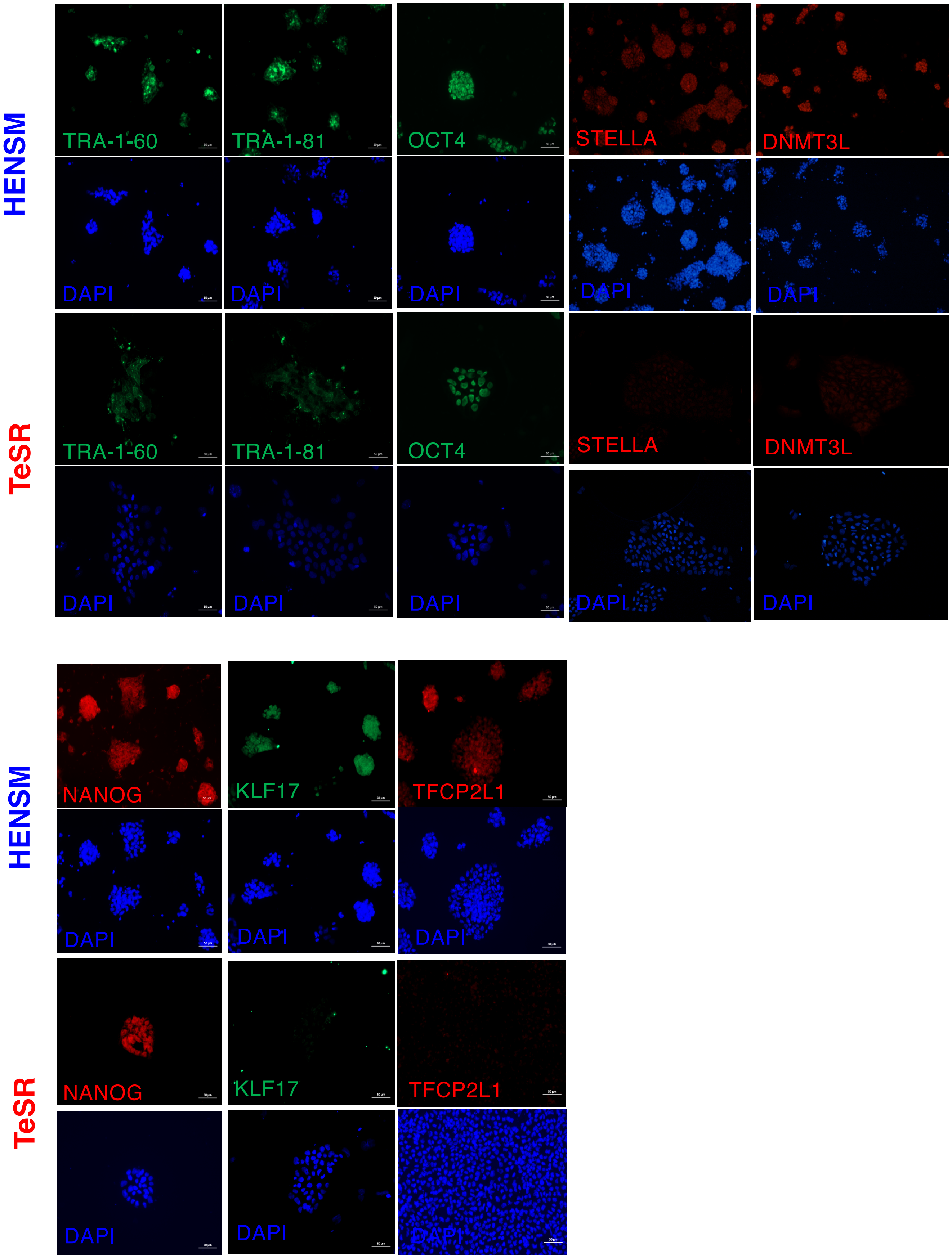
HENSM
TeSR
TRA-1-60
TRA-1-81
OCT4
STELLA
DNMT3L
DAPI
NANOG
KLF17
TFCP2L1

Figure S5

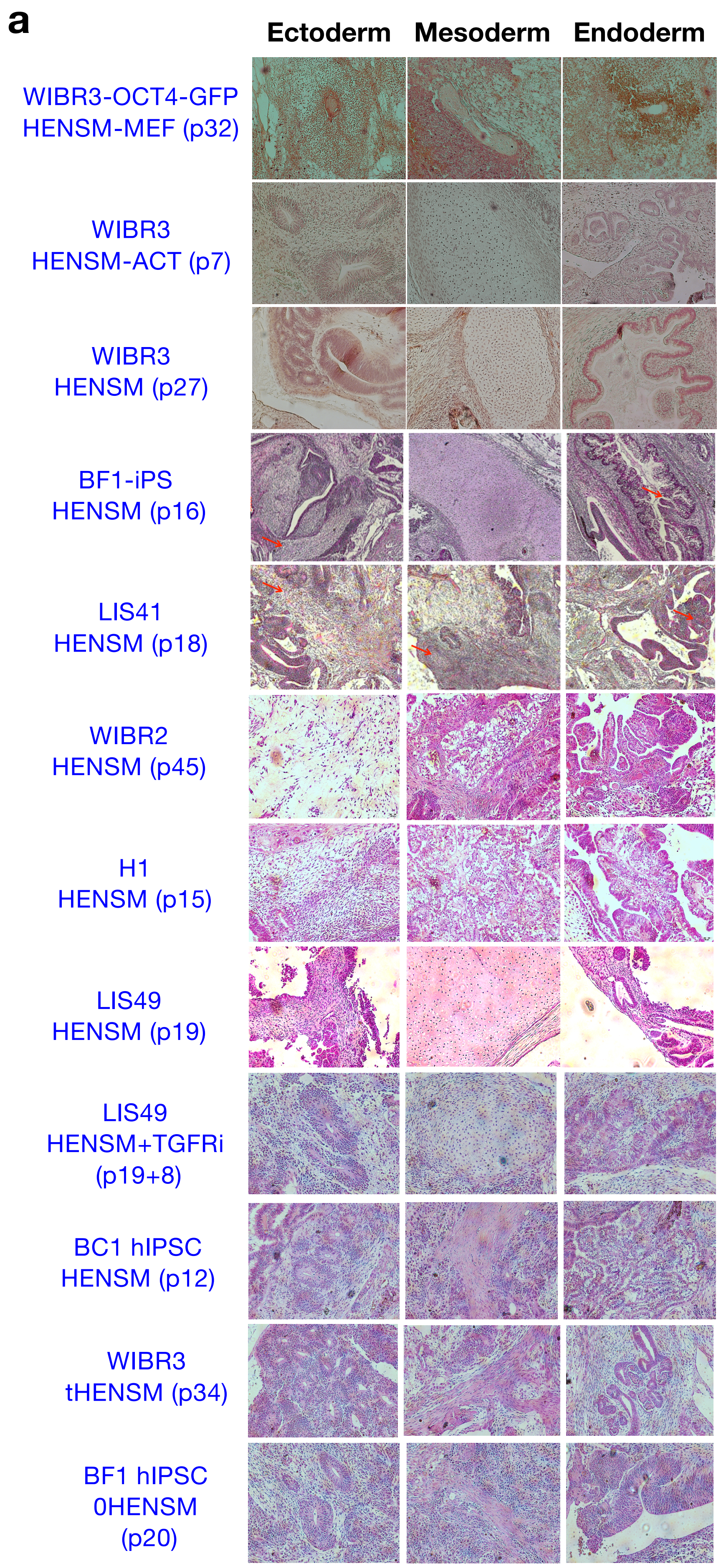


**b**

| Cell Line | Type | Karyotype |
|---|---|---|
| H1 | Primed (P54) | Normal |
| | Naïve HENSM-ACT (P54+40) | Duplication of Chr17 |
| HuES64 | Naïve HENSM (P+30) | Normal |
| LIS41 | RePrimed (P20+20) | Normal |
| | Naïve HENSM (P40) | Normal |
| LIS49 | Naïve HENSM-ACT (P20) | Duplication in Chr1 & deletion in Chr10 |
| | Naïve HENSM (P45) | Normal |
| WIBR3 | Naïve HENSM (P+25) and HENSM-ACT (P+31) | Normal |
| WIBR2 | Primed (P22) | Normal |
| | HENSM-ACT (P22+33) | Normal |

Figure S6

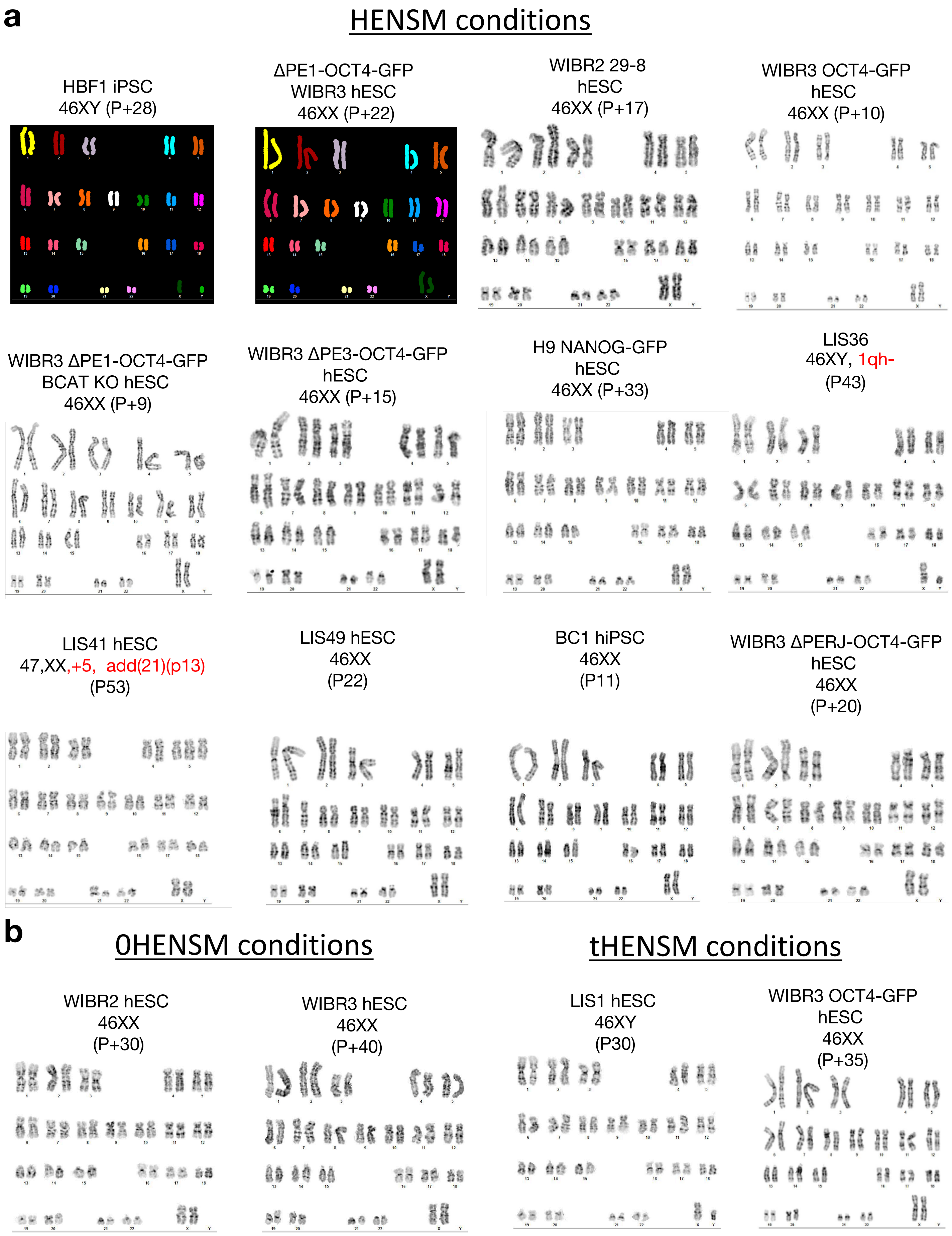
a
HENSM conditions
HBF1 iPSC
46XY (P+28)
ΔPE1-OCT4-GFP
WIBR3 hESC
46XX (P+22)
WIBR2 29-8
hESC
46XX (P+17)
WIBR3 OCT4-GFP
hESC
46XX (P+10)
WIBR3 ΔPE1-OCT4-GFP
BCAT KO hESC
46XX (P+9)
WIBR3 ΔPE3-OCT4-GFP
hESC
46XX (P+15)
H9 NANOG-GFP
hESC
46XX (P+33)
LIS36
46XY, 1qh-
(P43)
LIS41 hESC
47,XX,+5, add(21)(p13)
(P53)
LIS49 hESC
46XX
(P22)
BC1 hiPSC
46XX
(P11)
WIBR3 ΔPERJ-OCT4-GFP
hESC
46XX
(P+20)
b
0HENSM conditions
tHENSM conditions
WIBR2 hESC
46XX
(P+30)
WIBR3 hESC
46XX
(P+40)
LIS1 hESC
46XY
(P30)
WIBR3 OCT4-GFP
hESC
46XX
(P+35)

Figure S7

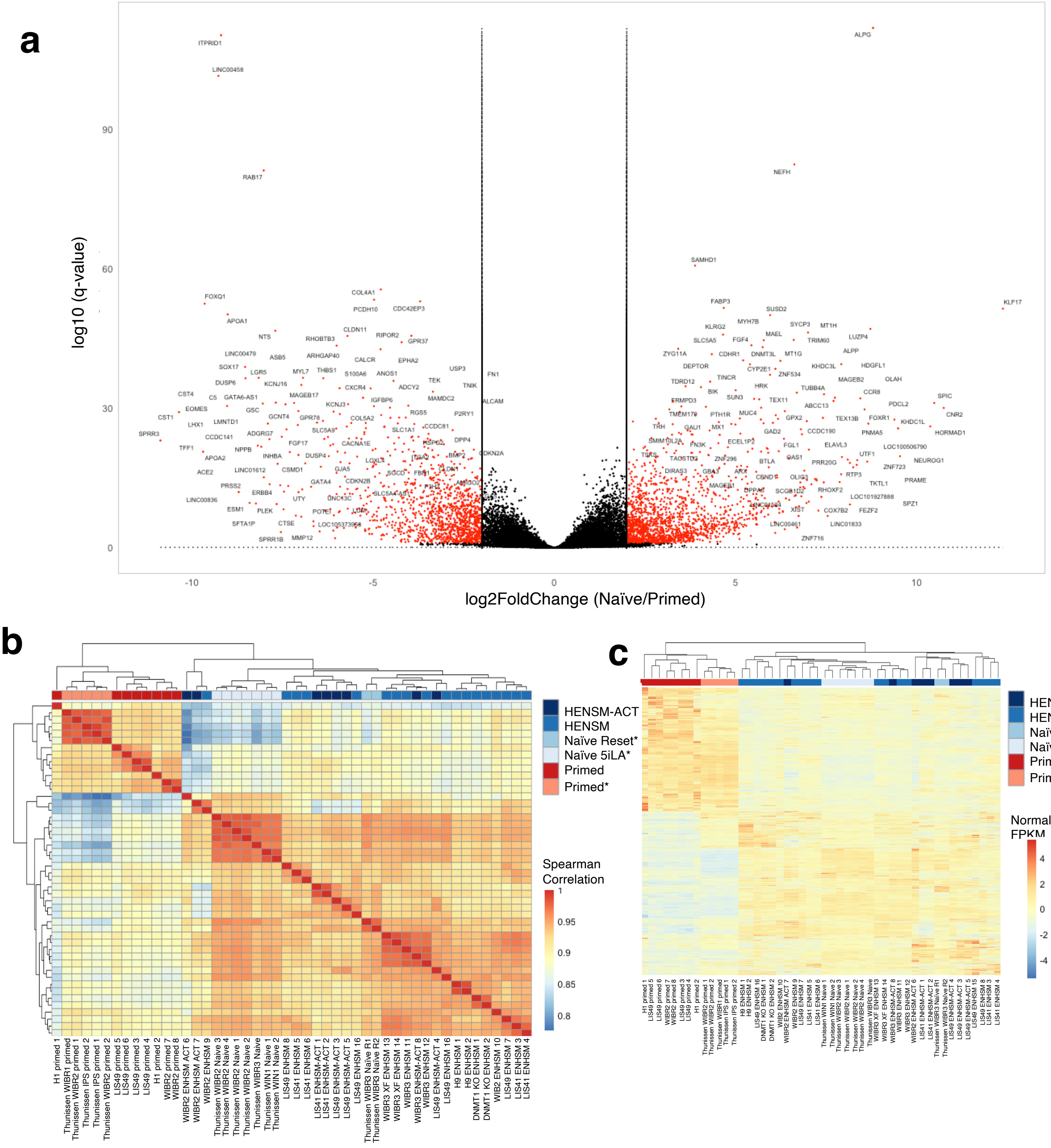
a
log10 (q-value)
log2FoldChange (Naïve/Primed)
-10
-5
0
5
10
90
60
30
0
b
HENSM-ACT
HENSM
Naïve Reset*
Naïve 5iLA*
Primed
Primed*
Spearman
Correlation
1
0.95
0.9
0.85
0.8
c
HENSM-ACT
HENSM
Naïve Reset*
Naïve 5iLA*
Primed
Primed*
Normalized
FPKM
4
2
0
-2
-4

Figure S8

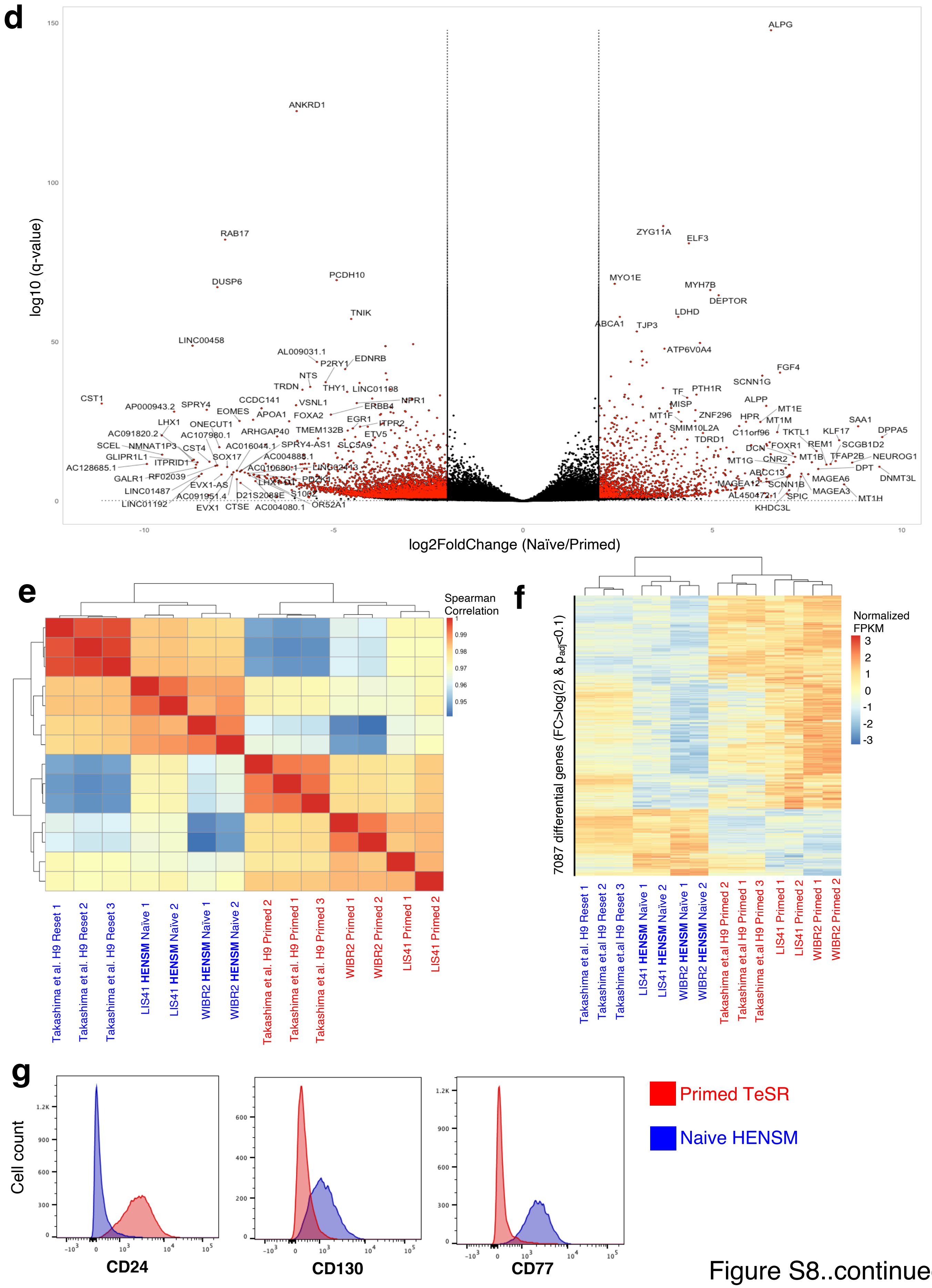


Figure S8..continued

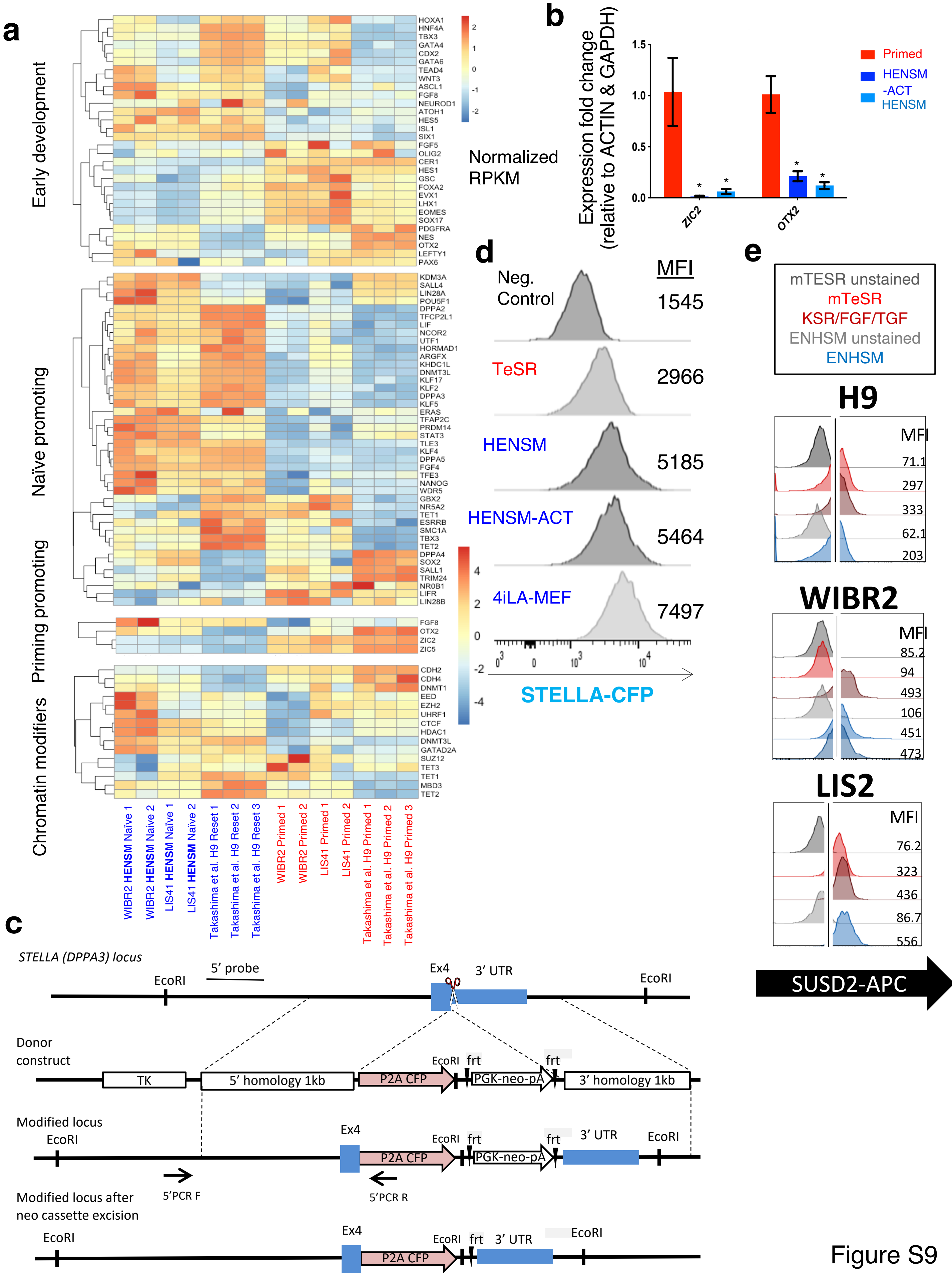
a
Early development
Naïve promoting
Priming promoting
Chromatin modifiers
Normalized RPKM
WIBR2 HENSM Naïve 1
WIBR2 HENSM Naïve 2
LIS41 HENSM Naïve 1
LIS41 HENSM Naïve 2
Takashima et al. H9 Reset 1
Takashima et al. H9 Reset 2
Takashima et al. H9 Reset 3
WIBR2 Primed 1
WIBR2 Primed 2
LIS41 Primed 1
LIS41 Primed 2
Takashima et al. H9 Primed 1
Takashima et al. H9 Primed 2
Takashima et al. H9 Primed 3
b
Expression fold change (relative to ACTIN & GAPDH)
Primed
HENSM
-ACT HENSM
ZIC2
OTX2
c
STELLA (DPPA3) locus
5' probe
EcoRI
Ex4
3' UTR
Donor construct
TK
5' homology 1kb
P2A CFP
PGK-neo-pA
3' homology 1kb
frt
Modified locus
5'PCR F
5'PCR R
Modified locus after neo cassette excision
d
Neg. Control
MFI
1545
TeSR
2966
HENSM
5185
HENSM-ACT
5464
4iLA-MEF
7497
STELLA-CFP
e
mTESR unstained
mTeSR
KSR/FGF/TGF
ENHSM unstained
ENHSM
H9
MFI
71.1
297
333
62.1
203
WIBR2
85.2
94
493
106
451
473
LIS2
76.2
323
436
86.7
556
SUSD2-APC

Figure S9

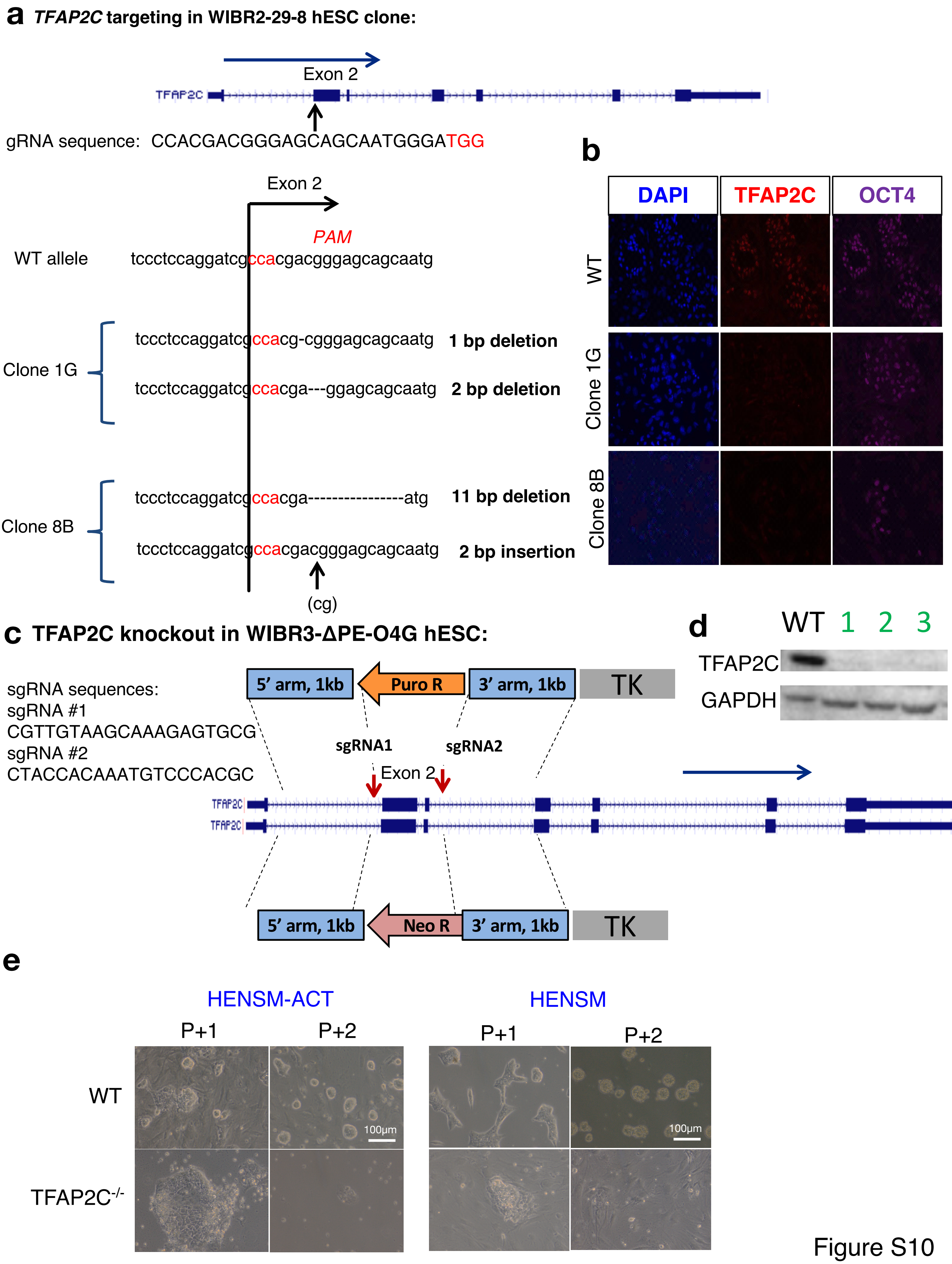
a TFAP2C targeting in WIBR2-29-8 hESC clone:
Exon 2
TFAP2C
gRNA sequence: CCACGACGGGAGCAGCAATGGGATGG
Exon 2
PAM
WT allele tccctccaggatcgccacgacgggagcagcaatg
Clone 1G
tccctccaggatcgccacg-cgggagcagcaatg 1 bp deletion
tccctccaggatcgccacga---ggagcagcaatg 2 bp deletion
Clone 8B
tccctccaggatcgccacga----------------atg 11 bp deletion
tccctccaggatcgccacgacgggagcagcaatg 2 bp insertion
(cg)
b
DAPI
TFAP2C
OCT4
WT
Clone 1G
Clone 8B
c TFAP2C knockout in WIBR3-ΔPE-O4G hESC:
sgRNA sequences:
sgRNA #1
CGTTGTAAGCAAAGAGTGCG
sgRNA #2
CTACCACAAATGTCCCACGC
5' arm, 1kb
Puro R
3' arm, 1kb
TK
sgRNA1
sgRNA2
Exon 2
TFAP2C
TFAP2C
5' arm, 1kb
Neo R
3' arm, 1kb
TK
d
WT 1 2 3
TFAP2C
GAPDH
e
HENSM-ACT
HENSM
P+1
P+2
P+1
P+2
WT
TFAP2C-/-
100μm
100μm

Figure S10

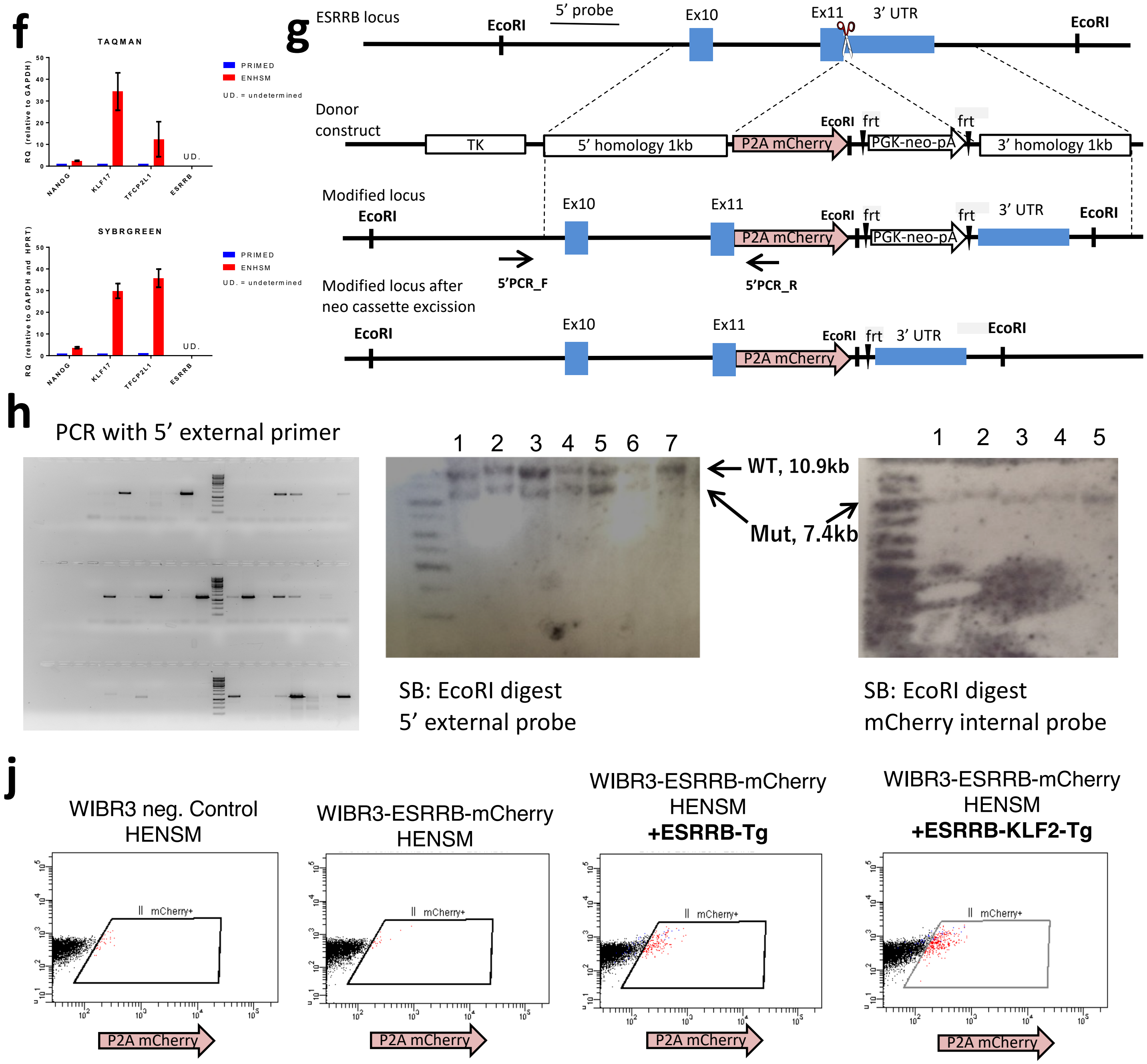


Figure S10..continued

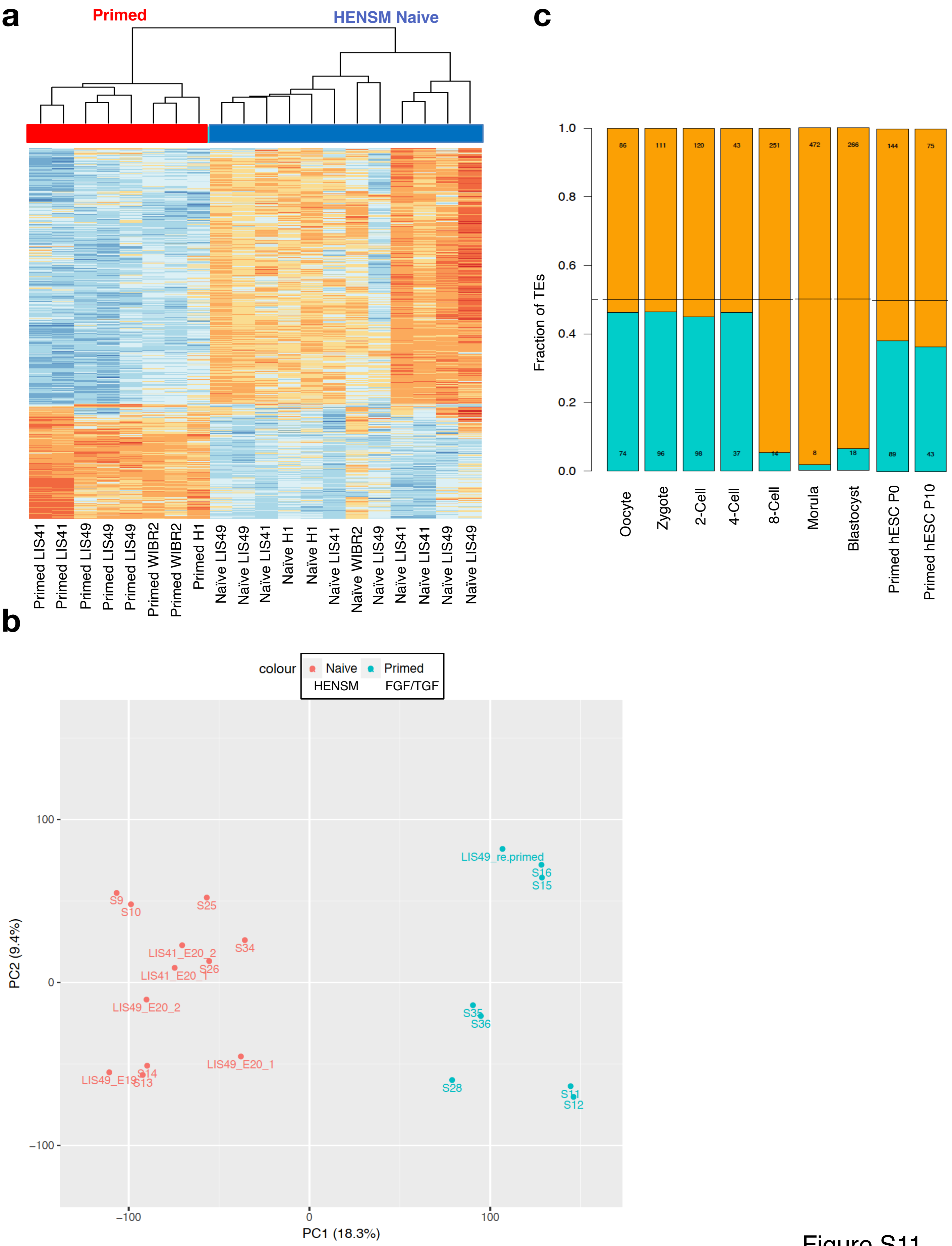
a
Primed
HENSM Naive
Primed LIS41
Primed LIS41
Primed LIS49
Primed LIS49
Primed LIS49
Primed WIBR2
Primed WIBR2
Primed H1
Naïve LIS49
Naïve LIS49
Naïve LIS41
Naïve H1
Naïve H1
Naïve LIS41
Naïve WIBR2
Naïve LIS49
Naïve LIS41
Naïve LIS41
Naïve LIS49
Naïve LIS49
b
colour
Naive HENSM
Primed FGF/TGF
LIS49_re.primed
S16
S15
S9
S10
S25
LIS41_E20_2
S34
S26
LIS41_E20_1
LIS49_E20_2
S35
S36
LIS49_E20_1
S14
LIS49_E19
S13
S28
S11
S12
100
0
−100
PC2 (9.4%)
−100
0
100
PC1 (18.3%)
c
Fraction of TEs
1.0
0.8
0.6
0.4
0.2
0.0
86
111
120
43
251
472
266
144
75
74
96
98
37
14
8
18
89
43
Oocyte
Zygote
2-Cell
4-Cell
8-Cell
Morula
Blastocyst
Primed hESC P0
Primed hESC P10

Figure S11

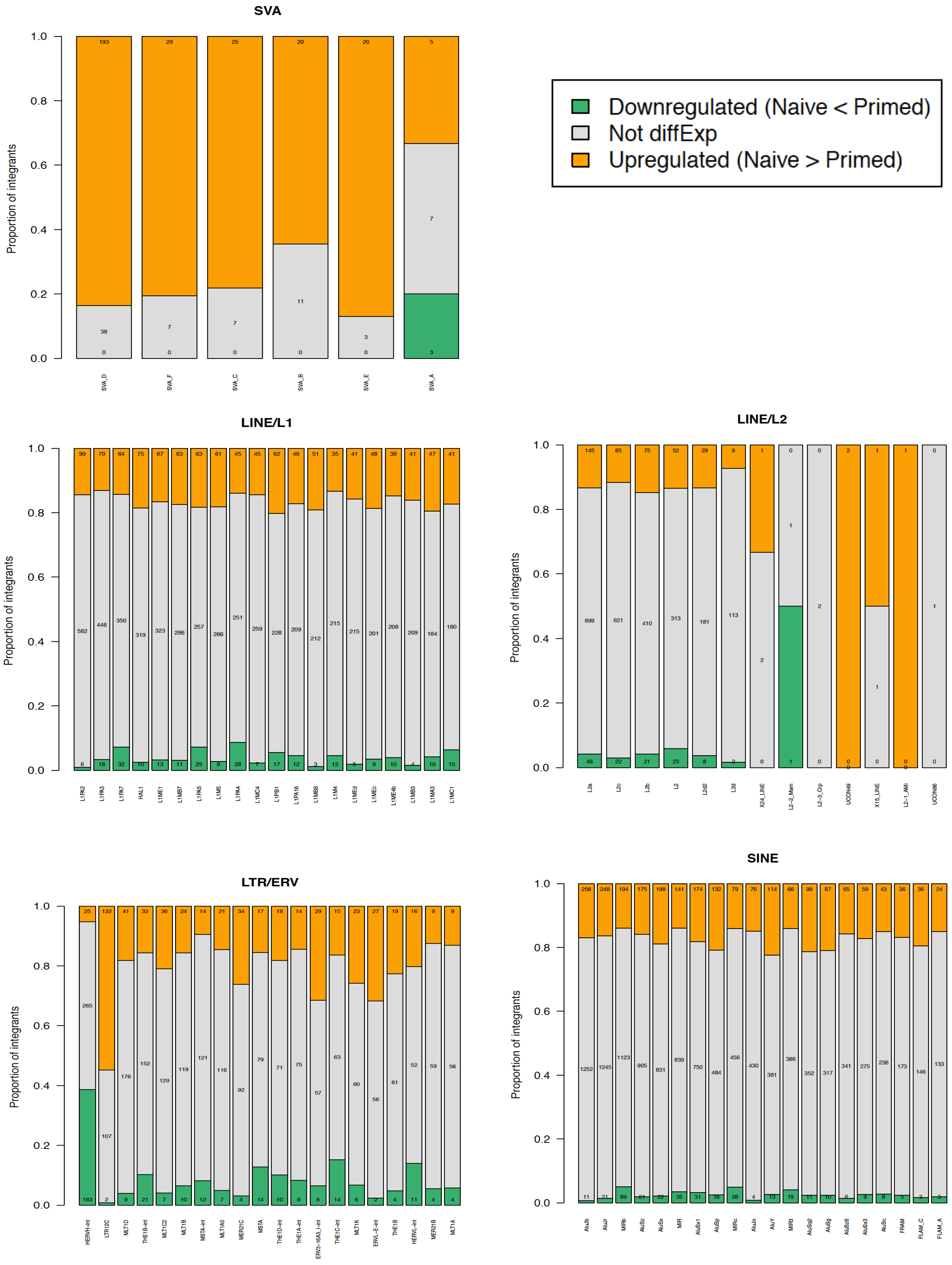
SVA
LINE/L1
LINE/L2
LTR/ERV
SINE
Proportion of integrants
Downregulated (Naive < Primed)
Not diffExp
Upregulated (Naive > Primed)

Figure S12

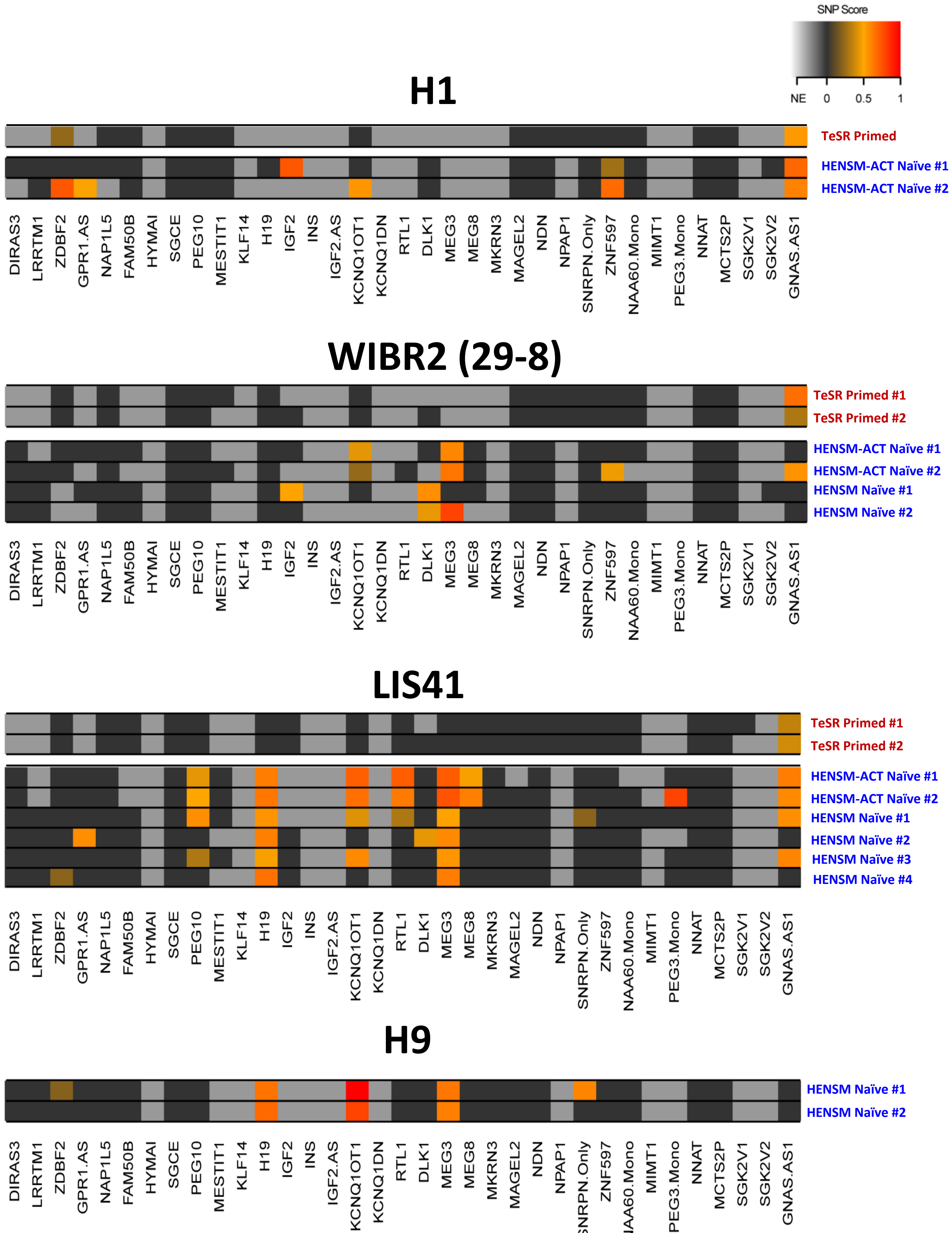
SNP Score
NE 0 0.5 1
H1
TeSR Primed
HENSM-ACT Naïve #1
HENSM-ACT Naïve #2
WIBR2 (29-8)
TeSR Primed #1
TeSR Primed #2
HENSM-ACT Naïve #1
HENSM-ACT Naïve #2
HENSM Naïve #1
HENSM Naïve #2
LIS41
TeSR Primed #1
TeSR Primed #2
HENSM-ACT Naïve #1
HENSM-ACT Naïve #2
HENSM Naïve #1
HENSM Naïve #2
HENSM Naïve #3
HENSM Naïve #4
H9
HENSM Naïve #1
HENSM Naïve #2
DIRAS3 LRRTM1 ZDBF2 GPR1.AS NAP1L5 FAM50B HYMAI SGCE PEG10 MESTIT1 KLF14 H19 IGF2 INS IGF2.AS KCNQ1OT1 KCNQ1DN RTL1 DLK1 MEG3 MEG8 MKRN3 MAGEL2 NDN NPAP1 SNRPN.Only ZNF597 NAA60.Mono MIMT1 PEG3.Mono NNAT MCTS2P SGK2V1 SGK2V2 GNAS.AS1

Figure S13

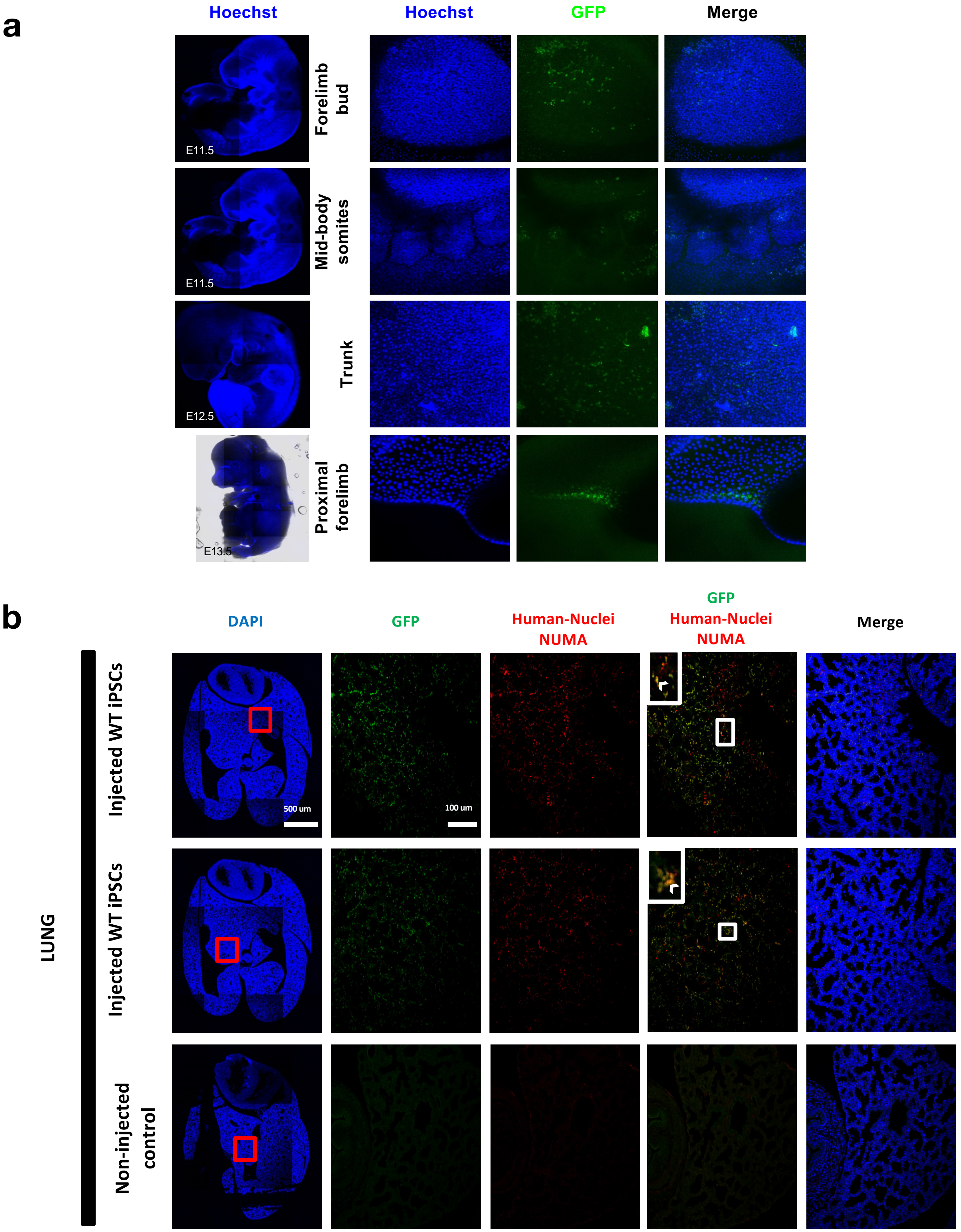
a
Hoechst
Hoechst
GFP
Merge
E11.5
Forelimb bud
E11.5
Mid-body somites
E12.5
Trunk
E13.5
Proximal forelimb
b
DAPI
GFP
Human-Nuclei NUMA
GFP Human-Nuclei NUMA
Merge
LUNG
Injected WT iPSCs
500 um
100 um
Injected WT iPSCs
Non-injected control

Figure S14

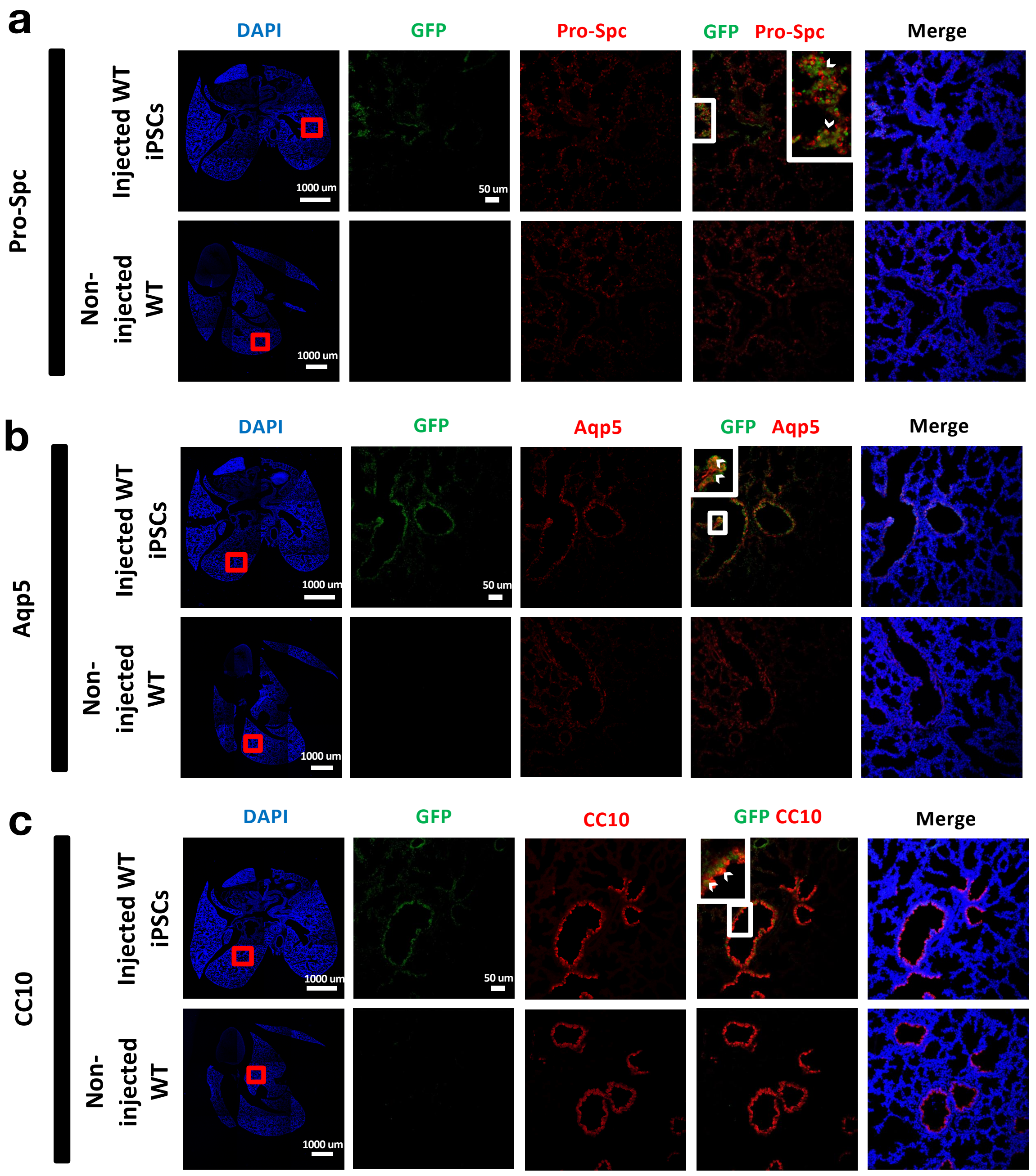
a
DAPI
GFP
Pro-Spc
GFP Pro-Spc
Merge
Pro-Spc
Injected WT iPSCs
Non-injected WT
1000 um
50 um
b
DAPI
GFP
Aqp5
GFP Aqp5
Merge
Aqp5
Injected WT iPSCs
Non-injected WT
1000 um
50 um
c
DAPI
GFP
CC10
GFP CC10
Merge
CC10
Injected WT iPSCs
Non-injected WT
1000 um
50 um

Figure S15

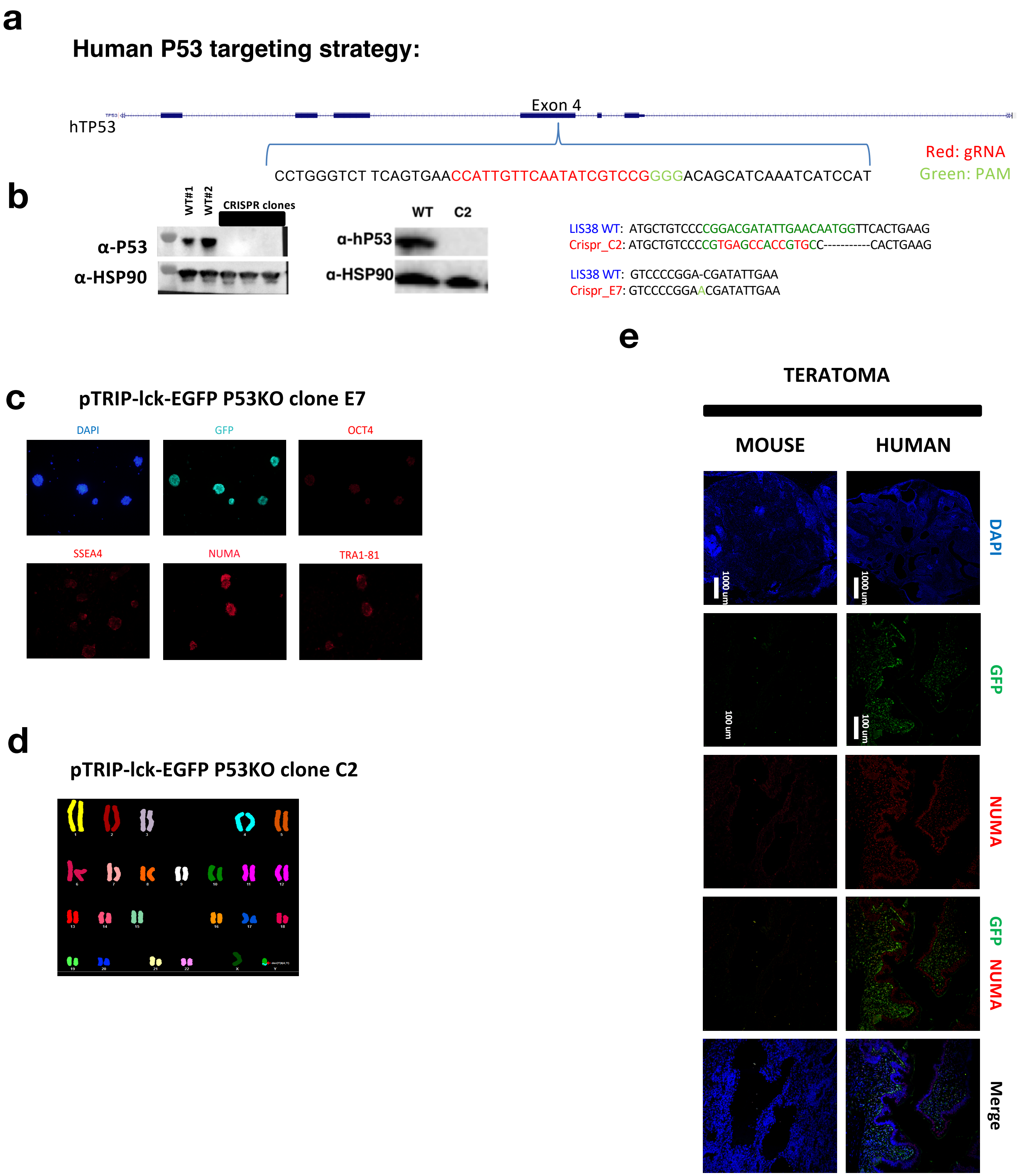
a
Human P53 targeting strategy:
Exon 4
hTP53
CCTGGGTCT TCAGTGAACCATTGTTCAATATCGTCCGGGGACAGCATCAAATCATCCAT
Red: gRNA
Green: PAM
b
WT#1
WT#2
CRISPR clones
α-P53
α-HSP90
WT
C2
α-hP53
α-HSP90
LIS38 WT: ATGCTGTCCCCGGACGATATTGAACAATGGTTCACTGAAG
Crispr_C2: ATGCTGTCCCCGTGAGCCACCGTGCC-----------CACTGAAG
LIS38 WT: GTCCCCGGA-CGATATTGAA
Crispr_E7: GTCCCCGGAACGATATTGAA
c
pTRIP-lck-EGFP P53KO clone E7
DAPI
GFP
OCT4
SSEA4
NUMA
TRA1-81
d
pTRIP-lck-EGFP P53KO clone C2
e
TERATOMA
MOUSE
HUMAN
DAPI
GFP
NUMA
GFP NUMA
Merge
1000 um
100 um

Figure S16

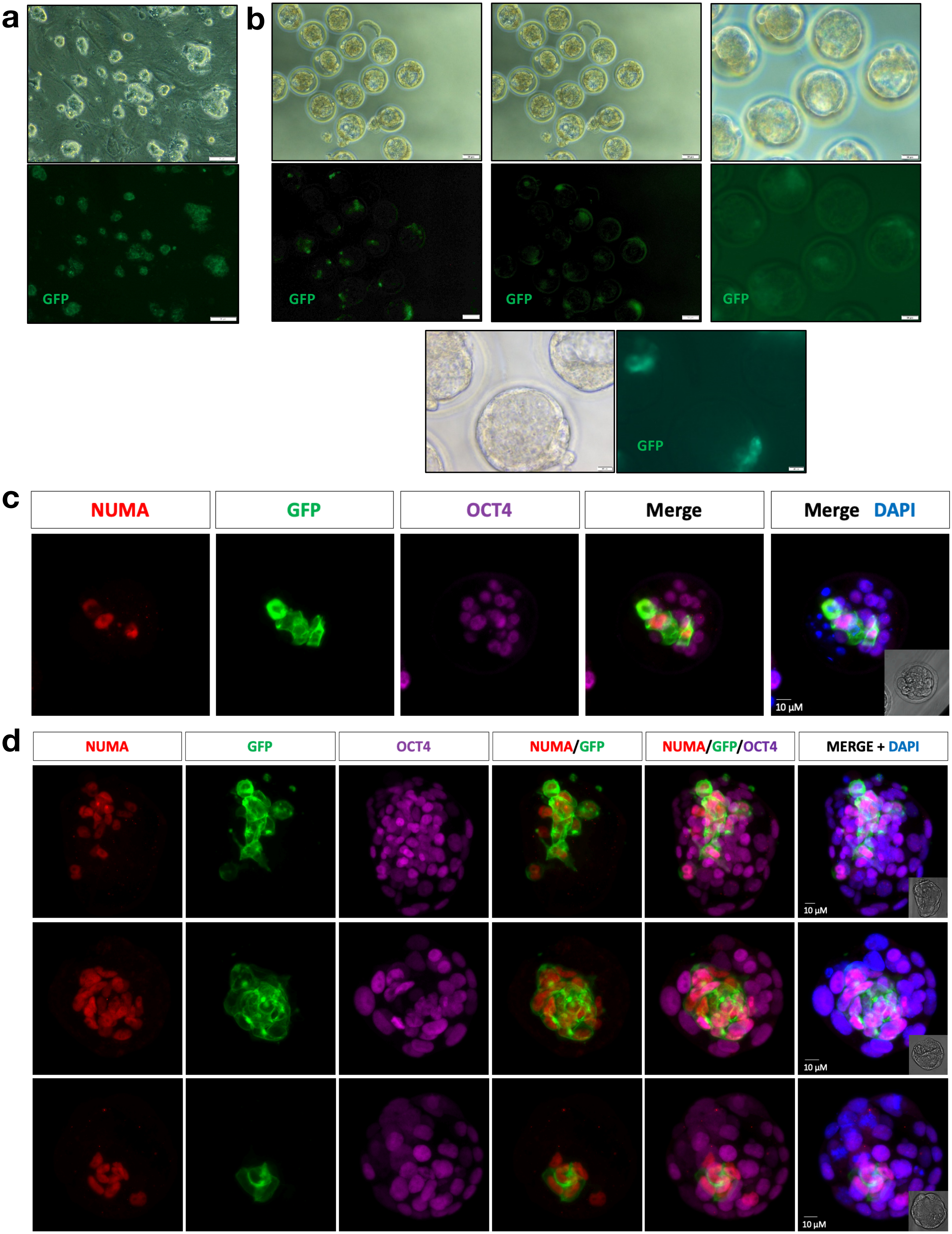

a
b
GFP
c
NUMA
GFP
OCT4
Merge
Merge DAPI
10 µM
d
NUMA
GFP
OCT4
NUMA/GFP
NUMA/GFP/OCT4
MERGE + DAPI
10 µM


Figure S17

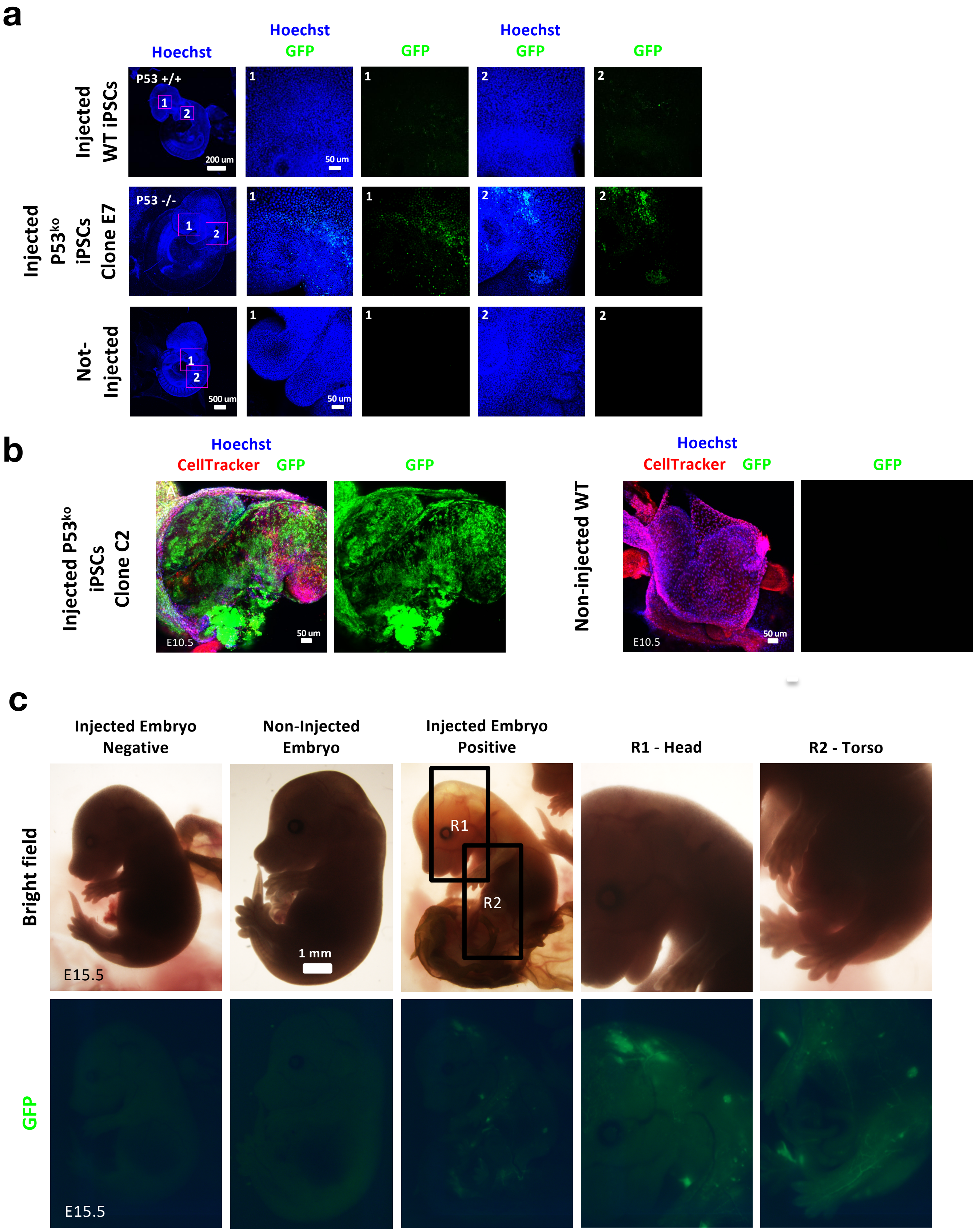
a
Hoechst
Hoechst
GFP
GFP
Hoechst
GFP
GFP
Injected WT iPSCs
P53 +/+
200 um
50 um
Injected P53ko iPSCs Clone E7
P53 -/-
Not-Injected
500 um
50 um
b
Hoechst
CellTracker GFP
GFP
Injected P53ko iPSCs Clone C2
E10.5
50 um
Non-injected WT
c
Injected Embryo Negative
Non-Injected Embryo
Injected Embryo Positive
R1 - Head
R2 - Torso
Bright field
GFP
E15.5
1 mm
R1
R2

Figure S18

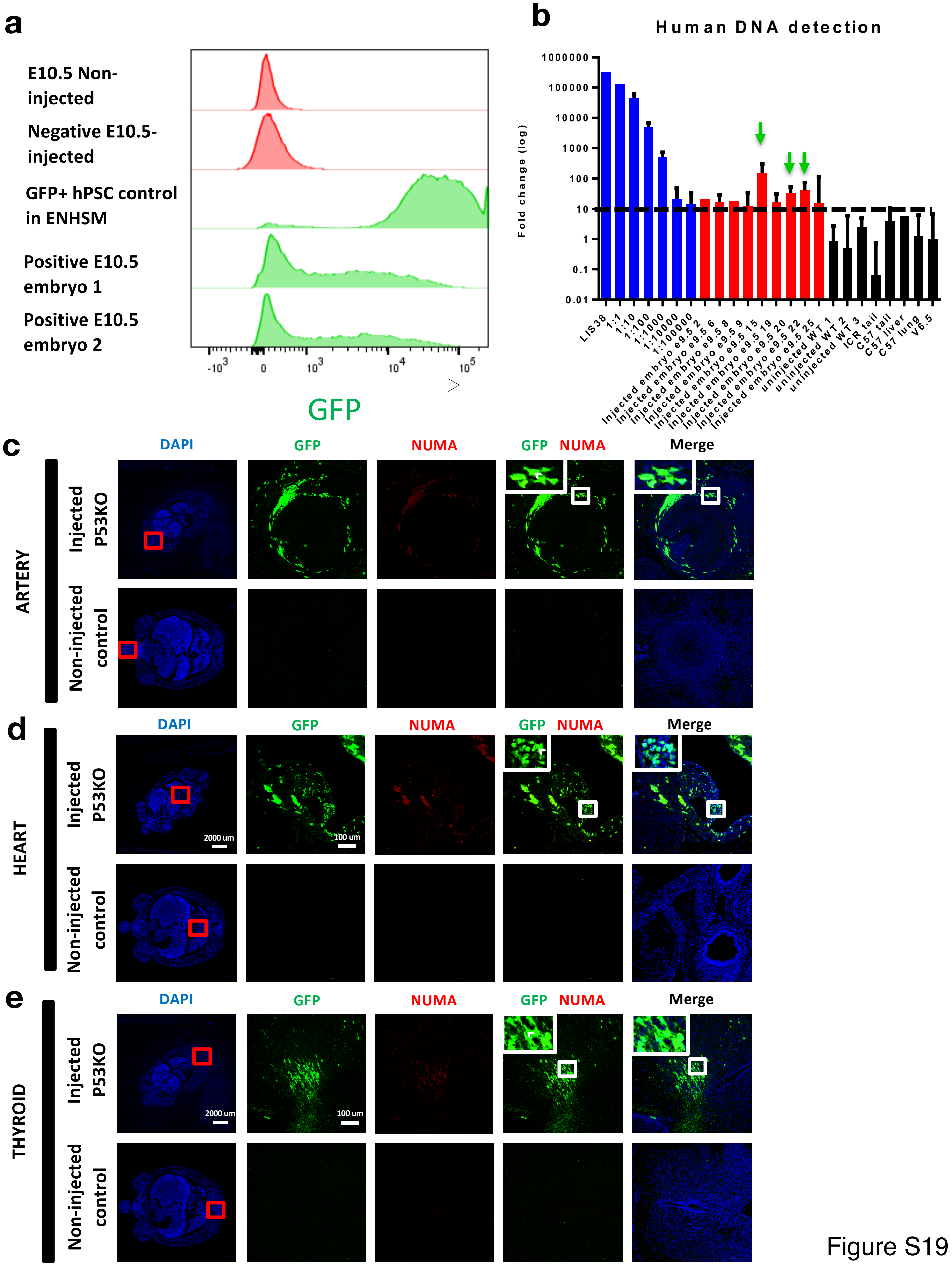
a
E10.5 Non-injected
Negative E10.5-injected
GFP+ hPSC control in ENHSM
Positive E10.5 embryo 1
Positive E10.5 embryo 2
GFP
b
Human DNA detection
Fold change (log)
LIS38
1:1
1:10
1:100
1:1000
1:10000
1:100000
Injected embryo e9.5 2
Injected embryo e9.5 6
Injected embryo e9.5 8
Injected embryo e9.5 9
Injected embryo e9.5 15
Injected embryo e9.5 19
Injected embryo e9.5 20
Injected embryo e9.5 22
Injected embryo e9.5 25
uninjected WT 1
uninjected WT 2
uninjected WT 3
ICR tail
C57 tail
C57 liver
C57 lung
V6.5
c
ARTERY
DAPI
GFP
NUMA
Merge
Injected P53KO
Non-injected control
d
HEART
2000 um
100 um
e
THYROID


Figure S19

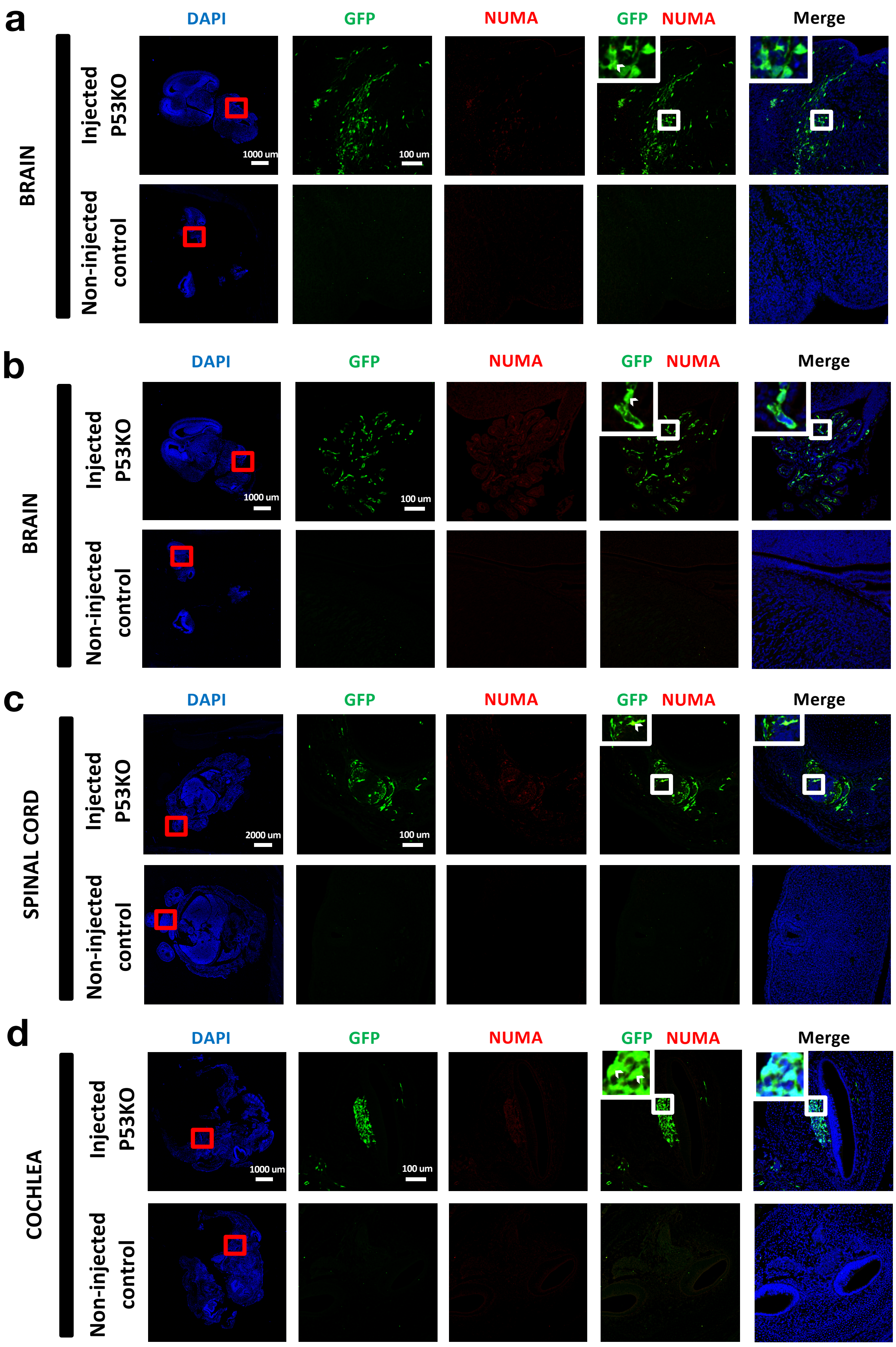
a
BRAIN
Injected P53KO
Non-injected control
DAPI
GFP
NUMA
GFP NUMA
Merge
1000 um
100 um
b
BRAIN
Injected P53KO
Non-injected control
DAPI
GFP
NUMA
GFP NUMA
Merge
1000 um
100 um
c
SPINAL CORD
Injected P53KO
Non-injected control
DAPI
GFP
NUMA
GFP NUMA
Merge
2000 um
100 um
d
COCHLEA
Injected P53KO
Non-injected control
DAPI
GFP
NUMA
GFP NUMA
Merge
1000 um
100 um

Figure S20

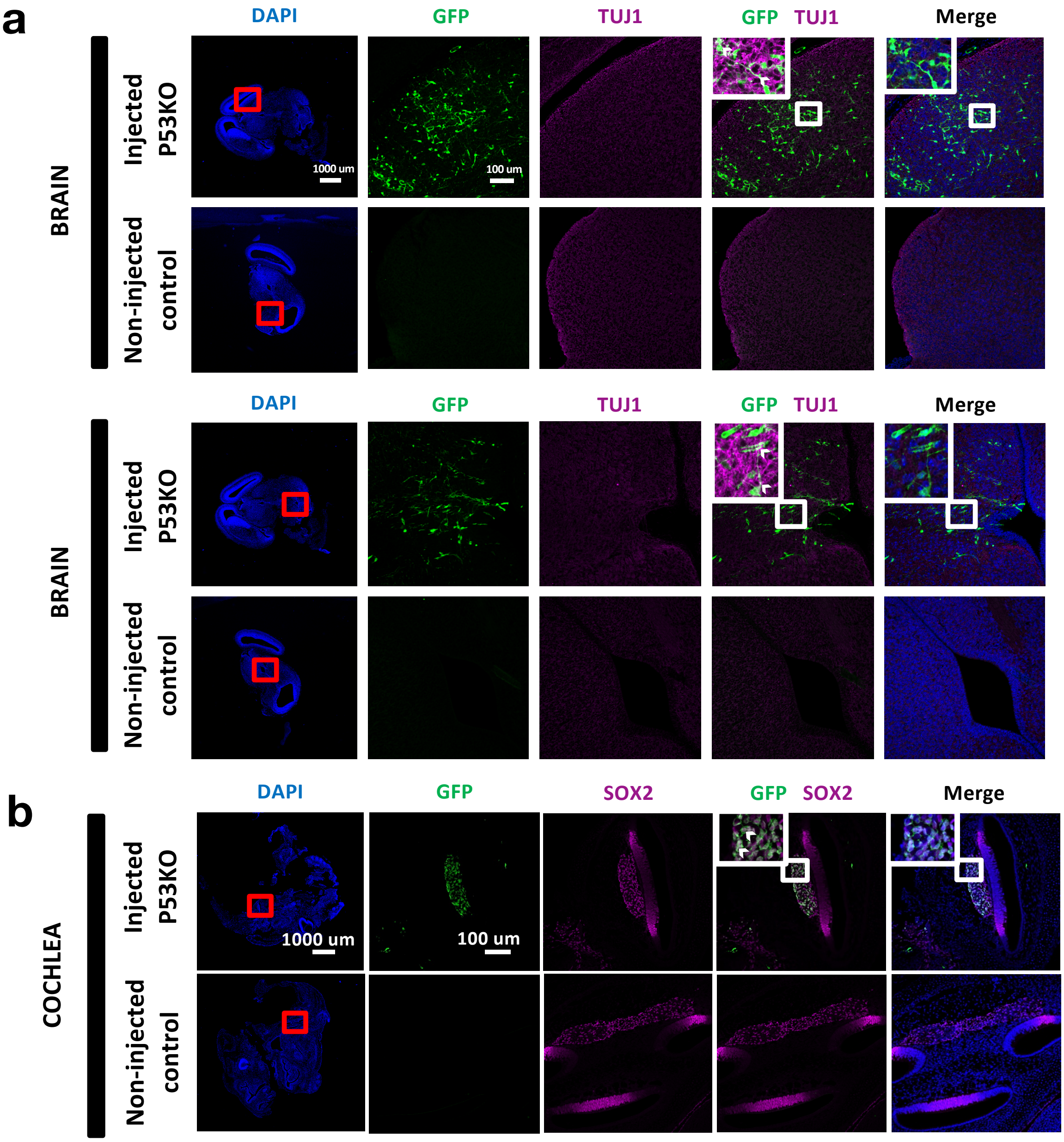
a
DAPI
GFP
TUJ1
GFP TUJ1
Merge
BRAIN
Injected P53KO
Non-injected control
1000 um
100 um
b
SOX2
GFP SOX2
COCHLEA


Figure S21

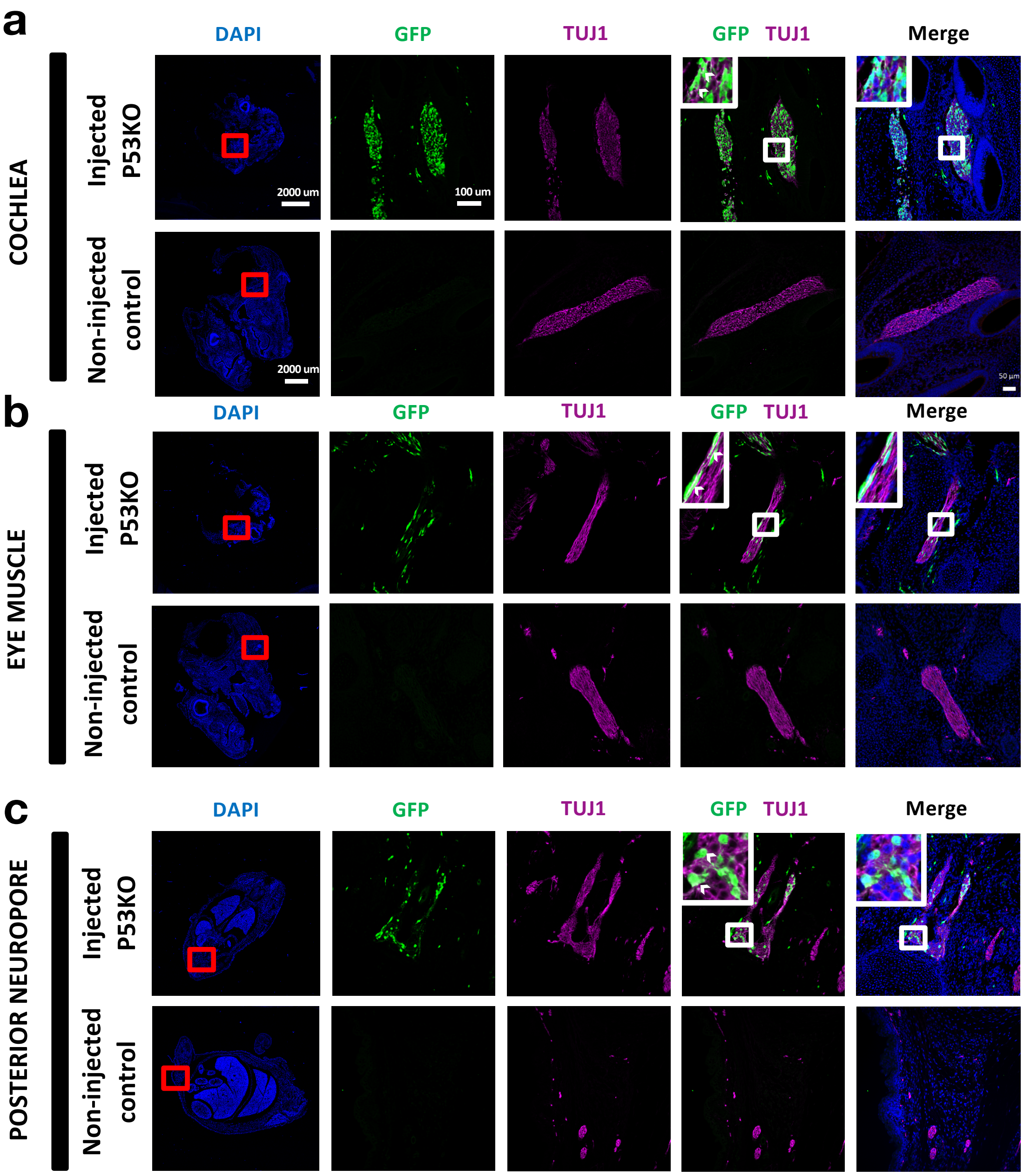
a
DAPI
GFP
TUJ1
GFP TUJ1
Merge
COCHLEA
Injected P53KO
Non-injected control
2000 um
100 um
50 µm
b
EYE MUSCLE
c
POSTERIOR NEUROPORE

Figure S22

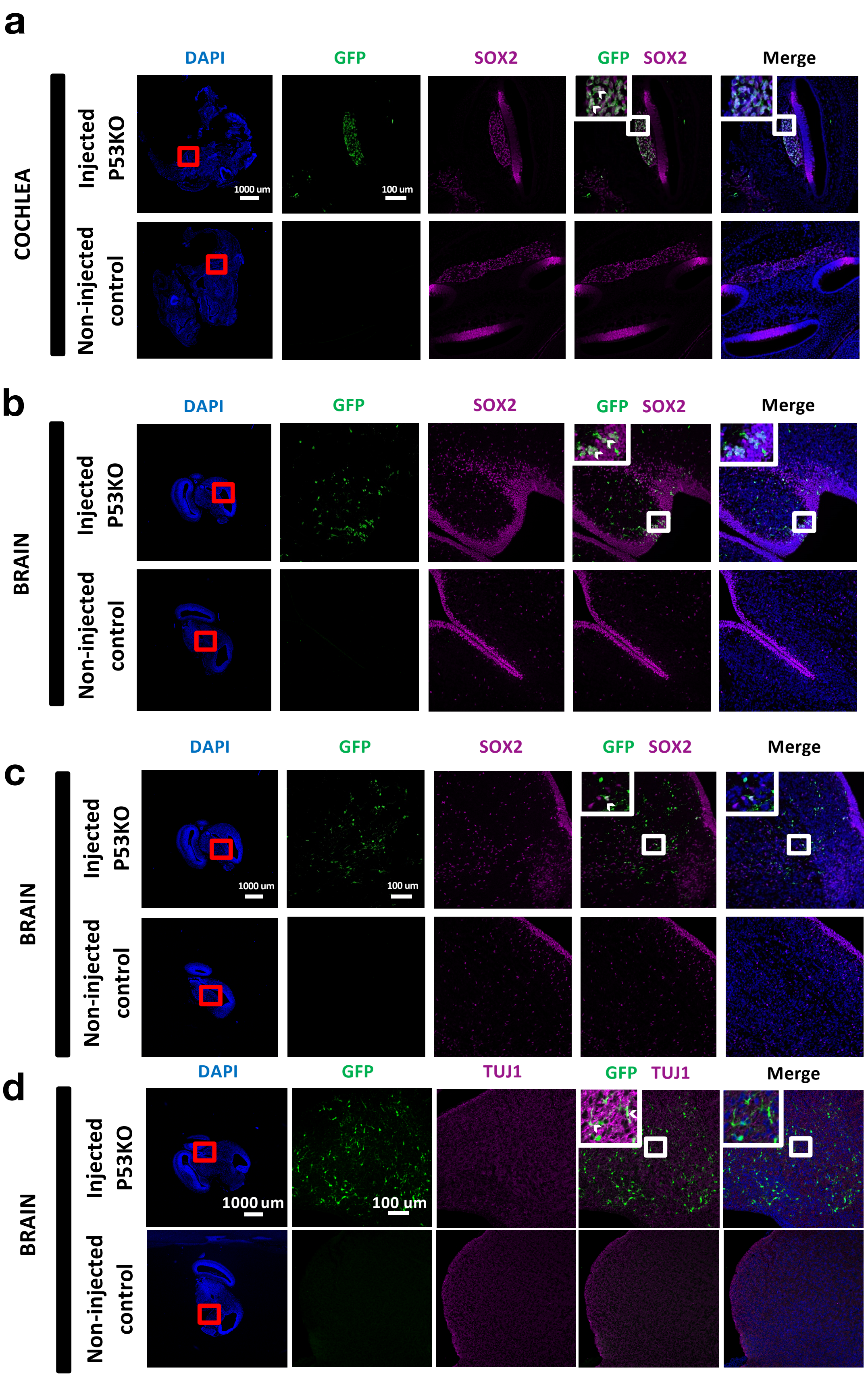
a
DAPI
GFP
SOX2
GFP SOX2
Merge
COCHLEA
Injected P53KO
Non-injected control
1000 um
100 um
b
DAPI
GFP
SOX2
GFP SOX2
Merge
BRAIN
Injected P53KO
Non-injected control
c
DAPI
GFP
SOX2
GFP SOX2
Merge
BRAIN
Injected P53KO
Non-injected control
1000 um
100 um
d
DAPI
GFP
TUJ1
GFP TUJ1
Merge
BRAIN
Injected P53KO
Non-injected control
1000 um
100 um

Figure S23

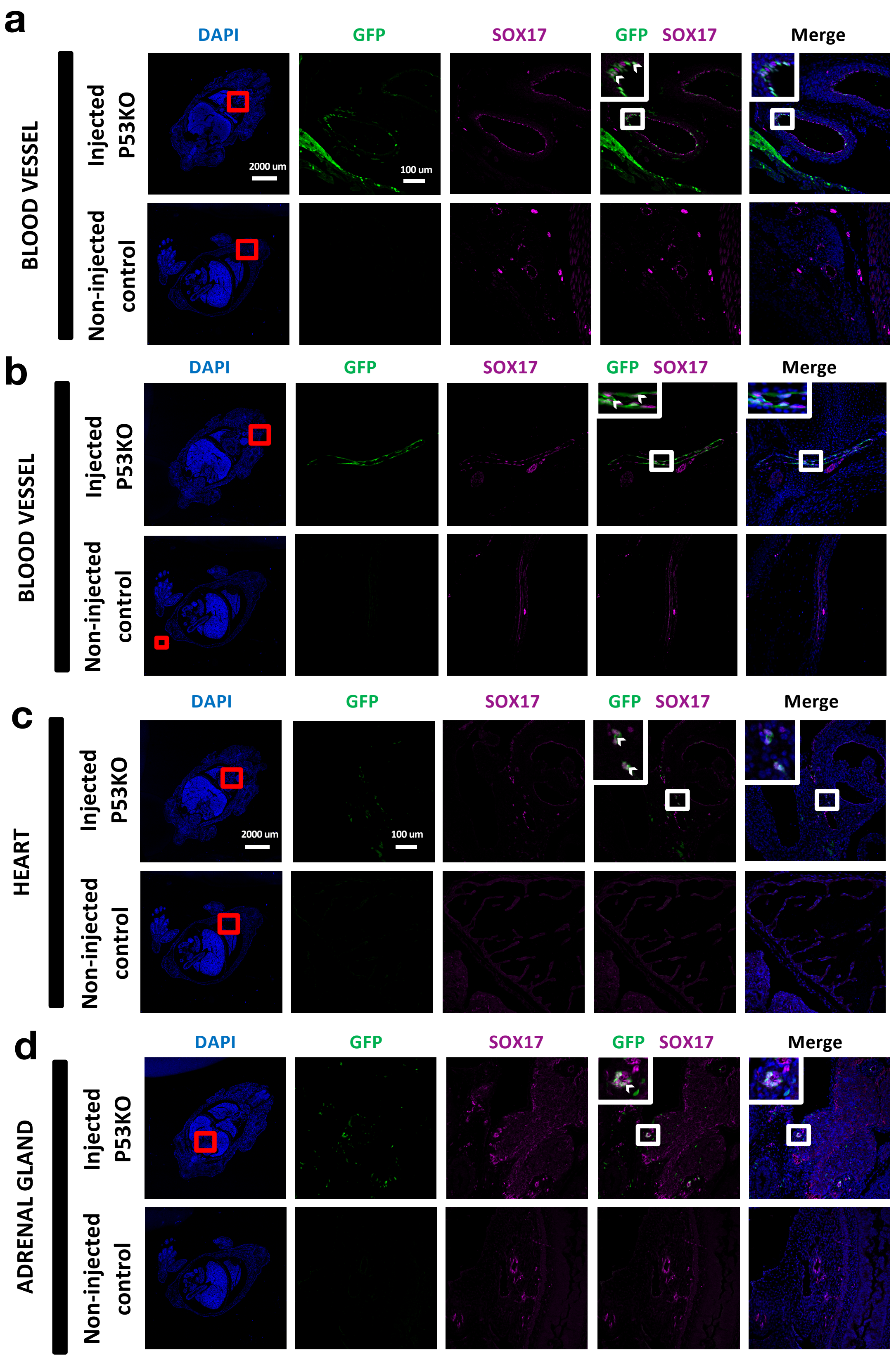
a
DAPI
GFP
SOX17
GFP SOX17
Merge
BLOOD VESSEL
Injected P53KO
Non-injected control
2000 um
100 um
b
DAPI
GFP
SOX17
GFP SOX17
Merge
BLOOD VESSEL
Injected P53KO
Non-injected control
c
DAPI
GFP
SOX17
GFP SOX17
Merge
HEART
Injected P53KO
Non-injected control
2000 um
100 um
d
DAPI
GFP
SOX17
GFP SOX17
Merge
ADRENAL GLAND
Injected P53KO
Non-injected control

Figure S24

**a**

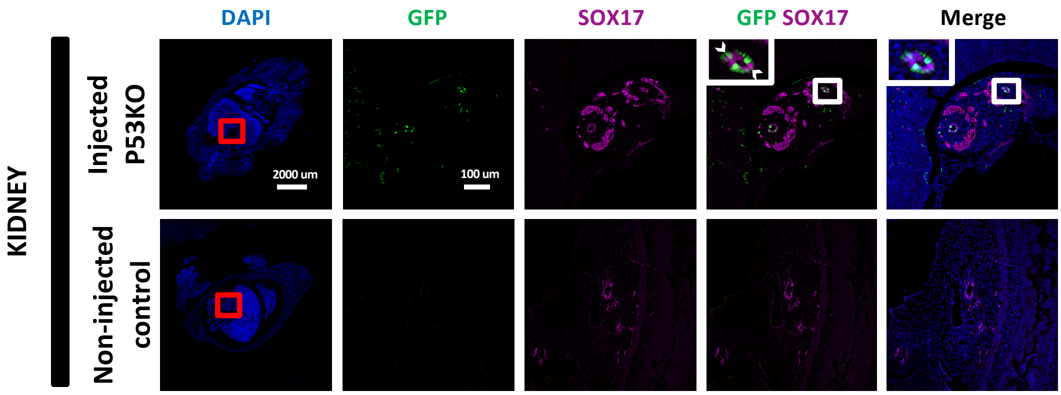


**b**

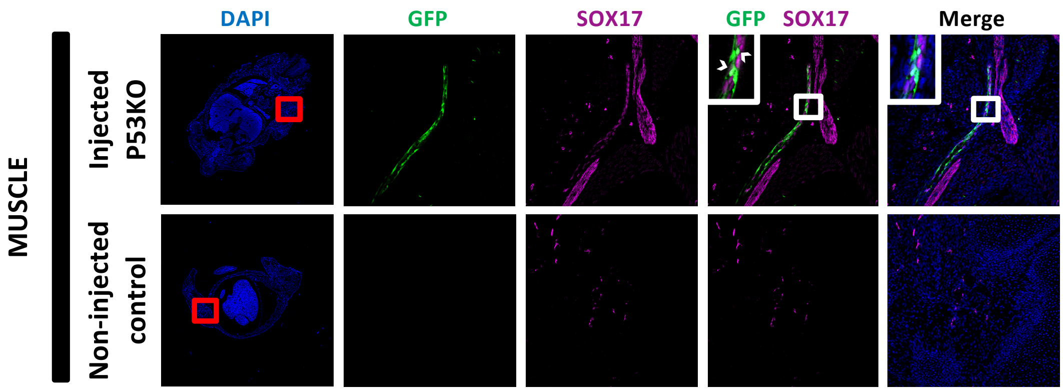


**c**

| Cell lines | Average chimeric embryos per micro-injection batch | Range % of human cells within a chimeric mouse embryo | Embryo age at analysis |
|---|---|---|---|
| Naïve WT hiPSCs | 1 (n=9) | 1-2% | E9.5 – E10.5 |
| **Naïve P53 KO hiPSC** | 4 (n=13) | 1-20% | E9.5 – E10.5 |
| Primed WT hiPSC | 0 (n=5) | 0% | E9.5 – E10.5 |
| Primed P53 KO hiPSC | 0 (n=8) | 0% | E9.5 – E10.5 |

Figure S25

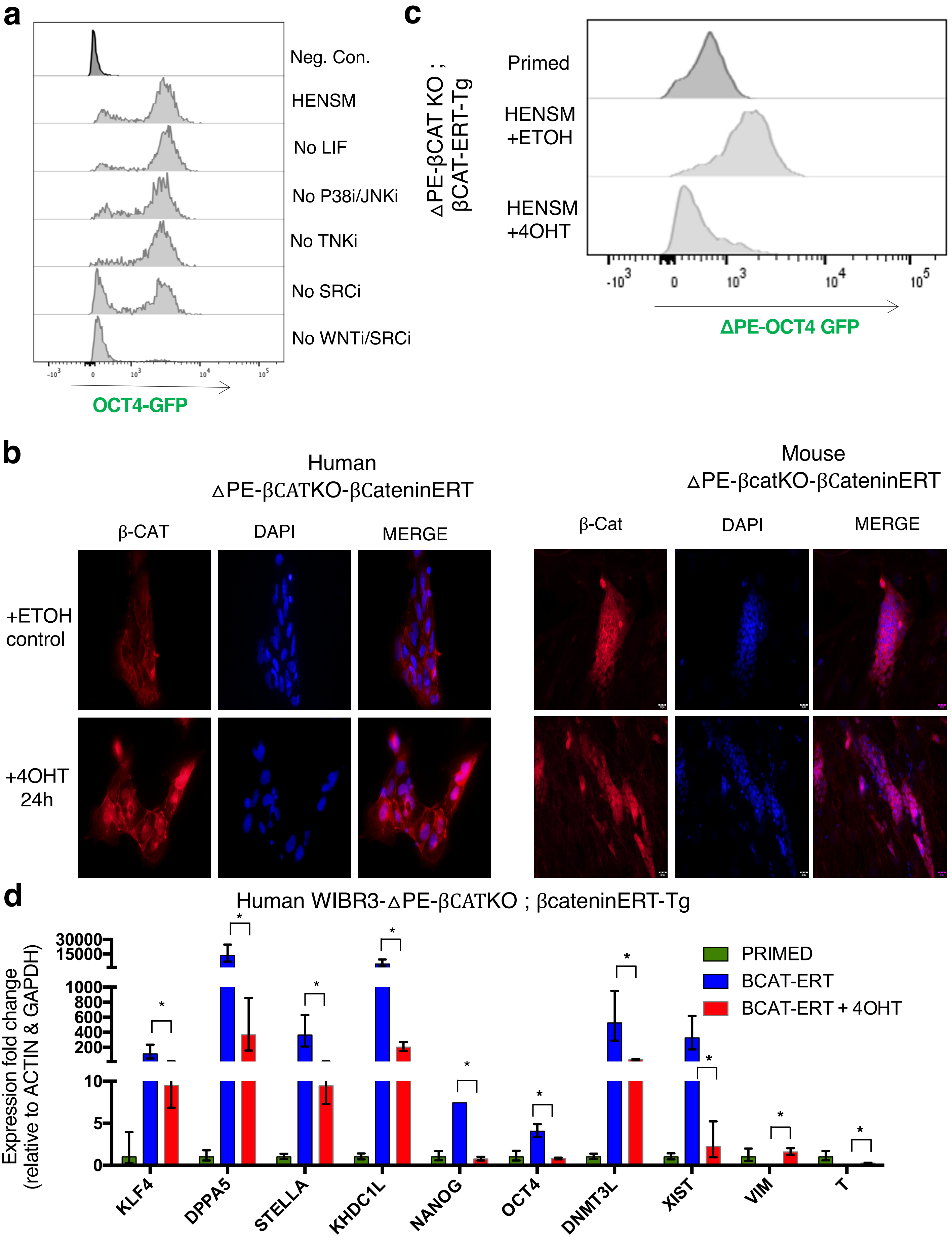
a
Neg. Con.
HENSM
No LIF
No P38i/JNKi
No TNKi
No SRCi
No WNTi/SRCi
OCT4-GFP
c
ΔPE-βCAT KO ; βCAT-ERT-Tg
Primed
HENSM +ETOH
HENSM +4OHT
ΔPE-OCT4 GFP
b
Human ΔPE-βCATKO-βCateninERT
Mouse ΔPE-βcatKO-βCateninERT
β-CAT
DAPI
MERGE
β-Cat
+ETOH control
+4OHT 24h
d
Human WIBR3-ΔPE-βCATKO ; βcateninERT-Tg
Expression fold change (relative to ACTIN & GAPDH)
PRIMED
BCAT-ERT
BCAT-ERT + 4OHT
KLF4
DPPA5
STELLA
KHDC1L
NANOG
OCT4
DNMT3L
XIST
VIM
T

Figure S26

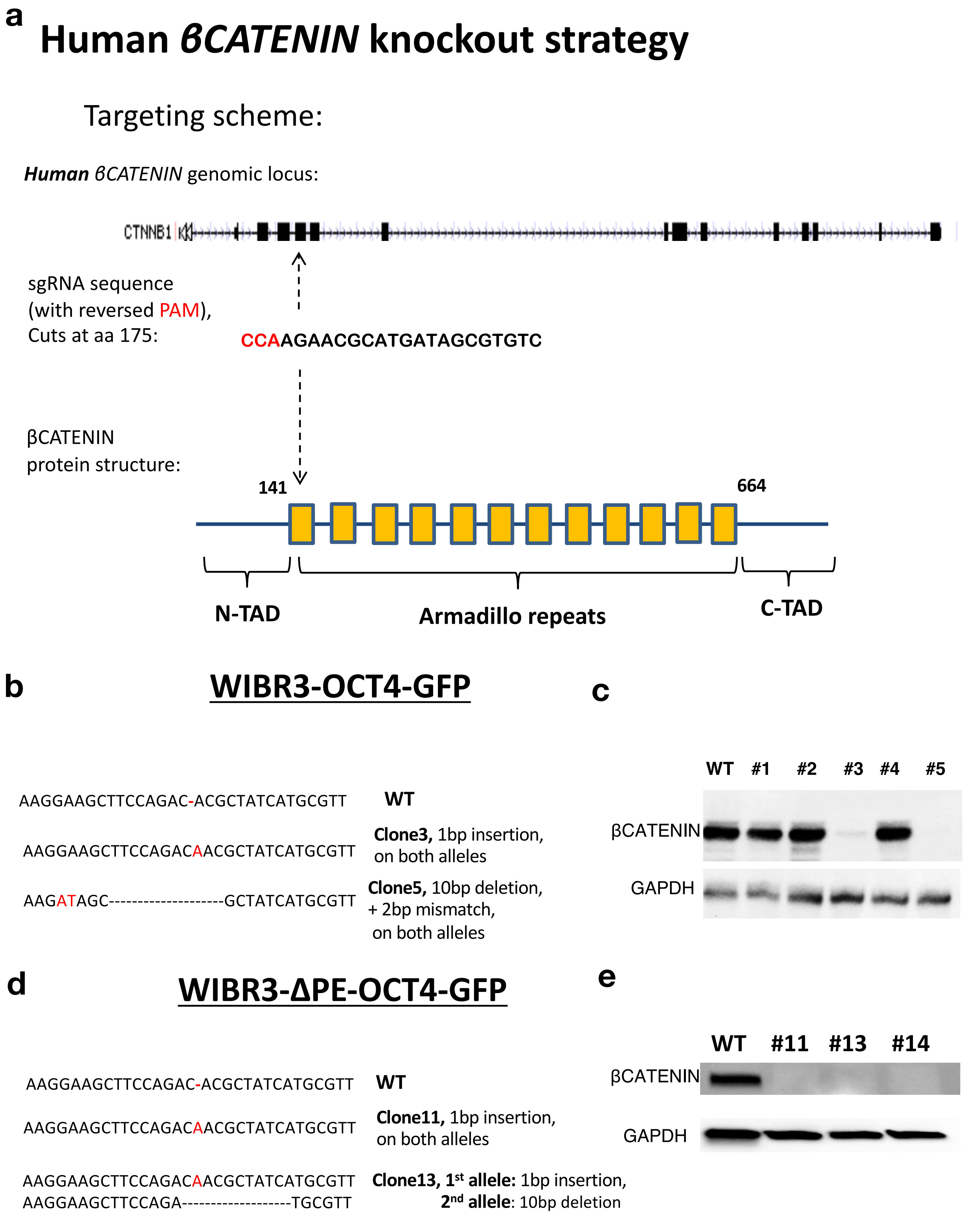
a
Human βCATENIN knockout strategy
Targeting scheme:
Human βCATENIN genomic locus:
CTNNB1
sgRNA sequence
(with reversed PAM),
Cuts at aa 175:
CCAAGAACGCATGATAGCGTGTC
βCATENIN
protein structure:
141
664
N-TAD
Armadillo repeats
C-TAD
b
WIBR3-OCT4-GFP
AAGGAAGCTTCCAGAC-ACGCTATCATGCGTT
WT
AAGGAAGCTTCCAGACAACGCTATCATGCGTT
Clone3, 1bp insertion,
on both alleles
AAGATAGC--------------------GCTATCATGCGTT
Clone5, 10bp deletion,
+ 2bp mismatch,
on both alleles
c
WT #1 #2 #3 #4 #5
βCATENIN
GAPDH
d
WIBR3-ΔPE-OCT4-GFP
AAGGAAGCTTCCAGAC-ACGCTATCATGCGTT
WT
AAGGAAGCTTCCAGACAACGCTATCATGCGTT
Clone11, 1bp insertion,
on both alleles
AAGGAAGCTTCCAGACAACGCTATCATGCGTT
AAGGAAGCTTCCAGA-------------------TGCGTT
Clone13, 1st allele: 1bp insertion,
2nd allele: 10bp deletion
e
WT #11 #13 #14
βCATENIN
GAPDH

Figure S27

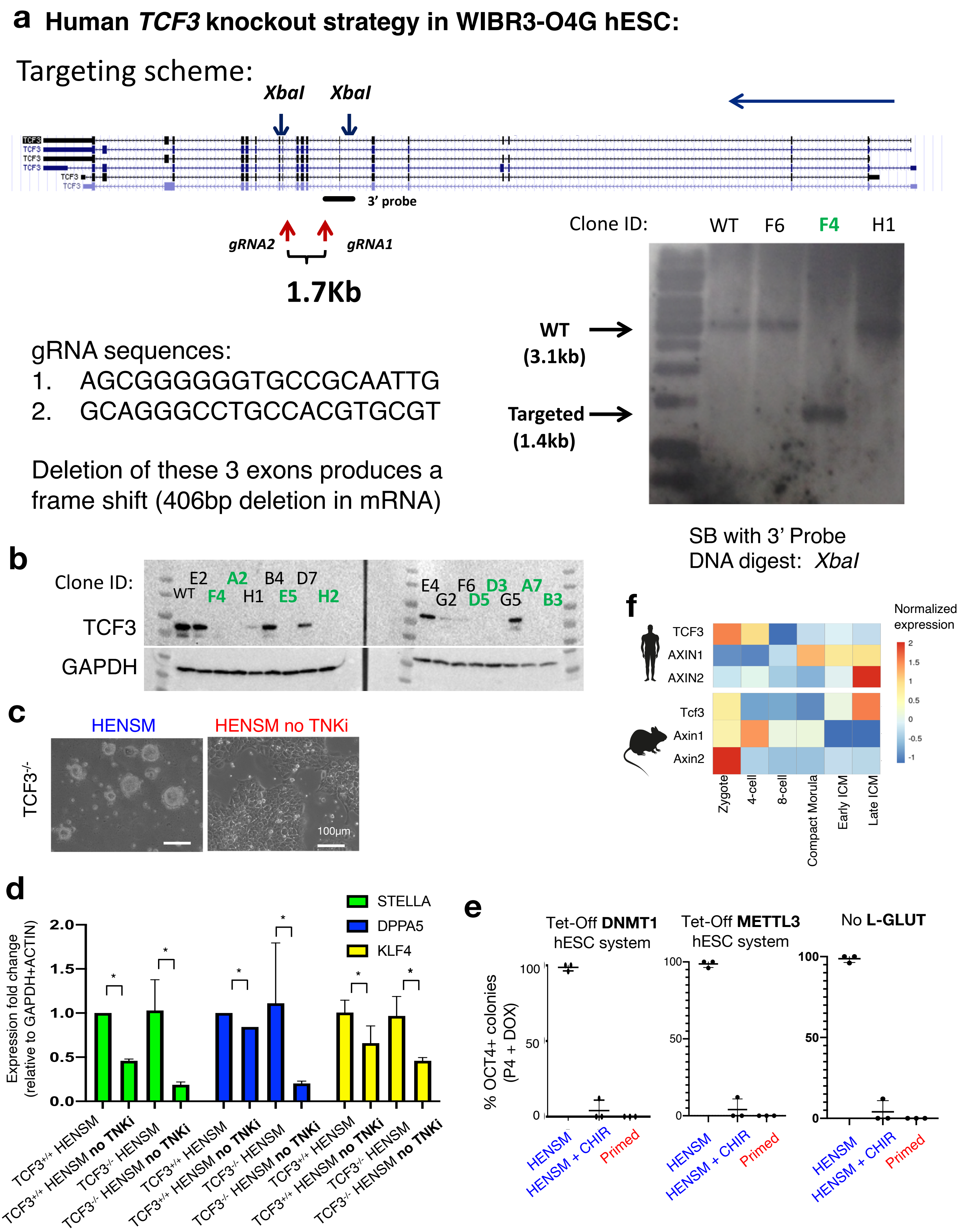
a
Human TCF3 knockout strategy in WIBR3-O4G hESC:
Targeting scheme:
XbaI
XbaI
3' probe
gRNA2
gRNA1
1.7Kb
gRNA sequences:
1. AGCGGGGGTGCCGCAATTG
2. GCAGGGCCTGCCACGTGCGT
Deletion of these 3 exons produces a frame shift (406bp deletion in mRNA)
Clone ID: WT F6 F4 H1
WT (3.1kb)
Targeted (1.4kb)
SB with 3' Probe
DNA digest: XbaI
b
Clone ID:
WT E2 F4 A2 H1 B4 E5 D7 H2
E4 G2 F6 D5 D3 G5 A7 B3
TCF3
GAPDH
c
HENSM
HENSM no TNKi
TCF3-/-
100µm
d
STELLA
DPPA5
KLF4
Expression fold change
(relative to GAPDH+ACTIN)
2.0
1.5
1.0
0.5
0.0
TCF3+/+ HENSM
TCF3+/+ HENSM no TNKi
TCF3-/- HENSM
TCF3-/- HENSM no TNKi
e
Tet-Off DNMT1 hESC system
Tet-Off METTL3 hESC system
No L-GLUT
% OCT4+ colonies
(P4 + DOX)
100
50
0
HENSM
HENSM + CHIR
Primed
f
Normalized expression
TCF3
AXIN1
AXIN2
Tcf3
Axin1
Axin2
Zygote
4-cell
8-cell
Compact Morula
Early ICM
Late ICM
2
1.5
1
0.5
0
-0.5
-1

Figure S28

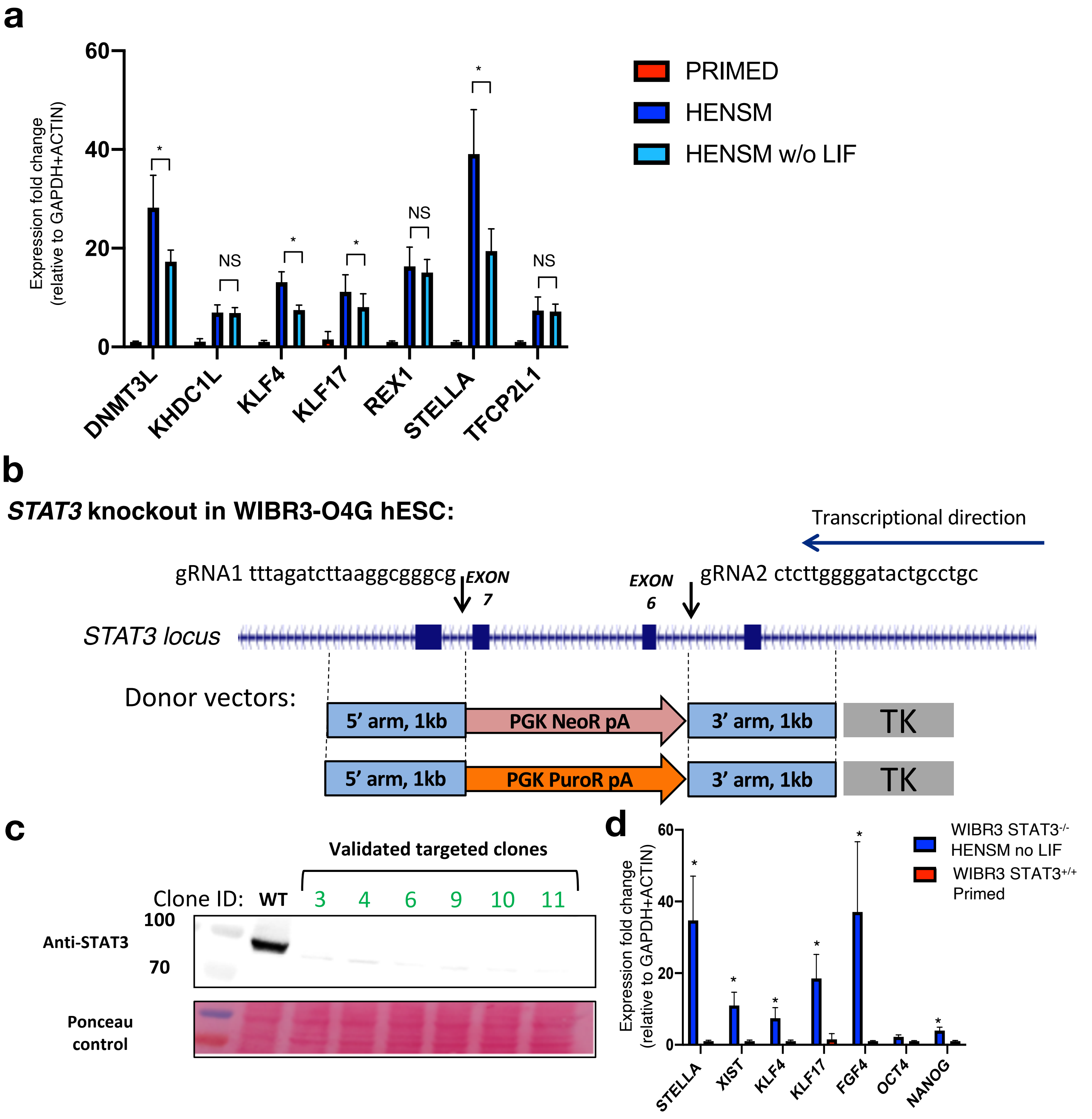
a
60
40
20
0
Expression fold change
(relative to GAPDH+ACTIN)
*
NS
*
*
NS
*
NS
DNMT3L
KHDC1L
KLF4
KLF17
REX1
STELLA
TFCP2L1
PRIMED
HENSM
HENSM w/o LIF
b
STAT3 knockout in WIBR3-O4G hESC:
Transcriptional direction
gRNA1 tttagatcttaaggcgggcg
EXON 7
EXON 6
gRNA2 ctcttggggatactgcctgc
STAT3 locus
Donor vectors:
5' arm, 1kb
PGK NeoR pA
3' arm, 1kb
TK
5' arm, 1kb
PGK PuroR pA
3' arm, 1kb
TK
c
Validated targeted clones
Clone ID:
WT
3
4
6
9
10
11
100
70
Anti-STAT3
Ponceau control
d
60
40
20
0
Expression fold change
(relative to GAPDH+ACTIN)
*
*
*
*
*
*
WIBR3 STAT3-/- HENSM no LIF
WIBR3 STAT3+/+ Primed
STELLA
XIST
KLF4
KLF17
FGF4
OCT4
NANOG

Figure S29

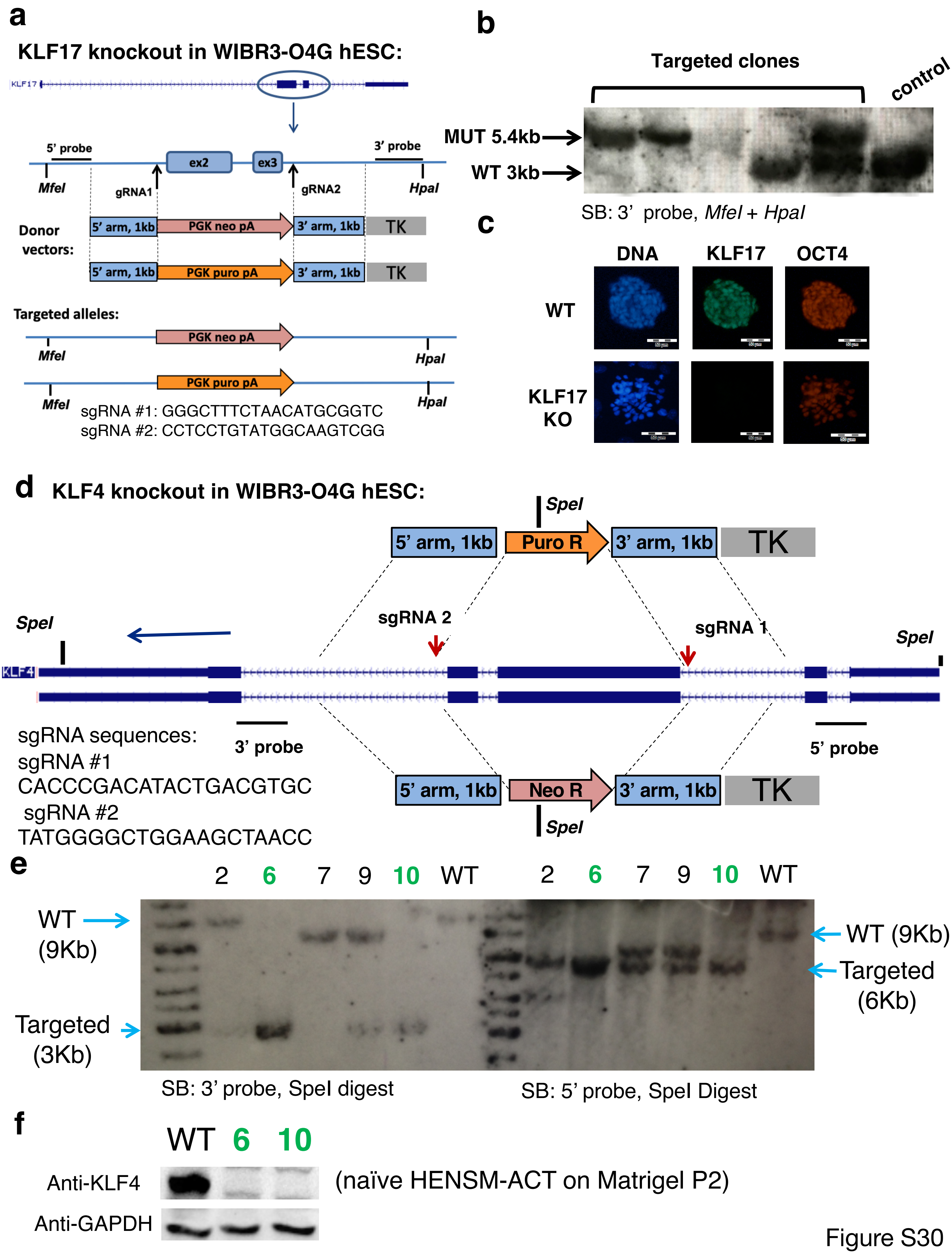
a
KLF17 knockout in WIBR3-O4G hESC:
KLF17
5' probe
3' probe
ex2
ex3
MfeI
gRNA1
gRNA2
HpaI
Donor vectors:
5' arm, 1kb
PGK neo pA
3' arm, 1kb
TK
5' arm, 1kb
PGK puro pA
3' arm, 1kb
TK
Targeted alleles:
PGK neo pA
MfeI
HpaI
PGK puro pA
MfeI
HpaI
sgRNA #1: GGGCTTTCTAACATGCGGTC
sgRNA #2: CCTCCTGTATGGCAAGTCGG
b
Targeted clones
control
MUT 5.4kb
WT 3kb
SB: 3' probe, MfeI + HpaI
c
DNA
KLF17
OCT4
WT
KLF17 KO
d
KLF4 knockout in WIBR3-O4G hESC:
SpeI
5' arm, 1kb
Puro R
3' arm, 1kb
TK
SpeI
sgRNA 2
sgRNA 1
SpeI
KLF4
sgRNA sequences:
sgRNA #1
CACCCGACATACTGACGTGC
sgRNA #2
TATGGGGCTGGAAGCTAACC
3' probe
5' probe
5' arm, 1kb
Neo R
3' arm, 1kb
TK
SpeI
e
2 6 7 9 10 WT 2 6 7 9 10 WT
WT (9Kb)
Targeted (3Kb)
WT (9Kb)
Targeted (6Kb)
SB: 3' probe, SpeI digest
SB: 5' probe, SpeI Digest
f
WT 6 10
Anti-KLF4
Anti-GAPDH
(naïve HENSM-ACT on Matrigel P2)

Figure S30

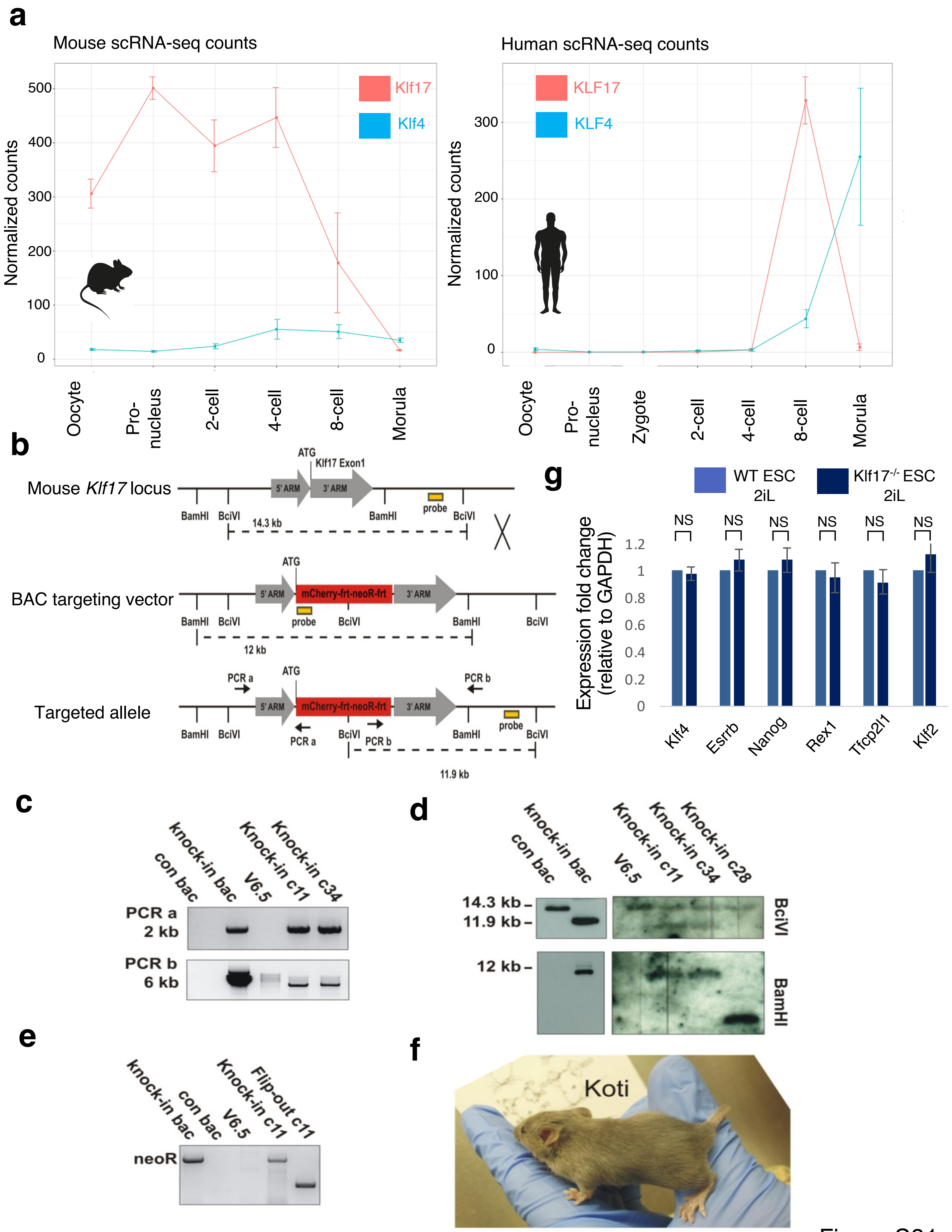
a
Mouse scRNA-seq counts
Klf17
Klf4
Normalized counts
0
100
200
300
400
500
Oocyte
Pro-nucleus
2-cell
4-cell
8-cell
Morula
Human scRNA-seq counts
KLF17
KLF4
Normalized counts
0
100
200
300
Oocyte
Pro-nucleus
Zygote
2-cell
4-cell
8-cell
Morula
b
Mouse Klf17 locus
ATG
Klf17 Exon1
5' ARM
3' ARM
BamHI
BciVI
14.3 kb
BamHI
probe
BciVI
BAC targeting vector
ATG
5' ARM
mCherry-frt-neoR-frt
3' ARM
BamHI
BciVI
probe
BciVI
BamHI
BciVI
12 kb
Targeted allele
PCR a
ATG
PCR b
5' ARM
mCherry-frt-neoR-frt
3' ARM
BamHI
BciVI
PCR a
BciVI
PCR b
BamHI
probe
BciVI
11.9 kb
g
WT ESC 2iL
Klf17-/- ESC 2iL
NS
Expression fold change (relative to GAPDH)
0
0.2
0.4
0.6
0.8
1
1.2
Klf4
Esrrb
Nanog
Rex1
Tfcp2l1
Klf2
c
con bac
knock-in bac
V6.5
Knock-in c11
Knock-in c34
PCR a 2 kb
PCR b 6 kb
d
con bac
knock-in bac
V6.5
Knock-in c11
Knock-in c34
Knock-in c28
14.3 kb
11.9 kb
12 kb
BciVI
BamHI
e
knock-in bac
con bac
V6.5
Knock-in c11
Flip-out c11
neoR
f
Koti

Figure S31

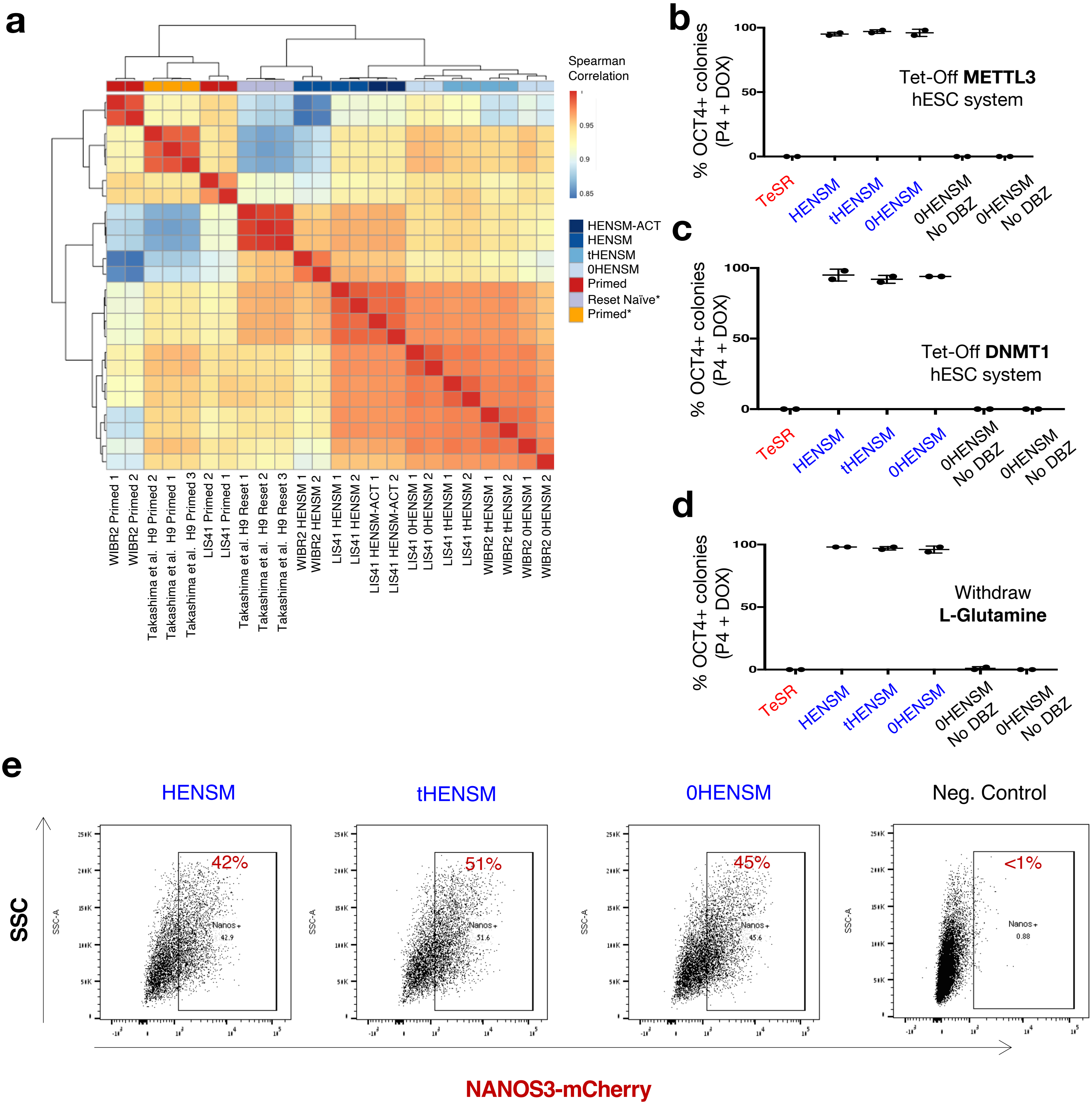
a
Spearman Correlation
HENSM-ACT
HENSM
tHENSM
0HENSM
Primed
Reset Naïve*
Primed*
WIBR2 Primed 1
WIBR2 Primed 2
Takashima et al. H9 Primed 2
Takashima et al. H9 Primed 1
Takashima et al. H9 Primed 3
LIS41 Primed 2
LIS41 Primed 1
Takashima et al. H9 Reset 1
Takashima et al. H9 Reset 2
Takashima et al. H9 Reset 3
WIBR2 HENSM 1
WIBR2 HENSM 2
LIS41 HENSM 1
LIS41 HENSM 2
LIS41 HENSM-ACT 1
LIS41 HENSM-ACT 2
LIS41 0HENSM 1
LIS41 0HENSM 2
LIS41 tHENSM 1
LIS41 tHENSM 2
WIBR2 tHENSM 1
WIBR2 tHENSM 2
WIBR2 0HENSM 1
WIBR2 0HENSM 2
b
% OCT4+ colonies (P4 + DOX)
Tet-Off METTL3 hESC system
TeSR
HENSM
tHENSM
0HENSM
0HENSM No DBZ
0HENSM No DBZ
c
% OCT4+ colonies (P4 + DOX)
Tet-Off DNMT1 hESC system
d
% OCT4+ colonies (P4 + DOX)
Withdraw L-Glutamine
e
HENSM
tHENSM
0HENSM
Neg. Control
SSC
42%
51%
45%
<1%
Nanos+
NANOS3-mCherry

Figure S32

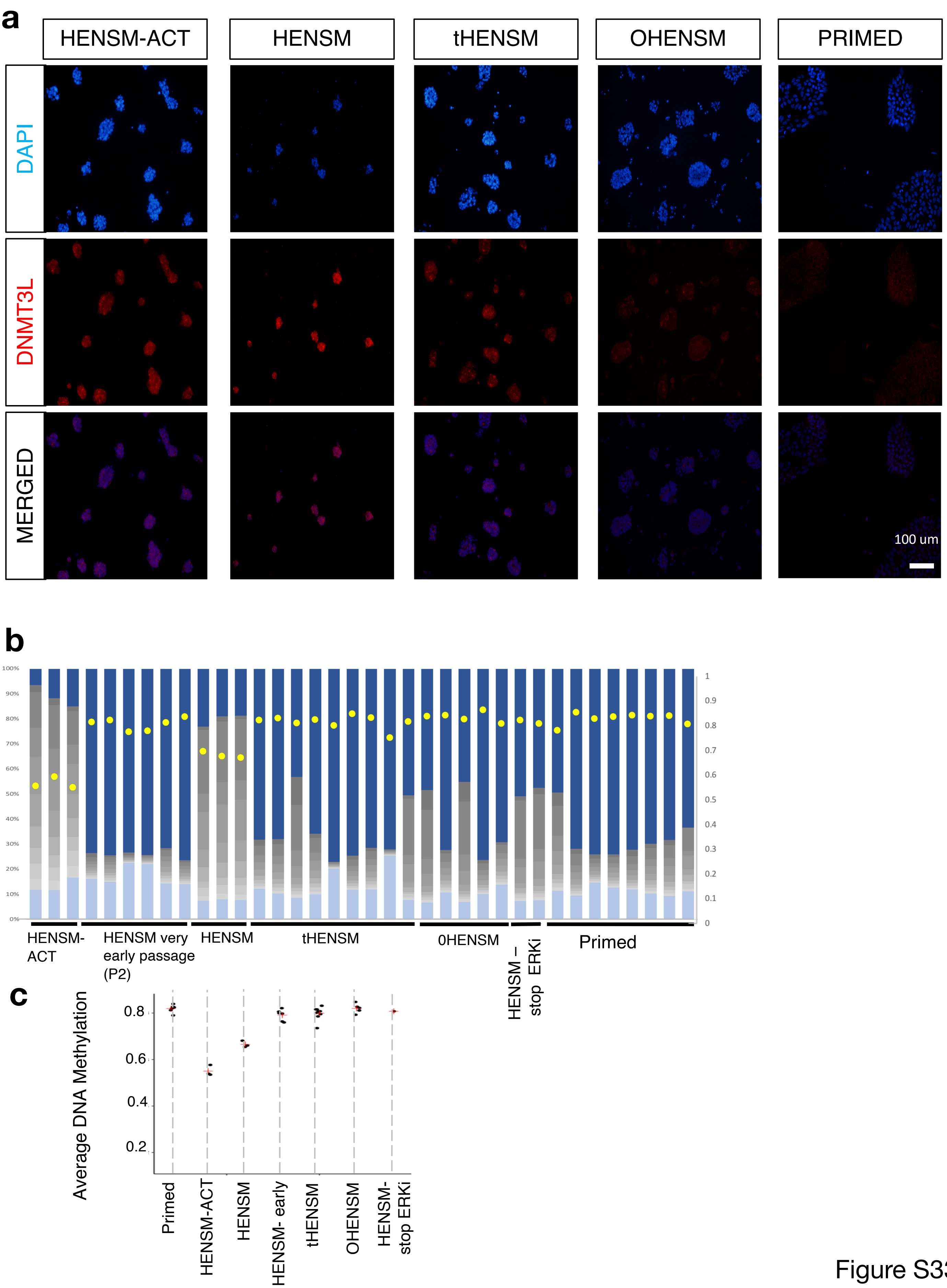
a
HENSM-ACT
HENSM
tHENSM
OHENSM
PRIMED
DAPI
DNMT3L
MERGED
100 um
b
HENSM-ACT
HENSM very early passage (P2)
HENSM
tHENSM
0HENSM
HENSM – stop ERKi
Primed
c
Average DNA Methylation
Primed
HENSM-ACT
HENSM
HENSM- early
tHENSM
OHENSM
HENSM- stop ERKi

Figure S33

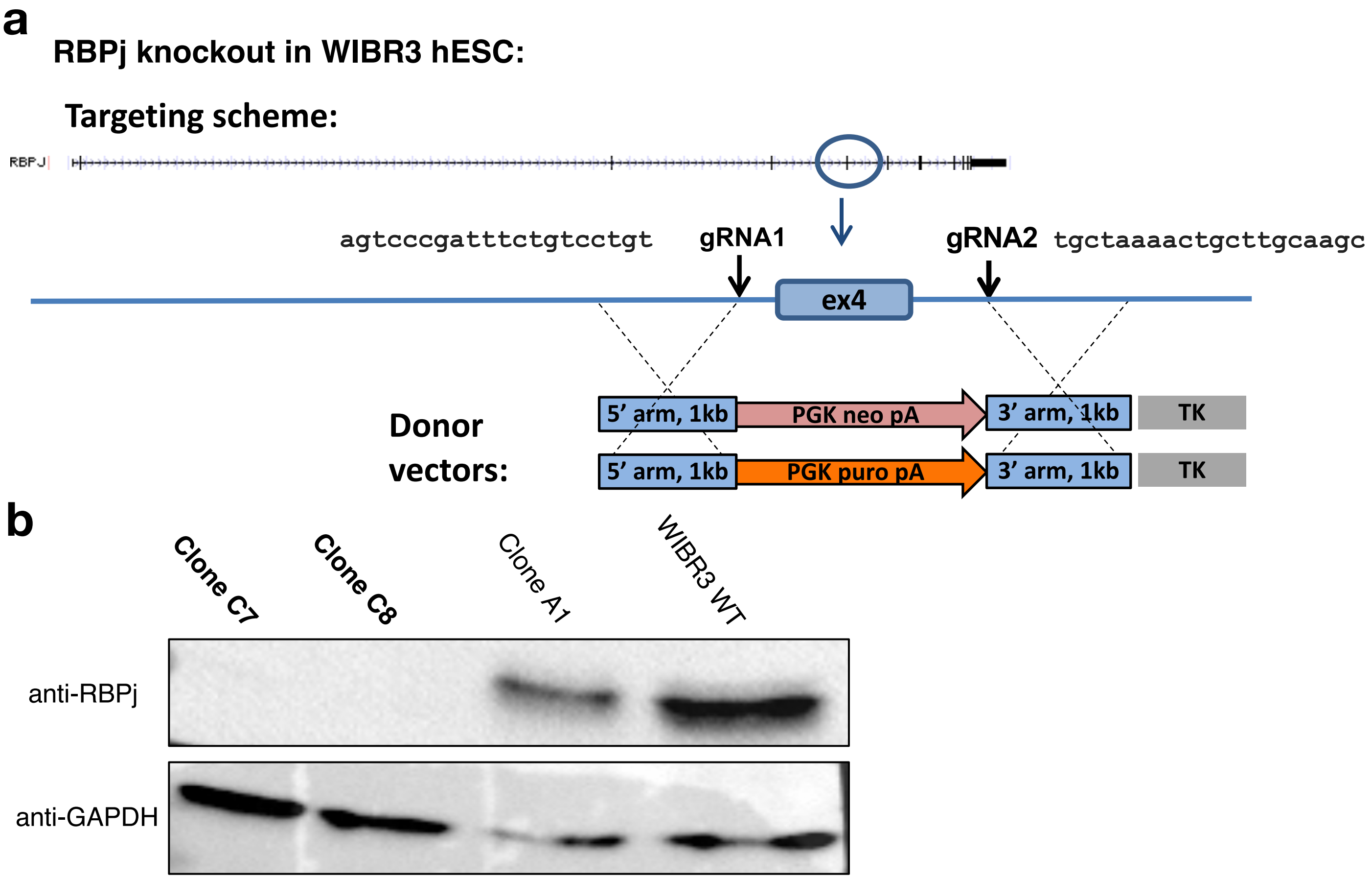
a
RBPj knockout in WIBR3 hESC:
Targeting scheme:
RBPJ
agtcccgatttctgtcctgt
gRNA1
ex4
gRNA2
tgctaaaactgcttgcaagc
Donor vectors:
5’ arm, 1kb
PGK neo pA
3’ arm, 1kb
TK
5’ arm, 1kb
PGK puro pA
3’ arm, 1kb
TK
b
Clone C7
Clone C8
Clone A1
WIBR3 WT
anti-RBPj
anti-GAPDH

Figure S34
